# Hybridizing salamanders experience accelerated diversification

**DOI:** 10.1101/760264

**Authors:** Austin H. Patton, Mark J. Margres, Brendan Epstein, Jon Eastman, Luke J. Harmon, Andrew Storfer

## Abstract

Whether hybridization generates or erodes species diversity has long been debated, but to date most studies have been conducted at small taxonomic scales. Salamanders (order Caudata) represent a taxonomic order in which hybridization plays a prevalent ecological and evolutionary role. We employed a recently developed model of trait-dependent diversification to test the hypothesis that hybridization impacts the diversification dynamics of species that are currently hybridizing. We find strong evidence supporting this hypothesis, showing that hybridizing salamander lineages have significantly greater net-diversification rates than non-hybridizing lineages. This pattern is driven by concurrently increased speciation rates and decreased extinction rates in hybridizing lineages. Our results support the hypothesis that hybridization can act as a generative force in macroevolutionary diversification.

## Introduction

A leading unresolved question in evolutionary biology is whether hybridization, defined as the interbreeding between two genetically distinct lineages^1^, acts as a creative or destructive evolutionary force^2–6^. The prevailing view in the animal literature is that hybridization constrains lineage diversification because hybrid lineages are often documented to be less fit than parentals^4,5,7^. Under this scenario, hybridization is predicted to increase extinction rates. Further, introgressive hybridization has the potential to “wash away” accumulating divergence among incompletely isolated lineages^8–10^, leading to a prediction of decreased speciation rates. In contrast, the prevailing view in the plant literature is that hybridization enhances adaptive potential by introducing novel genetic and phenotypic variation^11–13^. Reinforcement, or the accumulation of post-zygotic reproductive isolation through selection against hybrids^14^, has long been considered to expedite the speciation of diverging lineages^9^. Reinforcement has the potential to act even when heterosis occurs, as hybrids may be largely sterile, such as in *Triturus* newts^15,16^. Additionally, hybridization-mediated shuffling of old genetic variants may fuel rapid diversification, as outlined by the combinatorial view of speciation^17^. Accordingly, hybridization is predicted to increase speciation rates and/or decrease extinction rates.

Whereas evidence of hybrid speciation in plants has long been abundant, evidence for widespread hybrid speciation in animals is relatively scarce^6,10,18–21^. In allopolypoid hybrid speciation, a mode of speciation common in plants but rare in animals, nearly complete reproductive isolation may evolve in a single generation due to a change in ploidy^11^. In contrast, homoploid hybrid speciation typically has to occur in the face of continued gene flow, which acts to homogenize the diverging hybrid lineages^10^.

Interest in hybrid-mediated speciation has recently burgeoned, but studies have typically been limited in taxonomic scope. Studies of the effect of hybridization on diversification have most commonly been conducted among closely related pairs or small clades of taxa, and results have been equivocal^1,22–28^. Additionally, studies of hybridization often occur along different stages of the speciation continuum^29^, whereby hybridization can appear as a force that either facilitates or impedes speciation. Recent work highlights this uncertainty by demonstrating that the outcomes of hybridization depend on the underlying nature of selection pressures and demography^30^. We suggest that studies at broad phylogenetic and macroevolutionary timescales can help overcome these limitations by providing a phylogenetic context in which to view repeated hybridization events over evolutionary timescales and the consequent impact on lineage diversification rates.

Here, we conduct a taxonomically-broad test of the relationship of hybridization with macroevolutionary diversification rates. We study salamanders (order Caudata, ca. 716 spp as of October 2018^31^), which are particularly suitable for this study because hybridization is pervasive and has been studied extensively (i.e., nearly 1/3 of N. American species hybridize: Supplementary Fig. S1, Supplementary Table S1). Additionally, sufficient sequence data are available to resolve the phylogenetic relationships among most (~63%) taxa within this group^32^.

Salamanders present a valuable case study of hybridization, as numerous groups are comprised of speciose, yet morphologically and ecologically conservative species^23,33–38^. As a consequence of this frequent morphological and ecological conservatism, many species have come to be described on the basis of molecular differentiation (e.g.^39–44^). Many salamander groups have diversified through primarily non-ecological means of speciation (i.e.^23,33,35,45–48^). Salamander species have often diverged in allopatry/parapatry following restriction to refugial, isolated populations during periods of climatic fluctuations or via orogeny of mountain ranges (e.g.^49–51^). Following the evolution of incomplete reproductive isolation, young, diverging lineages may then come into secondary contact and hybridize (e.g.^23^). If salamanders are indeed predisposed to the evolution of incomplete reproductive isolation, then hybridization may play an important role in their diversification.

If hybridization plays a meaningful role in the diversification process, differences in diversification rates among hybridizing and non-hybridizing taxa are expected. Thus, we test the hypothesis that there is a difference in diversification rates (speciation and extinction rates) between contemporaneously hybridizing and non-hybridizing salamander lineages. Note that we are simply testing whether contemporary hybridization influences diversification rates, not whether ancient hybridization facilitated the present radiation as postulated by the hybrid swarm hypothesis^6,52^, because our experimental design cannot address this (See supplement). We replicate this test across four datasets to investigate the robustness of our results: 1) including all available data; 2) exclusion of species that do not exhibit sympatry (defined as <10% geographic range overlap) thereby lacking the opportunity to hybridize; 3) only the family Plethodontidae, which are the most widely hybridizing and diverse of the 10 salamander families; 4) all (nine) salamander families except the Plethodontidae.

## Materials and Methods

### Data Collection

We used the time-calibrated phylogeny of Amphibia^32^ as the source for downstream analyses. This tree^32^ was constructed using nine nuclear genes and three mitochondrial genes as data using RAxMLv7.2.8^53^, and time-calibrated using treePL^54^. Using the APE package (v.3.4^55^) in R, we extracted the subtree containing salamanders for subsequent analyses. This approach yielded a tree containing 469 of 716 extant species of salamanders (Supplementary Fig. S1^32^) representing approximately two-thirds of the known diversity.

Each species was scored as ‘non-hybridizable’ (NH) or ‘hybridizable’ (H), based on an extensive literature review of hybridization using the search engines Google Scholar and ISI Web of Knowledge. To do so we paired each species with the terms ‘hybrid’ and ‘introgress’ as well as dialectical and structural variants thereof (e.g., ‘hybridization’, ‘introgressed’) to search for cases of hybridization. Criteria for hybridizability under the first, “narrow” definition were the documentation of hybridization among natively distributed species, specifically the observation of heterospecific mating or hybrid offspring in the wild as detected by substantial and repeated morphological *and* molecular intermediacy. Criteria for hybridizability under the second, “broad” definition included those criteria described previously, as well as: 1) the observation of hybridization occurring in laboratory settings or among introduced and native species, and/or; 2) inference of historical introgression as determined using molecular lines of evidence (e.g. substantial and replicated genealogical discordance among molecular markers or detection via Approximate Bayesian Computation methods). This latter definition is less conservative than the former. We excluded cases where one or two individuals exhibited cyto-nuclear discordance between markers such as allozymes and mtDNA (as observed for members of *Sirenidae*^56^, and several Hynobiids^57–60^), as we did not perceive these cases to meet the criteria of substantial and replicated genealogical discordance. Although inference of hybridization via the detection of genealogical discordance warrants caution, our narrow definition of hybridization does not recognize these species as hybridizable. In total, we retrieved 56 papers (date-range: 1957-2017: Supplementary Table S1). We confirmed hybridization for roughly 11 and 13 percent of extant Caudates for the narrow and broad datasets, respectively (78 and 92 of 716 species: 17 and 20 percent of sampled taxa; Supplementary Table S1). Documented hybridization was absent from four families (Cryptobranchidae, Sirenidae, Proteidae, Rhyacotritonidae) in the narrow dataset and two families (Rhyacotritonidae, Sirenidae) in the broad dataset.

In addition to analyzing the entire salamander subclade from the Pyron^32^ amphibian phylogeny (469 total species), we compared diversification rates using three additional datasets, produced using both the narrow and broad datasets (i.e., for a total of eight datasets). In the first dataset, we required that “non-hybridizable” taxa were sympatric (>10% range overlap) with another species, thus possessing sufficient opportunity to hybridize. Consequently, species that were not sympatric with any other salamander taxa were excluded from the dataset. Therefore, species classified as non-hybridizing in this analysis may have limited opportunity to hybridize but have not been observed to do so. Although we cannot account for historical species distributions (e.g. species’ ranges may have previously overlapped), our primary (narrow) definition of hybridizability necessitates contemporary hybridization. Thus, our designation of sympatry occurs at the same time scale as our designation of hybridizability.

In the second and third datasets, we tested for a family-specific effect of hybridization on diversification rates. As plethodontids are the most diverse family of salamanders (Fig. 1B) and have been the subject of extensive study^37^, the greatest number of instances of hybridization have been documented in this group (Table S1). Additionally, many plethodontids of the genus *Plethodon* in the Eastern United States have been described as species on the basis of a threshold genetic distance (e.g..^39–42^). Coincidentally, nearly half of these species have been observed to hybridize in nature. To test for an effect of this relative taxonomic oversplitting, we excluded the *Plethodon glutinosus* group (Figure 1B) and repeated analyses under both the narrow and broad definitions of hybridization (see Supplement for additional details). Thus, we produced datasets including either only members of family Plethodontidae, or the nine families that exclude plethodontid salamanders. In total, ten analyses were conducted, using five trees for each of the narrow and broad datasets, respectively. Information on the total number of species included in each tree, as well as the number of species in each state (i.e., H or NH) may be found in Supplementary Table S1.

**Figure 1.**
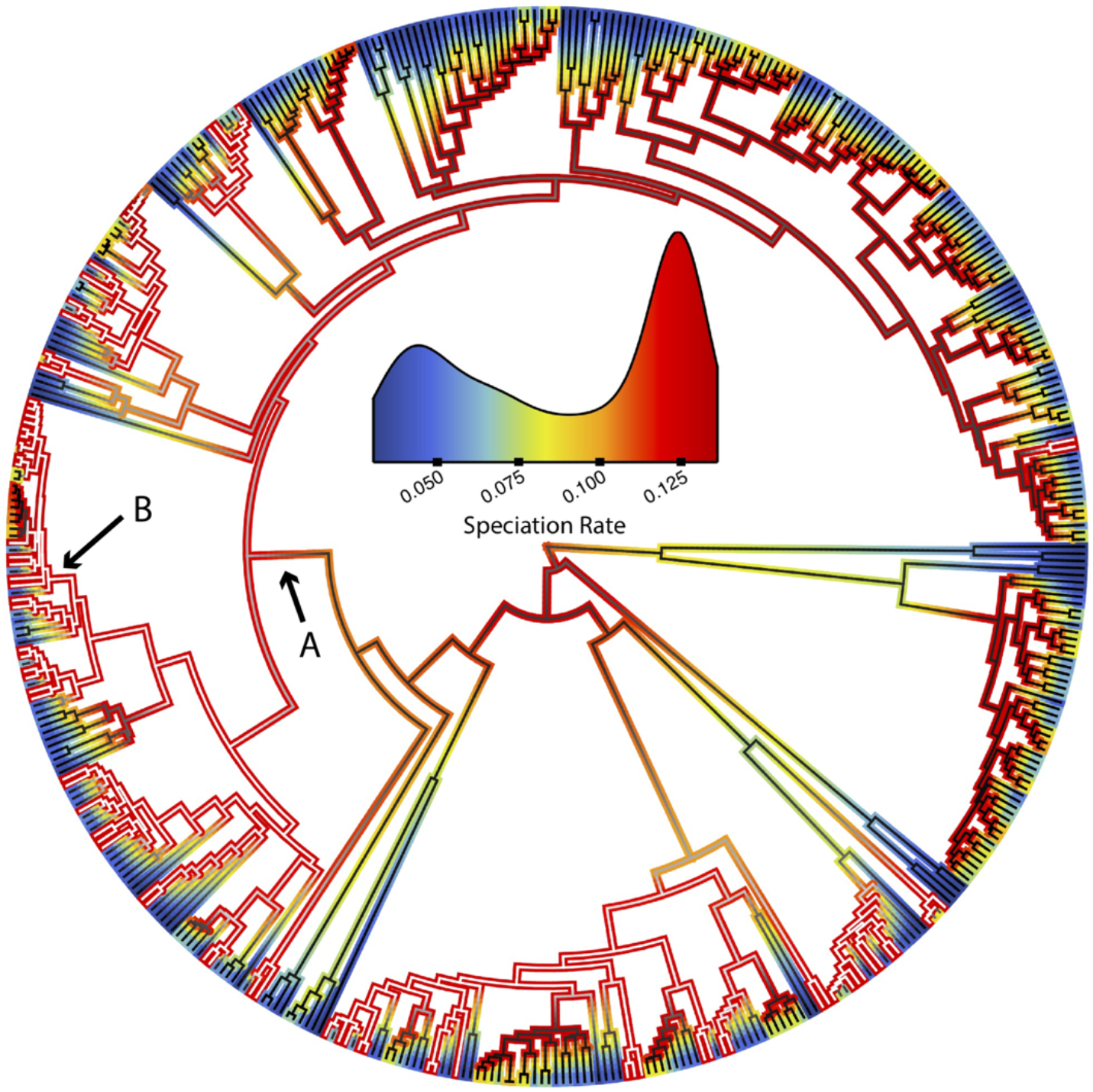
Inferred speciation rate along the salamander phylogeny for both hybridizing and non-hybridizing species. Branch outlines depict speciation rates corresponding to the inset density plot. Branch interiors depict the probability that a lineage is hybridizable (white) or non-hybridizable (black). The arrow labeled **A** denotes family Plethodontidae, and the arrow labeled **B** denotes the *Plethodon glutinosus* group.

### Assessment of Trait Dependent Diversification

We applied the HiSSE (Hidden State Speciation & Extinction^61^) trait-dependent diversification model to test for differences in speciation and extinction rates between hybridizing and non-hybridizing lineages. HiSSE infers speciation and extinction rates for a binary character while allowing for heterogeneity in diversification rate to exist within each character state. In the HiSSE model, hidden states are co-distributed with the trait of interest and account for unsampled traits that may simultaneously contribute to the diversification process. The hidden state need not be associated with any single trait but may instead be associated with a set of traits or suite of traits. Inclusion of the hidden state thus ameliorates the confounding effects of unsampled traits on diversification rate estimations by allowing for greater rate heterogeneity in the tree than in previous SSE models^61^. Thus, we are in essence measuring the impact of contemporary hybridization while controlling for other, correlated traits on diversification rate. Finally, by accounting for increased rate heterogeneity in character-dependent and character-independent (null) models through the inclusion of hidden states, model rejection properties are greatly improved^61^ relative to previous SSE models.

We evaluated a total of 14 competing models using HiSSE, half of which represent a model of character-dependent diversification and the remaining half represented models of character independence (Supplementary Tables S2-S9). Models of trait dependence varied in the number of hidden states included and in the number of free transitions among states. In character-dependent models, our four states where Hybridizing (H) and Non-hybridizing (NH), each being associated with one of the two hidden state (A & B) for four total character states/diversification regimes (H-**A**, H-**B**, NH-**A**, NH-**B**).

Additionally, to account for the fact that diversification rates may be biased by incomplete sampling of extant diversity within a phylogeny^62–64^, we assume that 20% of extant species of salamanders hybridize in nature. This value was chosen because it approximately equals the mean frequency of hybridization across our datasets (19.75). To explore the effect this assumption had on our results, we repeated these analyses assuming 1) we have sampled all extant hybridizing species (Supplementary Fig. S2), 2) our sampling of character states is proportional to their prevalence in nature (Supplementary Fig. S3), and 3) that 30% of extant species hybridize (Supplementary Fig. S4). Thus, while unable to designate all hybridizing taxa as such in our phylogeny due to their not being represented in the literature, we have explicitly addressed this uncertainty in our analysis. We elaborate upon our choice of sampling fractions in the Supplementary Materials.

To assess whether the inclusion of extremely young species impacted our diversification rate estimates, we ran a single analysis excluding species younger than 1MY, as per Beaulieu & O’Meara^61^ using the narrow dataset and all species. This in turn led to the removal of 14 species. Results were qualitatively identical to those obtained including these young species, so all subsequent analyses were conducted including them. To improve the performance of the Maximum Likelihood optimization procedure implemented in HiSSE, we used simulated annealing to first traverse the likelihood surface to identify optimal starting values for subsequent ML-optimization. Rather than reporting the results of individual model fits, we instead take the approach of investigating model-averaged parameter estimates for each sampled character state^65^ (see Supplementary Information for further justification). That is, parameter estimates obtained from each fitted model are averaged together such that their contribution to the average is proportional to their relative support (Akaike weights) among the set of candidate models (Supplementary Table S13). This leads the best supported models to have the greatest impact on the final model averaged parameter estimates. Diversification rates are returned for each sampled state respectively, as not all models include hidden states.

To test for significance among diversification rates inferred for each state, we calculated all possible ratios between non-hybridizing and hybridizing species’ model-averaged parameter estimates and calculating the proportion of comparisons in which the value for the non-hybridizing lineage is greater than that of hybridizing lineage. Thus, we obtain empirical P-values (reported in Supplementary Table S14) in which a value of 0 means in every comparison, hybridizing lineages were inferred to have rates greater than those of non-hybridizing lineages, and vice versa. This test is extremely conservative and tests the null hypothesis that non-hybridizing species always experience diversification rates greater than hybridizing species.

In summary, four phylogenies (all species, sympatric species, plethodontids, & non-plethodontids) were analyzed using two datasets (narrow & broad definitions of hybridization). Each data/tree-set combination was tested assuming the three aforementioned differences in prevalence of hybridization in nature. In all, a total of 32 rounds of model testing were performed, comparing seven models of trait-independent diversification and seven models of trait-dependent diversification for each sampling fraction. Further information on how these data/tree-sets were produced and analyzed may be found in the Supplementary Materials. Lastly, a description of our test of sensitivity to phylogenetic uncertainty may also be found in the Supplement (Supplementary Fig. S6 & S7; Supplementary Table S15)

To complement our HiSSE analyses in a manner that is largely insensitive to the potentially confounding relationship between branch lengths and propensity to hybridize (see discussion), we conducted sister clade comparisons. Specifically, using all comparisons of three or more taxa (i.e. two sister species hybridize, and the sister lineage does not), we used the method of Barraclough, Harvey and Nee^66^ to test the hypothesis that hybridizing clades had greater richness than non-hybridizing clades. To assess confidence, 1000 permutations of contrast signs were conducted. This test was repeated for each of the eight datasets described above (Supplementary Table S16).

To test for possible circularity of causality between diversification rates, species richness and opportunity to hybridize, we quantified the relationships between the three. Specifically, we tested for a relationship between 1) mean diversification rates and the proportion family hybridizing, 2) mean diversification rates and species richness, and 3) proportion family hybridizing and species richness (Supplementary Fig. S8). All analyses were conducted at the family level, and mean diversification rates were obtained using lineage-specific model-averaged diversification (speciation, extinction, net-diversification) rates. Simple linear regressions were conducted in R v3.6.1^67^.

## Results

Across both datasets (narrow and broad) assuming 20% of species hybridize, three of four analyzed phylogenies consistently found that hybridizing species experience increased speciation rates, decreased extinction rates, and therefore increased net-diversification rates (all significant: Figs. 1, 2 & 3; Tables 1, 2, Supplementary Tables S6 & S7). Net-diversification of hybridizing lineages in these trees were on average 4X greater than that of non-hybridizing lineages. Our results were insensitive to phylogenetic uncertainty (Supplementary Fig. S7; Supplementary Table S15).

**Figure 2.**
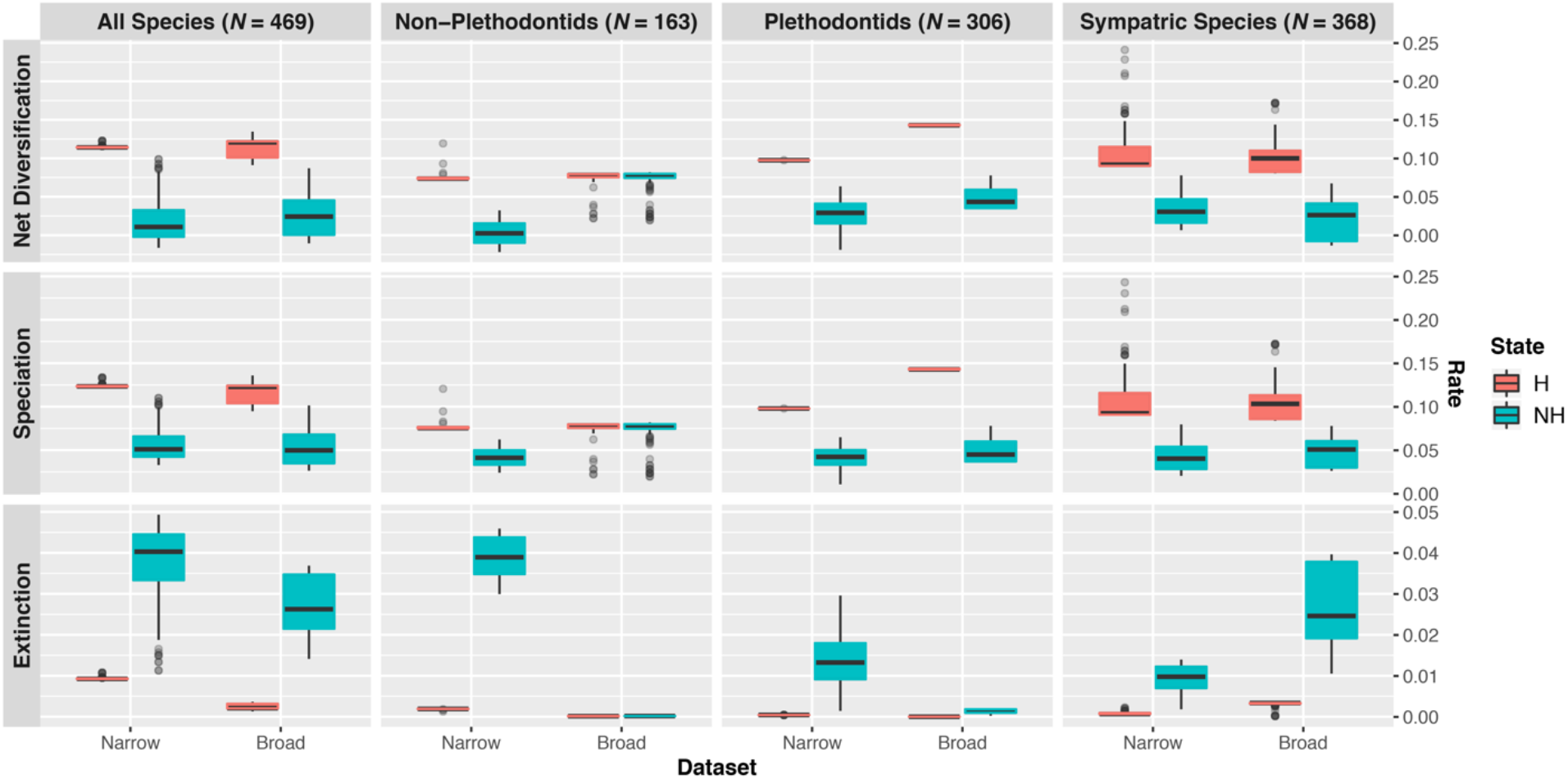
Model-averaged lineage-specific diversification rate estimates at the tips of the phylogeny assuming 20% of species hybridize. Results using different trees are displayed by column, whereas results for different parameters are displayed by row. Hybridizing lineages (H) are displayed in red, whereas non-hybridizing (NH) lineages are displayed in blue. Results for both the narrow and broad datasets are shown; the narrow dataset includes only instances of contemporary hybridization in nature among natively distributed species, whereas the broad datasets includes instances of historical introgression and non-natural hybridization.

**Table 1.** Model-averaged diversification rate estimates at the tips of the phylogeny ± 2 SE. Results assume 20% of extant species hybridize. Bold indicate parameter estimates that differ significantly among character states, with the boldened values as the larger rate estimate. Significance was determined by calculating all possible ratios between non-hybridizing and hybridizing species’ model-averaged parameter estimates and calculating the proportion of comparisons in which the value for the non-hybridizing lineage is greater than that of hybridizing lineage. This comparison thus produced an empirical P-value with which significance could be determined.

| Dataset | Tree | Speciation<br>$\lambda$ | | Extinction<br>$\mu$ | | Net Diversification<br>$r = \lambda - \mu$ | | Extinction Fraction<br>$\xi = \mu \div \lambda$ | | Turnover<br>$\tau = \lambda + \mu$ | |
| --- | --- | --- | --- | --- | --- | --- | --- | --- | --- | --- | --- |
|  |  | H | NH | H | NH | H | NH | H | NH | H | NH |
| Narrow | All Species | **0.124** ± 5.90e-4 | 0.055 ± 1.69e-3 | 9.39e-3 ± 8.62e-5 | **0.038** ± 8.36e-4 | **0.115** ± 5.04e-4 | 0.017 ± 2.54e-3 | 0.076 ± 3.20e-4 | **0.794** ± 0.036 | **0.134** ± 6.76e-4 | 0.094 ± 8.60e-4 |
| | Sympatric Species | **0.111** ± 8.140e-3 | 0.042 ± 1.73e-3 | 8.57e-4 ± 8.68e-5 | **9.45e-3** ± 3.5e-4 | **0.110** ± 8.04e-3 | 0.032 ± 2.08e-3 | 7.54e-3 ± 1.53e-4 | **0.283** ± 0.020 | **0.112** ± 8.22e-3 | 0.051 ± 1.38e-3 |
| | Plethodontids | **0.098** ± 5.50e-6 | 0.041 ± 1.48e-3 | 4.26e-4 ± 1.20e-8 | **0.014** ± 7.66e-4 | **0.098** ± 5.48e-6 | 0.028 ± 2.24e-3 | 4.35e-3 ± 1.63e-7 | **0.435** ± 0.051 | **0.098** ± 5.50e-6 | 0.055 ± 7.12e-4 |
| | Non-Plethodontids | **0.078** ± 3.38e-3 | 0.041 ± 1.73e-3 | 1.89e-3 ± 4.92e-5 | **0.039** ± 7.88e-4 | **0.076** ± 3.42e-3 | 2.2e-3 ± 2.5e-3 | 0.025 ± 1.19e-3 | **1.040** ± 0.066 | 8.00e-2 ± 3.32e-3 | 0.080 ± 9.80e-4 |
| Broad | All Species | **0.117** ± 2.72e-3 | 0.053 ± 2.04e-3 | 2.39e-3 ± 1.55e-4 | **0.027** ± 7.24e-4 | **0.115** ± 2.88e-3 | 0.026 ± 2.74e-3 | 0.021 ± 1.95e-3 | **0.647** ± 0.039 | **0.120** ± 2.58e-3 | 0.080 ± 1.37e-3 |
| | Sympatric Species | **0.106** ± 4.60e-3 | 0.047 ± 1.85e-3 | 3.02e-3 ± 1.64e-4 | **0.027** ± 1.10e-3 | **0.103** ± 4.74e-3 | 0.019 ± 2.96e-3 | 0.031 ± 2.04e-3 | **0.732** ± 0.057 | **0.109** ± 4.46e-3 | 0.074 ± 7.58e-4 |
| | Plethodontids | **0.143** ± 1.79e-4 | 0.049 ± 1.64e-3 | 3.48e-6 ± 1.33e-6 | **1.40e-3** ± 6.66e-5 | **0.143** ± 1.78e-4 | 0.048 ± 1.70e-3 | 2.42e-5 ± 9.24e-6 | **0.033** ± 2.26e-3 | **0.143** ± 1.80e-4 | 0.051 ± 1.57e-3 |
| | Non-Plethodontids | 0.071 ± 5.42e-3 | 0.072 ± 2.88e-3 | 1.44e-4 ± 1.04e-5 | **1.66e-4** ± 3.72e-7 | 0.071 ± 5.42e-3 | 0.072 ± 2.88e-3 | 2.38e-3 ± 4.80e-4 | **2.60e-3** ± 2.52e-4 | 0.071 ± 5.42e-3 | 0.072 ± 2.88e-3 |

**Table 2.**
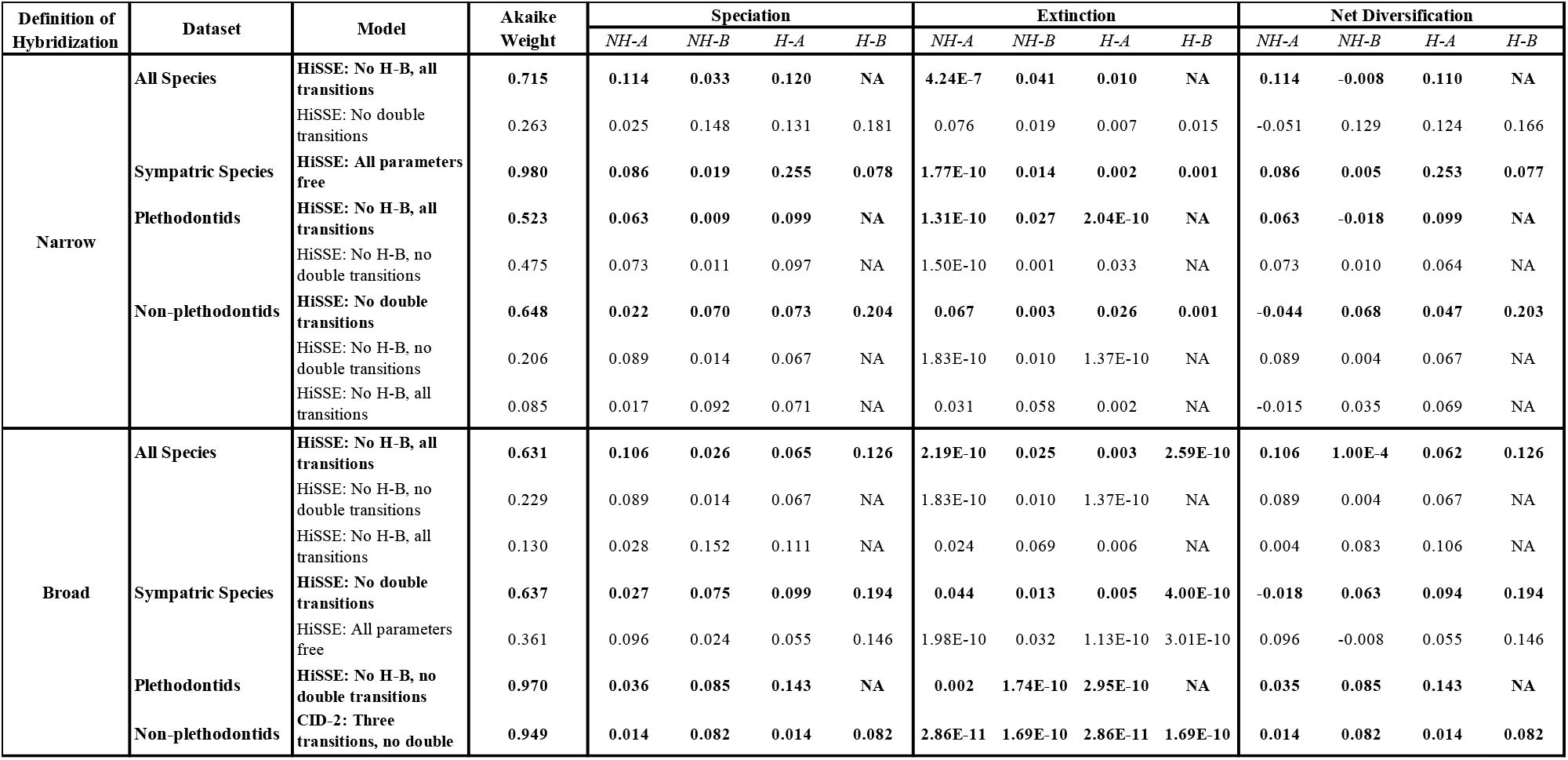
Best-fit models assuming 20% of species hybridize. Included are models that received >5% Akaike weights for their respective analyses. For each dataset, the best fit model is bold. Maximum-likelihood parameter estimates for speciation, extinction, and net diversification are reported. Non-hybridizing is abbreviated as NH, Hybridizing as H; A and B indicate the two hidden states.

In all cases of inferred trait dependent diversification, a HiSSE model was the best supported (Supplementary Tables S2-S9). Of these HiSSE models, two groups emerged: one in which hybridizing species did not harbor a second hidden state (All Species – Narrow, Plethodontids – Narrow & Broad), and one in which both hybridizing and non-hybridizing taxa had two hidden states (Table 2). The former of these (without a second hidden state for hybridizing taxa) is interpretable as meaning that there is less diversification rate heterogeneity experienced by hybridizing taxa than by non-hybridizing taxa. Interestingly, non-hybridizing taxa were sometimes inferred to have slightly negative net-diversification rates (Table 2). In the case of the best-fit model using all taxa assuming 20% of species hybridize, this leads to an expected waiting time of 128 million years before the next net-loss in diversity. These negative rates are not persistent however; examination of ancestral state reconstructions indicates that the hidden state responsible (NH-B) for these rates is distributed primarily along the tips (Supplementary Figs S9 – S12).

As the average magnitude of increase in speciation rate across all 32 analyses (95% CI: 0.0557 ± 0.0059 species/MY) is significantly greater than the average decrease in extinction rate (95% CI: 0.0145 ± 0.0037 species/MY), we conclude it is primarily differential speciation that is driving the increase in net-diversification in hybridizing lineages. Interpreted as a waiting time, this means that hybridizing species, on average, speciate every 9.2 (95% CI: 8.84 – 9.62) million years and go extinct every 160.7 (118.97 – 247.96) million years.

Analysis of the tree containing all species recovered strong signal of trait dependent diversification, with significant differences between hybridization or non-hybridizing lineages (as measured by empirical P-values) found between states for all five parameter estimates (Figs. 2 & 3; Tables 1, 2, Supplementary Tables S6 – S7), regardless of whether the broad or narrow criterion was used. Speciation (λ), net-diversification (*r* = λ – μ), and turnover rate (τ = λ + μ) were greater in hybridizing than non-hybridizing lineages, whereas extinction rate (μ) and extinction fraction (ε = μ ÷ λ) were lower. Increases were on average 125% (95% CI: 117 – 133%) for speciation, 576% (486 – 699%) for net diversification and 43% (41 – 45%) for turnover. In contrast, decreases were on average 305% (292 – 317%) for extinction and 945% (893 – 997%) for extinction fraction (Fig. 2). Additional analyses that assume different percentages of extant species hybridize produced qualitatively similar results and are discussed in the supplement (Supplementary Figs. S2-S4, Supplementary Tables S2-14).

**Figure 3.**
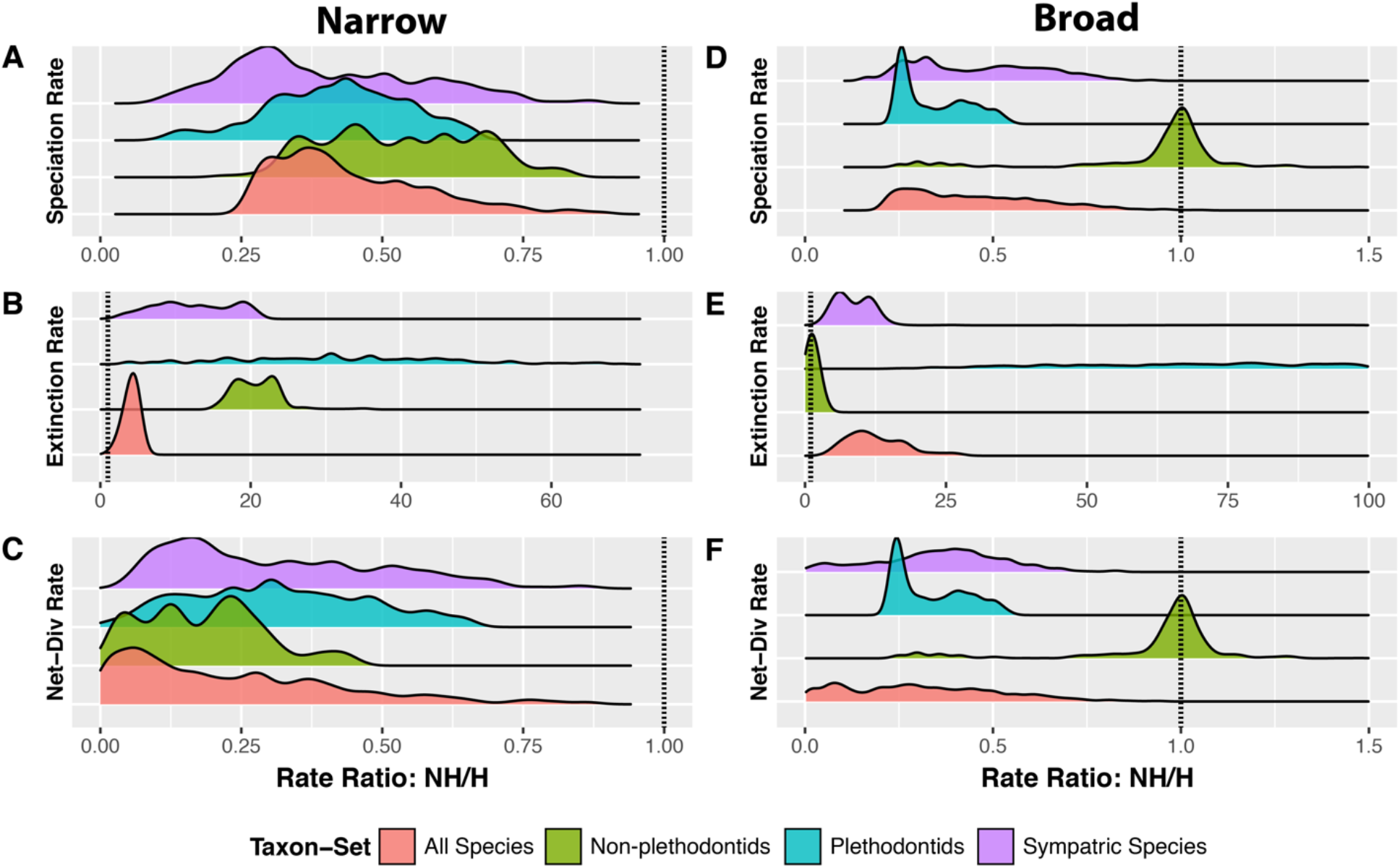
Comparison of model averaged parameter estimates among character states. Results reported here are those assuming 20% of species hybridize. A, B and C illustrate the distributions of non-hybridizing to hybridizing lineages diversification rates as estimated at the tips of the phylogeny using the narrow dataset, whereas D, E and F are those using the broad dataset. A value > 1 corresponds to a comparison in which non-hybridizing lineages experience rates greater than those of hybridizing lineages and vice-versa. Dotted vertical lines are placed at 1, at which rates are equal among states.

Analysis of the tree including only sympatric taxa regardless of the hybridization criterion (narrow versus broad) datasets recovered patterns identical to those obtained using the complete phylogeny (Figs. 2 & 3; Tables 1, 2, Supplementary Tables S6-S7). Speciation rate, net-diversification rate, and turnover increased by 164% (135 – 196%; λ), 244% (199 – 295%; *r*), and 120% (98 – 142%; τ) respectively in hybridizing versus non-hybridizing lineages. In contrast, extinction rate and the extinction fraction decreased in hybridizing lineages by 1003% (864 – 1172%; μ) and 3653% (3319 – 4002%; ε) respectively.

Similarly, plethodontids exhibited the same patterns described above (Figs. 2 & 3; Tables 1, 2, Supplementary Tables S6-S7). Hybridizing lineages experienced rates of speciation, net-diversification and turnover that were on average 139% (130 – 148%; λ), 250% (224 – 281%; *r*), and 78% (76 – 81%; τ) greater than those experienced by non-hybridizing lineages. Extinction rate and extinction fraction were reduced by 3186% (3007 – 3366%; μ) and 9000% (8727 – 11073%; ε) in hybridizing lineages relative to non-hybridizing lineages.

In contrast, analysis of non-plethodontids revealed greater ambiguity as to the impact of hybridization on diversification rates. Whereas the narrow dataset inferred trait-dependent diversification in which hybridization drove increased diversification, the broad dataset did not (Figs. 2 & 3; Tables 1, 2, Supplementary Tables S6-S7). Using the narrow dataset, all rates except turnover were found to differ significantly (Supplementary Tables S6-S7, S13-S14). Speciation rate and net diversification were 90% (75 – 107%; λ) and 3355% (1444 – 26573%; *r*) greater in hybridizing lineages, whereas extinction rate and extinction fraction were reduced by 1964% (1871 – 2062%; μ) and 4060% (3619 – 4545%; ε) in hybridizing lineages relative to non-hybridizing lineages. Analysis of the broad dataset revealed no significant differences and parameter estimates between hybridizing and non-hybridizing species.

Sister clade contrasts broadly supported results of the HiSSE analysis. That is, hybridizing clades were found to have significantly greater species richness than non-hybridizing clades for all datasets except for plethodontid salamanders. Details on significance of these tests may be found in Supplemental Table S16.

Rapidly diversifying species may have greater opportunity to hybridize due to an associated increase in species richness. We did not recover evidence supporting this interpretation. Both mean diversification rate (Adjusted *R^2^*: λ = −0.095; μ = −0.084; *r* = −0.092) and proportion of family hybridizing (Adjusted *R^2^* = −0.112) are decoupled from family species richness, despite a significant correlation (*P*: λ = 0.0002; μ = 0.0004; *r* = 0.0002) between mean diversification rate and proportion family hybridizing (Adjusted *R^2^*: λ = 0.822; μ = 0.786; *r* = 0.811; Supplementary Fig. S8).

## Discussion

Here, we show strong support that contemporary hybridization is correlated with elevated diversification rates in the order Caudata. Net-diversification of hybridizing species tends to be significantly greater than that of non-hybridizing species, driven primarily by a coincident increase in speciation and decrease in extinction rates (Figs. 2, 3; Tables 1, 2). The accelerated diversification of hybridizing salamanders appears ephemeral, however; rate differences rapidly become less pronounced deeper in the tree due to turnover of the hidden states (Fig. 1, Supplementary Figs. S9-S12). Possible mechanisms leading to this result include frequent range expansions and contractions (i.e.^68^) that have been documented in salamanders (e.g.^69–71^) and the process of reinforcement which has long been recognized to contribute to the diversification process^14,72–74^. We outline the potential contribution of each below.

Salamanders often exhibit substantial genetic differentiation at small geographic distances (e.g., 200m^75^) owing to limited dispersal abilities and low rates of gene flow^76^, thus leading to an abundance of opportunities to evolve in allopatry. Additionally, terrestrial species such as the *Plethodontid* salamanders of the southeastern United States experience elevational range expansions and contractions associated with climatic change^71^. Perhaps this combination of the primarily sessile nature of many salamander species and frequent repeated secondary contact leads to hybridization occurring regularly across evolutionary timescales. Under these scenarios, hybridization may then play a creative evolutionary role in the diversification process similar to that observed in haplochromine cichlids^24,25,77^. Allopatric speciation of haplochromine cichlids has occurred in lakes that frequently have dried, split, and reformed, whereas sympatric speciation has occurred within lakes in which lineages exhibited extreme habitat specificity and have been reproductive isolated at fine spatial scales^78^. Under these circumstances, hybridization may have afforded genetic rescue from the consequences of small population size by providing increased standing genetic variation and thus expedited adaptation to novel stressors, as in Lake Victorian cichlids post-colonization^25^.

Such hypotheses of repeated contractions, expansions, and secondary contact of salamander populations have been well supported, across both North^79–81^ and South America^50,82^, as well as in Europe^83–85^. Often associated with glaciation/deglaciation or orogeny of mountain ranges, geological events may act as species pumps for salamanders (i.e.^86^). However, while periodic geographic range expansion and contraction may initiate speciation, the reproductive isolation that evolves may be incomplete, predisposing the young species/diverging lineages to hybridization (i.e.^23^). As a result, hybridization may commonly occur in salamanders during periods of climatic fluctuations.

There is now clear evidence for latitudinal and elevational range shifts mediated by climate change^87–91^ and a consequent increase in frequency of hybridization among previously isolated taxa^92–95^. An informed understanding of the influence of hybridization on macroevolutionary diversification may thus provide invaluable context for contemporary processes. This possibility of climate-change mediated hybridization has already been demonstrated in plethodontid salamanders (*P. shermani* & *P. teyahalee*^71^), as well as in ecologically divergent subspecies of salamandrid salamanders (*S. salamandra*^96^). Thus, it seems likely that salamander species worldwide, particularly those found at high elevations due to their more limited potential geographic distributions, may experience a heightened frequency of hybridization as climate change advances. While generalizations regarding the outcome of hybridization should be made with caution^30^, our study indicates that perhaps speciation reversal^97–99^ need not be the expectation. Rather, our study implies that hybridization may facilitate adaptation to novel conditions under climate change, leading to diversification of new salamander lineages.

Here, we show the novel result of a strong correlation of contemporary hybridization with elevated speciation and net diversification at a large taxonomic scale. However, reinforcement, defined as the strengthening of prezygotic reproductive isolation in sympatry^14^, is intrinsically intertwined with hybridization. Reinforcement has been documented both experimentally^72^ and observationally^74^ to accelerate the initiation and/or completion of the speciation process^73,100,101^. For instance, reinforcement is likely to play an important role in the speciation process due to strong interspecific sexual selection and mate choice in plethodontids^102^. Indeed, patterns of sexual isolation among populations of *Plethodon jordani* and *P. teyahalee* match expectations of reinforcement^14^, with sexual selection being stronger in sympatry than in allopatry^103^. Although we cannot currently quantify the contribution of reinforcement to diversification rate differences using our data, we urge further research measuring the degree of association between contemporaneous hybridization and reinforcement among taxa. Nonetheless, were reinforcement to play a role in the production of the patterns observed in this study, the very occurrence/process of hybridization would be the ultimate driver (i.e., cannot have reinforcement without hybridization). Under such a scenario, our study design is well-suited to identify such a signal.

Although a generative role of hybridization is robustly supported across three of our four datasets, evidence for such a role outside of the Plethodontidae is more limited (Table 1, Supplementary Tables S2-S14). We find two possible explanations for this finding. Firstly, the positive association between diversification rates and hybridization may be unique to Plethodontid salamanders. However, family Plethodontidae is the largest extant family of salamanders, comprising approximately 2/3rds of the present diversity (471 of 716 species: amphibiaweb.org). Thus, our observation of hybridization facilitating the diversification process applies to the majority of salamanders and implies that, at a minimum, contemporaneous hybridization does not impede the diversification process of extant salamanders.

Secondly, it is highly probable that our analysis of non-plethodontid salamanders is lacking in power. SSE models have long been known to lose much of their power when dealing with small number of OTUs (trees < 300 taxa^104^). For example, for trees of 300 species, BiSSE attains a power of at most 50%, with power dropping below 15% of trees of 100 taxa^104^. Our phylogeny of non-plethodontids includes only 167 species; that we detected a positive relationship between hybridization and speciation rates using our narrow (most conservative) dataset despite such reduced power is a testament to the strength of the signal in our data. Whereas our larger datasets [complete (469 spp), sympatric (368 spp) and plethodontids (306 spp)] have greater power, our lack of detection of a relationship between hybridization and diversification rates in non-plethodontids using our broad definition of hybridization is perhaps unsurprising, given the low power of the analysis (also see Supplementary Materials for an elaboration of power). Although the power of HiSSE under such scenarios has not been specifically established, accuracy of parameter estimation does decay with decreasing tree size^61^. Consequently, we cautiously interpret the results of the analysis of non-plethodontids (163 spp). Interestingly, sister clade comparisons consistently supported a positive relationship between hybridization and species richness in non-plethodontids, despite not supporting such a relationship in plethodontids (Supplementary Table S16). These results are insensitive to branch-lengths, thereby ameliorating potential concerns related to the relationship between hybridization, branch-lengths, and diversification rates^23^.

Importantly, parameter estimates are largely reasonable. For instance, the greatest speciation rate inferred by any analysis (Plethodontids assuming 30% of taxa hybridize using the full tree: Supplementary Table S13), of 0.159 species/million years (MY) can be interpreted as a waiting time, such that on average, hybridizing species speciate every 6.29 MY. Extinction rates appear less reliably estimated however; some estimates functionally equal zero, leading to the large percent decrease in extinction rates observed for hybridizing relative to non-hybridizing species. In some cases, extinction rates in non-hybridizing taxa leads to negative net-diversification rates. That being said, averaged extinction rates across all analyses for hybridizing and non-hybridizing taxa led to more reasonable waiting times of 127.9 and 54.2 MY respectively. Further, recent studies have documented even more negative net-diversification rates than inferred herein^105^. Taken together, it appears that extinction plays an important role in the diversification of salamanders, leading to a reduction in net-diversification rates towards the present relative to hybridizing species.

An important question regarding the interpretation of our results is the relationship between lineage diversification rates, species richness, and opportunity to hybridize. Because the relationship between lineage diversification rate and opportunity to hybridize are not necessarily independent, rapidly diversifying lineages may simply have greater opportunity to hybridize due to increased diversification rates. Although a legitimate concern, we did not find evidence that the increased diversification rates we observe are due to increased family-level species richness leading to increased opportunity to hybridize (Supplementary Fig. S8). Further, it is unlikely that non-random taxonomic sampling has biased our results, as there is no relationship between clade specific sampling fraction and frequency of hybridization (Supplementary Fig. S5).

Although the ability of methods to accurately infer extinction rates has been debated recently^63,106,107^, we emphasize that our results are robust to this concern. Our central result, that hybridizing lineages experience increased net diversification, is driven by both increased speciation rates and decreased extinction rates. Further, in nearly all cases, the magnitude of increase of speciation rate is greater than that of the decrease in extinction. Thus, our results are likely robust even to inaccuracies in the estimation of extinction rate.

An important distinction between our study and most previous studies investigating the influence that hybridization exerts on the diversification process is that of the time-scale at which hybridization is being assessed. Following Seehausen’s^6^ landmark paper “Hybridization and Adaptive Radiation,” tests and discussion of his hypothesis, that ancient, widespread hybridization facilitates adaptive radiation became abundant in the literature (e.g.^18,22,23,108,109^). Whereas much of the subsequent studies focused on *ancient* hybridization, our study instead focuses on the effects of *contemporary* hybridization.

In a pertinent study, Wiens et al.^23^, tested the hybrid swarm hypothesis in the *Plethodon glutinosus* group (indicated in Fig. 1B) using two nuclear and two mitochondrial genes. They did not recover strongly supported evidence of genealogical discordance at the base of this group; these results were interpreted as not being supportive of Seehausen’s hypothesis. Further, they identified a positive relationship between age of species and reproductive isolation. They argue that the observed relationship between diversification rate and hybridization in this group was a consequence of this relationship. Although a legitimate concern, we argue that this hybridization is likely to still have biologically relevant consequences on diversification rates. Specifically, hybridization may either 1) facilitate the divergence of these young species i.e. through reinforcement/strengthening of prezygotic isolation, or 2) erode their divergence leading to species collapse. Whereas the former hypothesis predicts increased speciation rates, the latter predicts increased extinction rates. We find strong, consistent evidence in favor of the former.

We explicitly tested the hypothesis that contemporary hybridization plays a creative role in the diversification process in the broadest taxonomic and temporal scale study to date, and our observations strongly supported the predictions of this hypothesis. Specifically, hybridization was found to be correlated with both increased speciation rates and decreased extinction rates, resulting in increased net diversification rates relative to non-hybridizing lineages. Although other factors certainly contribute to the observed diversification dynamics, we have shown that hybridization plays a significant role, while accounting for hidden, correlated states in our analysis. Nearly all studies of hybridization have focused on individual case studies in which hybridization results in species collapse^98^ or promotes diversification in a single species group^12,13,22,27^. Such studies are necessarily limited in the extent to which their results may be generalized^30^, particularly because results were equivocal across studies. Consequently, we advocate that our approach can be applied at broad taxonomic and evolutionary timescales to facilitate robust tests of the role of hybridization in the lineage diversification process. We anticipate our results are broadly generalizable to animal groups in which homoploid hybridization occurs because only 17 species of salamanders are known to be polyploid^110^, and our dataset includes only seven (*Ambystoma mexicanum, A. barbouri, A. jeffersonium, A. laterale, A. texanum, A. tigrinum*, and *Lissotriton vulgaris*) hybridizing polyploid taxa (none of which are plethodontids). Our study adds to the growing evidence that hybridization may fuel rapid diversification (e.g.^52^) and is a compliment to speciation genomics studies characterizing the genomic basis of this process (e.g.^17,111^). Herein we have shown that hybridization may act as a generative force across a phylogenetic order, and additional studies at such macroevolutionary scales are needed to determine if this pattern holds more generally across the tree of life.

## Acknowledgements

We thank Adam Leaché, David Weisrock & Maria Servedio for comments on earlier versions of the manuscript. We also thank Daniel Caetano, Josef Uyeda, Rosana Zenil-Ferguson and Jeremy Beaulieu for insightful discussions regarding the study and for technical advice. The authors have no conflicts of interest to declare.

## Funding

This work was supported by NSF DEB-1316549 AS, NSF DEB 1208912 to LJH, and the WSU School of Biological Sciences Elling Endowment Fund to AHP.

## Author Contributions

A.H.P. and J.E. conceived of and designed the study, A.H.P. conducted all analyses, and wrote the manuscript. A.H.P, M.J.M, B.E., J.E., L.J.H, and A.S. contributed to revisions of the manuscript. A.H.P., J.E., & M.J.M. conducted the literature review and B.E. contributed to early analyses.

## Notes

#### Summary of Updates

Revised text and inclusion of additional analysis following review.

## References

1. Barton, N. H. & Hewitt, G. M. Analysis of hybrid zones. Annu. Rev. Ecol. Syst. 16, 113–148 (1985).

2. Darwin, C. On the Origin of Species by Means of Natural Selection. (Murray. (1859).

3. Fisher, R. The Genetical Theory of Natural Selection. (Clarendon Press. (1930).

4. Mayr, E. Animal species and evolution. (Harvard University Press. (1963).

5. Futuyma, D. J. On the role of species in anagenesis. Am. Nat. 130, 465–473 (1987).

6. Seehausen, O. Hybridization and adaptive radiation. Trends Ecol. Evol. 19, 198–207 (2004).

7. Grant, P. R. & Grant, B. R. Hybridization of bird species. Science (80-.). 256, 193–197 (1992).

8. Seehausen, O., Van Alphen, J. J. & Witte, F. Cichlid fish diversity threatened by eutrophication that curbs sexual selection. Science (80-.). 277, 1808–1811 (1997).

9. Turelli, M., Barton, N. H. & Coyne, J. A. Theory and speciation. Trends Ecol. Evol. 16, 330–343 (2001).

10. Buerkle, C. A., Morris, R. J., Asmussen, M. A. & Rieseberg, L. H. The likelihood of homoploid hybrid speciation. Heredity (Edinb). 84, 441–451 (2000).

11. Dowling, T. E. & Secor, C. L. the Role of Hybridization and Introgression in the Diversification of Animals. Annu. Rev. Ecol. Syst. 28, 593–619 (1997).

12. Rieseberg, L. H., Archer, M. A. & Wayne, R. K. Transgressive segregation, adaptation and speciation. Heredity (Edinb). 83, 363 (1999).

13. Soltis, P. S. & Soltis, D. E. The role of hybridization in plant speciation. Annu. Rev. Plant Biol. 60, 561–588 (2009).

14. Coyne, J. A. & Orr, H. A. Speciation. (Sinauer Associates Sunderland. (MA, 2004).

15. Francillon-Vieillot, H., Arntzen, J. W. & Geraudie, J. Age, Growth and Longevity of Sympatric Triturus cristatus, T. marmoratus and Their Hybrids (Amphibia, Urodela): A Skeletochronological Comparison. J. Herpetol. 24, 13 (1990).

16. Arntzen, J. W., Jehle, R., Bardakci, F., Burke, T. & Wallis, G. P. Asymmetric Viability of Reciprocal-Cross Hybrids Between Crested and Marbled News (Triturus cristatus and T. marmoratus). Evolution (N. Y). 63, 1191–1202 (2009).

17. Marques, D. A., Meier, J. I. & Seehausen, O. A combinatorial view on speciation and adaptive radiation. (Trends in ecology & evolution, 2019).

18. Mallet, J. Hybrid speciation. Nature 446, 279–83 (2007).

19. Mavárez, J. & Linares, M. Homoploid hybrid speciation in animals. Mol. Ecol. 17, 4181–4185 (2008).

20. Gross, B. & Rieseberg, L. H. The ecological genetics of homoploid hybrid speciation. J. Hered. 96, 241–252 (2005).

21. Schumer, M., Rosenthal, G. G. & Andolfatto, P. How common is homoploid hybrid speciation? Evolution (N. Y). 68, 1553–1560 (2014).

22. Mavárez, J. et al. Speciation by hybridization in Heliconius butterflies. Nature 441, 868–871 (2006).

23. Wiens, J. J., Engstrom, T. N. & Chippindale, P. T. Rapid diversification, incomplete isolation, and the “speciation clock” in North {A}merican salamanders (genus Plethodon): testing the hybrid swarm hypothesis of rapid radiation. Evolution (N. Y). 60, 2585–2603 (2006).

24. Keller, I. et al. Population genomic signatures of divergent adaptation, gene flow and hybrid speciation in the rapid radiation of Lake Victoria cichlid fishes. Mol. Ecol. 22, 2848–2863 (2013).

25. Meier, J. I. et al. Ancient hybridization fuels rapid cichlid fish adaptive radiations. Nat. Commun. 8, 14363 (2017).

26. Grant, P. R. & Grant, B. R. No Title. (Darwin’s finches on Daphne Major island. (Princeton University Press, 2014).

27. MacLeod, A. et al. Hybridization masks speciation in the evolutionary history of the Galápagos marine iguana. Proc. R. Soc. B 282, 20150 (2015).

28. Yakimowski, S. B. & Rieseberg, L. H. The role of homoploid hybridization in evolution: a century of studies synthesizing genetics and ecology. Am. J. Bot. 101, 1247–1258 (2014).

29. Seehausen, O. & others. Genomics and the origin of species. Nat. Rev. Genet. 15, 176–192 (2014).

30. Gompert, Z. & Buerkle, C. A. What, if anything, are hybrids: enduring truths and challenges associated with population structure and gene flow. Evol. Appl. 9, 909–923 (2016).

31. Berkeley, U. of C. Species by the numbers. (2018). at <http://www.amphibiaweb.org/amphibian/speciesnums.html>

32. Pyron, R. A. Biogeographic analysis reveals ancient continental vicariance and recent oceanic dispersal in amphibians. Syst. Biol. 63, 779–797 (2014).

33. Wiens, J. J. Speciation and Ecology Revisited: Phylogenetic Niche Conservatism and the Origin of Species. Evolution (N. Y). 58, 193–197 (2004).

34. Kozak, K. H., Weisrock, D. W. & Larson, A. Rapid lineage accumulation in a non-adaptive radiation: Phylogenetic analysis of diversification rates in eastern North American woodland salamanders (Plethodontidae: Plethodon). Proc. R. Soc. B Biol. Sci. 273, 539–546 (2006).

35. Wake, D. B. Problems with Species: Patterns and Processes of Species Formation in Salamanders. Ann. Missouri Bot. Gard. 93, 8–23 (2006).

36. Vences, M. & Wake, D. B. in Amphibian Biology, Vol. 6, Systematics (eds. Heatwole, H. H. & Tyler, M.) 2613–2669 (Surrey Beatty & Sons, Chipping Norton, Australia, 2007).

37. Wake, D. B. What Salamanders have Taught Us about Evolution. (2009). doi:10.1146/annurev.ecolsys.39.110707.173552

38. Blankers, T., Adams, D. C. & Wiens, J. J. Ecological radiation with limited morphological diversification in salamanders. J. Evol. Biol. 25, 634–646 (2012).

39. Highton, R. Speciation in eastern North {A}merican salamanders of the genus Plethodon. Annu. Rev. Ecol. Syst. 26, 579–600 (1995).

40. Highton, R. Geographic protein variation and speciation in the Plethodon dorsalis complex. Herpetologica 1997, 345–356 (1997).

41. Highton, R. Geographic protein variation and speciation in the salamanders of the Plethodon cinereus group with the description of two new species. Herpetologica 1999, 43–90 (1999).

42. Highton, R. & Peabody, R. in The Biology of Plethodontid Salamanders. (eds. Bruce, R. C., Jaeger, R. G. & Houck, L. D.) 31–93 (Kluwer Academic/Plenum Publishers, 2000).

43. Wake, D. B. & Jockusch, E. L. in The Biology of Plethodontid Salamanders 95–119 (Springer US, 2000). doi:10.1007/978-1-4615-4255-1_4

44. Patton, A. et al. A New Green Salamander in the Southern Appalachians: Evolutionary History of Aneides aeneus and Implications for Management and Conservation with the Description of a Cryptic Microendemic Species. Copeia 107, (2019).

45. Rundell, R. J. & Price, T. D. Adaptive radiation, nonadaptive radiation, ecological speciation and nonecological speciation. Trends Ecol. Evol. 24, 394–399 (2009).

46. Czekanski-Moir, J. E. & Rundell, R. J. The ecology of nonecological speciation and nonadaptive radiations. Trends Ecol. Evol. (2019).

47. Garcia-Paris, M., Good, D. A., Parra-Olea, G. & Wake, D. B. Biodiversity of Costa Rican salamanders: Implications of high levels of genetic differentiation and phylogeographic structure for species formation. Proc. Natl. Acad. Sci. 97, 1640–1647 (2000).

48. Parra-Olea, G. & Wake, D. B. Extreme morphological and ecological homoplasy in tropical salamanders. Proc. Natl. Acad. Sci. U. S. A. 98, 7888–7891 (2001).

49. Vieites, D. R., Min, M. S. & Wake, D. B. Rapid diversification and dispersal during periods of global warming by plethodontid salamanders. Proc. Natl. Acad. Sci. U. S. A. 104, 19903–19907 (2007).

50. Parra-Olea, G., Windfield, J. C., Velo-Antón, G. & Zamudio, K. R. Isolation in habitat refugia promotes rapid diversification in a montane tropical salamander. J. Biogeogr. 39, 353–370 (2012).

51. Bryson, R. W. et al. Phylogenomic insights into the diversification of salamanders in the Isthmura bellii group across the Mexican highlands México View project Phylogeography and contact zones in Ctenosaura pectinata View project Phylogenomic insights into the diversification of salamanders in the Isthmura bellii group across the Mexican highlands. Artic. Mol. Phylogenetics Evol. (2018). doi:10.1016/j.ympev.2018.03.024

52. Gillespie, R. G. et al. Comparing Adaptive Radiations Across Space, Time, and Taxa. J. Hered. 89154, 14853 (2020).

53. Stamatakis, A. RAxML-VI-HPC: maximum likelihood-based phylogenetic analyses with thousands of taxa and mixed models. Bioinformatics 22, 2688–2690 (2006).

54. Smith, S. A. & O’meara, B. C. treePL: divergence time estimation using penalized likelihood for large phylogenies. Bioinformatics 28, 2689–2690 (2012).

55. Paradis, E., Claude, J. & Strimmer, K. APE: analyses of phylogenetics and evolution in {R} language. Bioinformatics 20, 289–290 (2004).

56. Liu, F. G. R., Moler, P. E. & Miyamoto, M. M. Phylogeography of the salamander genus Pseudobranchus in the southeastern United States. Mol. Phylogenet. Evol. 39, 149–159 (2006).

57. Fu, J. & Zeng, X. How many species are in the genus Batrachuperus? A phylogeographical analysis of the stream salamanders (family Hynobiidae) from southwestern China. Mol. Ecol. 17, 1469–1488 (2008).

58. Yoshikawa, N., Matsui, M., Nishikawa, K., Misawa, Y. & Tanabe, S. Allozymic Variation in the Japanese Clawed Salamander, Onychodactylus japonicus (Amphibia: Caudata: Hynobiidae), with Special Reference to the Presence of Two Sympatric Genetic Types. Zoolog. Sci. 27, 33–40 (2010).

59. Yoshikawa, N., Matsui, M. & Nishikawa, K. Genetic Structure and Cryptic Diversity of Onychodactylus japonicus (Amphibia, Caudata, Hynobiidae) in Northeastern Honshu, Japan, as Revealed by Allozymic Analysis. Zoolog. Sci. 29, 229–237 (2012).

60. Yoshikawa, N. & Matsui, M. Two new Salamanders of the genus Onychodactylus from Eastern Honshu, Japan (Amphibia, Caudata, Hynobiidae). (2014). doi: 10.11646/zootaxa.3866.1.3

61. Beaulieu, J. M. & O’Meara, B. C. Detecting hidden diversification shifts in models of trait dependent speciation and extinction. Syst. Biol. 65, 583–601 (2016).

62. Stadler, T. How can we improve accuracy of macroevolutionary rate estimates? Syst. Biol. 62, 321–329 (2013).

63. Beaulieu, J. M. & O’Meara, B. C. Extinction can be estimated from moderately sized molecular phylogenies. 69, 1036–1043 (2015).

64. Rabosky, D. L. & Goldberg, E. E. Model inadequacy and mistaken inferences of trait-dependent speciation. Syst. Biol. 64, 340–55 (2015).

65. Caetano, D. S., O’Meara, B. C. & Beaulieu, J. M. Hidden state models improve state-dependent diversification approaches, including biogeographical models. Evolution (N. Y). (2018).

66. Barraclough, T. G., Harvey, P. H. & Nee, S. Rate of rbcL gene sequence evolution and species diversification in flowering plants (angiosperms). Proceedings of the Royal Society of London. Ser. B. Biol. Sci. 263, 589–591 (1996).

67. R Core Team. R: A Language and Environment for Statistical Computing. (2018). at <https://www.r-project.org/>

68. Hewitt, G. M. Some genetic consequences of ice ages, and their role in divergence and speciation. Biol. J. Linn. Soc. 58, 247–276 (1996).

69. Kozak, K. H., Blaine, R. A. & Larson, A. Gene lineages and eastern North {A}merican palaeodrainage basins: phylogeography and speciation in salamanders of the Eurycea bislineata species complex. Mol. Ecol. 15, 191–207 (2006).

70. Shepard, D. B. & Burbrink, F. T. Phylogeographic and demographic effects of Pleistocene climatic fluctuations in a montane salamander, Plethodon fourchensis. Mol. Ecol. 18, 2243–2262 (2009).

71. Walls, S. C. The role of climate in the dynamics of a hybrid zone in Appalachian salamanders. Glob. Chang. Biol. 15, 1903–1910 (2009).

72. Rice, W. R. & Hostert, E. E. Laboratory experiments on speciation: what have we learned in 40 years? 47, 1637–1653 (1993).

73. Servedio, M. R. & Noor, M. A. The role of reinforcement in speciation: theory and data. Annu. Rev. Ecol. Evol. Syst. 34, 339–364 (2003).

74. Matute, D. R. & Ortiz-Barrientos, D. Speciation: The strength of natural selection driving reinforcement. Curr. Biol. 24, 955–957 (2014).

75. Zamudio, K. R. & Wieczorek, A. M. Fine-scale spatial genetic structure and dispersal among spotted salamander (Ambystoma maculatum) breeding populations. Mol. Ecol. 16, 257–274 (2007).

76. Emel, S. L. & Storfer, A. A decade of amphibian population genetic studies: synthesis and recommendations. Conserv. Genet. 13, 1685–1689 (2012).

77. Meier, J. I. et al. The coincidence of ecological opportunity with hybridization explains rapid adaptive radiation in Lake Mweru cichlid fishes. Nat. Commun. 10, (2019).

78. Rico, C. & Turner, G. F. Extreme microallopatric divergence in a cichlid species from Lake Malawi. Mol. Ecol. 11, 1585–1590 (2002).

79. Church, S. A., Kraus, J. M., Mitchell, J. C., Church, D. R. & Taylor, D. R. Evidence for Multiple Pleistocene Refugia in the Postglacial Expansion of the Eastern Tiger salamander, Ambystoma Tigrinum Tigrinum. Evolution (N. Y). 57, 372–383 (2003).

80. Crespi, E. J., Rissler, L. J. & Browne, R. A. Testing Pleistocene refugia theory: phylogeographical analysis of Desmognathus wrighti, a high-elevation salamander in the southern Appalachians. Mol. Ecol. 12, 969–984 (2003).

81. Steele, C. A. & Storfer, A. Phylogeographic incongruence of codistributed amphibian species based on small differences in geographic distribution. Mol. Phylogenet. Evol. 43, 468–479 (2007).

82. Zamudio, K. R. & Savage, W. K. Historical Isolation, Range Expansion, and Secondary Contact of Two Highly Divergent Mitochondrial Lineages in Spotted Salamanders (Ambystoma maculatum). Evolution (N. Y). 57, 1631–1652 (2003).

83. Alexandrino, J., Froufe, E., Arntzen, J. W. & Ferrand, N. Genetic subdivision, glacial refugia and postglacial recolonization in the golden-striped salamander, Chioglossa lusitanica (Amphibia: Urodela). Mol. Ecol. 9, 771–781 (2000).

84. Steinfartz, S., Veith, M. & Tautz, D. Mitochondrial sequence analysis of Salamandra taxa suggests old splits of major lineages and postglacial recolonizations of Central Europe from distinct source populations of Salamandra salamandra. Mol. Ecol. 9, 397–410 (2000).

85. Mattoccia, M., Marta, S., Romano, A. & Sbordoni, V. Phylogeography of an Italian endemic salamander (genus Salamandrina): glacial refugia, postglacial expansions, and secondary contact. Biol. J. Linn. Soc. 104, 903–992 (2011).

86. Kozak, K. H. & Wiens, J. J. Niche conservatism drives elevational diversity patterns in Appalachian salamanders. Am. Nat. 176, 40–54 (2010).

87. Parmesan, C. & Yohe, G. A globally coherent fingerprint of climate change impacts across natural systems. Nature 421, 37–42 (2003).

88. Root, T. L., Hall, P., K. R., S. & S. H. Rosenzweig C. & Pounds, J. A. Fingerprints of global warming on wild animals and plants. Nature 421, 57–60 (2003).

89. Kelly, A. E. & Goulden, M. L. Rapid shifts in plant distribution with recent climate change. Proc. Natl. Acad. Sci. 105, 11823–11826 (2008).

90. Chen, I. C., Hill, J. K., Ohlemüller, R., Roy, D. B. & Thomas, C. D. Rapid range shifts of species associated with high levels of climate warming. Science (80-.). 333, 1024–1026 (2011).

91. Elsen, P. R. & Tingley, M. W. Global mountain topography and the fate of montane species under climate change. Nat. Clim. Chang. 5, 772–776 (2015).

92. Garroway, C. J. et al. Climate change induced hybridization in flying squirrels. Glob. Chang. Biol. 16, 113–121 (2010).

93. Hoffmann, A. A. & Sgrò, C. M. Climate change and evolutionary adaptation. Nature 470, 479–85 (2011).

94. Chunco, A. J. Hybridization in a warmer world. Ecol. Evol. 4, 2019–2031 (2014).

95. McQuillan, M. A. & Rice, A. M. Differential effects of climate and species interactions on range limits at a hybrid zone: potential direct and indirect impacts of climate change. Ecol. Evol. 5, 5120–5137 (2015).

96. Pereira, R. J., Martínez-Solano, I. & Buckley, D. Hybridization during altitudinal range shifts: nuclear introgression leads to extensive cyto-nuclear discordance in the fire salamander. Mol. Ecol. 25, 1551–1565 (2016).

97. Seehausen, O. Conservation: losing biodiversity by reverse speciation. Curr. Biol. 16, 334–R337 (2006).

98. Seehausen, O., Takimoto, G., Roy, D. & Jokela, J. Speciation reversal and biodiversity dynamics with hybridization in changing environments. Mol. Ecol. 17, 30–44 (2008).

99. Vonlanthen, P. & others. Eutrophication causes speciation reversal in whitefish adaptive radiations. Nature 482, 357–362 (2012).

100. Hoskin, C. J., Higgie, M., McDonald, K. R. & Moritz, C. Reinforcement drives rapid allopatric speciation. Nature 437, 1353–1356 (2005).

101. Pfennig, K. S. Reinforcement as an initiator of population divergence and speciation. Curr. Zool. 62, 145–154 (2016).

102. Arnold, S. J., Reagan, N. L. & Verrell., P. A. Reproductive isolation and speciation in plethodontid salamanders. Herpetologica 49, 216–228 (1993).

103. Reagan, N. L. Evolution of sexual isolation in salamanders of the genus Plethodon (Doctoral dissertation). (University of Chicago, 1992).

104. Davis, M. P., Midford, P. E. & Maddison, W. Exploring power and parameter estimation of the BiSSE method for analyzing species diversification. BMC Evol. Biol. 13, 38 (2013).

105. Freyman, W. A. & Höhna, S. Stochastic character mapping of state-dependent Diversification reveals the tempo of evolutionary decline in self-compatible Onagraceae lineages. Syst. Biol. 68, 505–519 (2019).

106. Rabosky, D. L. Extinction rates should not be estimated from molecular phylogenies. Evolution (N. Y). 64, 1816–1824 (2010).

107. Rabosky, D. L. Challenges in the estimation of extinction from molecular phylogenies: a response to Beaulieu and O’Meara. Evolution (N. Y). 70, 218–228 (2016).

108. Kearney, M. Hybridization, glaciation and geographical parthenogenesis. Trends Ecol. Evol. 20, 495–502 (2005).

109. Willis, B. L., van Oppen, M. J., Miller, D. J., Vollmer, S. V & Ayre, D. J. The role of hybridization in the evolution of reef corals. Annu. Rev. Ecol. Evol. Syst 37, 489–517 (2006).

110. Schmid, M., Evans, B. J. & Bogart, J. P. Polyploidy in Amphibia. Cytogenetic and Genome Research 145, 315–330 (2015).

111. Runemark, A., Vallejo-Marin, M. & Meier, J. I. Eukaryote hybrid genomes. PLOS Genet. 15, e1008404 (2019).

