## Supplement for "Hybridizing salamanders experience accelerated diversification"

### Supplementary Materials

#### *Materials and Methods*

##### Assessment of Trait-Dependent Diversification: HiSSE

Through the incorporation of hidden states, the model may be reparameterized to generate a set of competing models, varied in the number of hidden states contained in the model, as well as whether diversification and/or transition rates are allowed to vary among states. Implemented in a likelihood framework, information theoretic approaches (e.g. AIC, AICc) may thus be used to assess fit among competing models. We implemented 14 competing models that varied in the number of hidden states, as well as in the number of unique sets of transition and diversification rates included in the model (Supplementary Table S1). Each of these models were assessed using four datasets described in the manuscript. Briefly, these include 1) all species, 2) only species in sympatry with another species, 3) only plethodontids, and 4) only non-plethodontids. We did not implement HiSSE models assuming equal transition rates among states as the rarity and dispersion of our trait across the tree (Fig. S1) would force the estimation of unrealistically high rates of both speciation and extinction to explain their distribution were transition rates forced to be equal.

By default, HiSSE produces simple transformations of the Speciation Rate ($\lambda$) and Extinction Rate ($\mu$). Specifically, it produces rate estimates of the Extinction Fraction ($\varepsilon = \mu \div \lambda$) and Turnover Rate ($\tau = \lambda + \mu$). While the untransformed rate estimates ($\lambda$ & $\mu$) are conventionally reported, difficulties in obtaining precise estimates of extinction rates in the SSE framework are well documented[63]. However, this may be ameliorated to an extent through the transformation of rate estimates into ε and τ[61]. Consequently, we report each of the four diversification rates in addition to Net Diversification which is simply $\lambda - \mu$. Transition rates among character states are reported, although we are less interested in these than we are in estimates for diversification rates.

We choose not to use the semi-parametric FiSSE[112], as this method necessitates symmetric transition rates among character-states. As our character-states exhibit dramatic asymmetry in transition rates, this approach is unsuitable. Further, the DR statistic[113] used as a heuristic for speciation rate is only accurate under a pure-birth model and does not capture the net-diversification rate[114]. Thus, if extinction plays an important role in the diversification process, FiSSE cannot detect it and may be confounded by it.

**Additional Datasets**

To define sympatry, we needed to collect range polygons, calculate range areas, and calculate areas of intersection for all pairwise combinations of species. Using these range areas and range intersections, we then were able to calculate the percentage of overlap for all pairwise combinations. Range polygons of all available Caudates were downloaded from the IUCN Redlist Webpage[115]. We then converted these polygons to the Mollweide equal area projection, and we then calculated range areas and areas of pairwise intersections using the R packages rgdal v1.2-8[116], PBSmapping v2.70.4[117], and fuzzySim v1.7.9[118]. The resultant matrix was then used to manually calculate percentage overlaps, and only those species sharing at least 10% of their range with another species were retained. Because the *Plethodon glutinosus* group (Figure 1B) has been the subject of extensive taxonomic revision using molecular data (e.g. refs [39–42]), there exists the distinct possibility that the signature between hybridization and diversification rates in this group leads to biased inferenced. To account for this potential confounding effect, we re-analyzed the complete tree excluding the *Plethodon glutinosus* group, using both the narrow and broad definitions of hybridization.

**Assessing Diversification Rate Differences: All Datasets**

In lieu of direct model comparison and reporting parameters under a single model, we adopt the approach of Caetano et al.[65] and report model-averaged parameter estimates. Although the inclusion of hidden states and non-trivial null models greatly improves HiSSE's model rejection properties over related SSE models[61], comparing model-averaged param[65]eter estimates between states enables HiSSE to distinguish between trait-dependent and independent diversification under the most pathological circumstances[65]. Consequently, we report model-averaged parameter estimates for hybridizing and non-hybridizing species. Using this approach, we obtain lineage-specific diversification rates which we then use to conduct an empirical test of diversification rate differences. By making all possible comparisons between non-hybridizing and hybridizing lineages as a ratio (NH/H), we obtain an empirical p-value that communicates the probability that non-hybridizing lineages possess diversification rates that are greater-than or less-than those of hybridizing lineages.

**Incomplete taxon-sampling**

As implemented by HiSSE, sampling fractions are specified for each state, rather than for taxonomic groups. Importantly, it is assumed that sampling of each state is random; to test whether non-random sampling could bias our results[119], we ask whether clade-specific sampling fraction predicts prevalence of hybridization. We conduct this test at both the family and genus level (Supplementary Fig. S5), regressing clade-specific sampling fraction against the (sampled) proportion of each clade that is currently hybridizing.

**Sensitivity to Phylogenetic Uncertainty**

We next tested whether our results are sensitive to phylogenetic uncertainty. Topological uncertainty for the tree used elsewhere in this study[32] was quantified using Shimodaira–Hasegawa-Like implementation of the approximate likelihood-ratio test. Consequently, we sampled 100 trees from the posterior distribution of trees inferred by Jetz and Pyron[120] (Supplementary Fig. S6). These trees were inferred using imputed data for many taxa and so were not used for other analyses herein. Using these 100 trees, we repeated our HiSSE analysis using all species and assuming 20% of species hybridize according to our narrow definition of hybridization. We subsequently 1) summarized model support across replicate analyses (Supplementary Table S15), and 2) calculated the model averaged diversification rates for hybridizing and non-hybridizing taxa from each replicate analysis (Supplementary Fig. S7). This pooled set of model averaged diversification rates was then analyzed using a Welch's two-sample t-test in R.

#### *Results*

**Incomplete taxon-sampling**

Non-random taxonomic sampling has also been shown to undermine the performance of macroevolutionary methods such as those that estimate diversification rates[119]. However, we find no evidence that clade-specific sampling fraction predicts the frequency of hybridization (Supplementary Fig. S5) in the respective families (Adjusted $R^2$ = 5.2e-7, $P$ = 1) or genera (Adjusted $R^2$ = 3.0e-3, $P$ = 0.67).

**Sensitivity to Phylogenetic Uncertainty**

Our findings appear robust to phylogenetic uncertainty. Using 100 phylogenies drawn from the posterior distribution of Jetz & Pyron[120], a model of trait-independent diversification was never favored (Supplementary Table S15). In other words, trait-dependent diversification was always supported as the best-fit model. Further, as in our main analyses, a HiSSE model was always favored over a simpler BiSSE model. Investigation of model averaged parameter estimates reveals that the mean net-diversification rate of hybridizing salamanders is always greater than those of non-hybridizing salamanders (Supplementary Fig. S7). As with the main analyses, this is driven by concurrent increases in speciation rates and decreases in extinction rates.

**Exclusion of *Plethodon glutinosus* group**

Reanalysis following exclusion of the *Plethodon glutinosus* group supported our main finding that hybridizing species of salamanders experience heightened diversification rates (Supplementary Fig. S13 & S14, Supplementary Table S17). That is, when analyzed under both the narrow and broad definitions of hybridization, models of trait dependent diversification in which hybridizing salamanders experience heightened rates of diversification were overwhelmingly supported.

#### *Discussion*

We believe that the frequency of hybridization in nature is equal to or greater than that represented in the literature due to the substantial effort necessary to document hybridization. Thus, although we suspect that there is a bias in the detection of hybridization towards well studied species in our dataset, many of the species included in our phylogeny have poorly understood ecologies (e.g. Bolitoglossans, Hybobiids). Consequently, we anticipate that the frequency of detection of hybridization in these understudied groups is likely to be equally underestimated for those species included in our phylogeny, as well as for those that are not.

##### Statistical properties of SSE Methods

Davis et al.[104] documented the potential artifactual consequences of extreme tip-ratio biases in character states. Specifically, they state that "In general, investigators should be cautious using the BiSSE method when one of the binary characters in question is exceedingly rare in their data sets (less than 10%)." As hybridization is observed in ~20% of taxa (tip ratio ~5:1), our dataset thus does not meet their criteria for concern. Further, the tip ratio biases most strongly influence inference of rates of character change followed by extinction rates. As the changes in net diversification in our analyses are most strongly dictated by changes in speciation rather than extinction rates, we have little reason to be concerned that the overall findings of our study are confounded by tip-ratio asymmetries.

Lastly, we are not concerned about the impacts of phylogenetic pseudoreplication in our trait. That is, autapomorphies of rapidly diversifying clades may be spuriously attributed to the rapidity of diversification by SSE methods[112,121]. As our trait is broadly distributed about order Caudata, the potential for phylogenetic pseudoreplication is negligible.

**Future Directions**

With our data (12 genes), we cannot test the hypothesis that ancient hybridization facilitated the diversification of salamanders through the detection of genealogical discordance at the base of the radiation. The methods best suited to testing this hypothesis, namely the ABBA-BABA class of methods which test for an excess of shared, derived allelic variants using a three-, four- or five-taxon tree (e.g. refs. [122–125]), require genomic datasets larger than our own to have appreciable power[126,127]. Thus, tests of Seehausen's hypotheses are only feasible at the macroevolutionary scale using phylogenomic datasets, an avenue of research we encourage.

### References


1. Barton, N. H. & Hewitt, G. M. Analysis of hybrid zones. *Annu. Rev. Ecol. Syst.* **16,** 113–148 (1985).
2. Darwin, C. *On the Origin of Species by Means of Natural Selection. (Murray*. (1859).
3. Fisher, R. *The Genetical Theory of Natural Selection. (Clarendon Press*. (1930).
4. Mayr, E. *Animal species and evolution. (Harvard University Press*. (1963).
5. Futuyma, D. J. On the role of species in anagenesis. *Am. Nat.* **130,** 465–473 (1987).
6. Seehausen, O. Hybridization and adaptive radiation. *Trends Ecol. Evol.* **19,** 198–207 (2004).
7. Grant, P. R. & Grant, B. R. Hybridization of bird species. *Science (80-. ).* **256,** 193–197 (1992).
8. Seehausen, O., Van Alphen, J. J. & Witte, F. Cichlid fish diversity threatened by eutrophication that curbs sexual selection. *Science (80-. ).* **277,** 1808–1811 (1997).
9. Turelli, M., Barton, N. H. & Coyne, J. A. Theory and speciation. *Trends Ecol. Evol.* **16,** 330–343 (2001).
10. Buerkle, C. A., Morris, R. J., Asmussen, M. A. & Rieseberg, L. H. The likelihood of homoploid hybrid speciation. *Heredity (Edinb).* **84,** 441–451 (2000).
11. Dowling, T. E. & Secor, C. L. the Role of Hybridization and Introgression in the Diversification of Animals. *Annu. Rev. Ecol. Syst.* **28,** 593–619 (1997).
12. Rieseberg, L. H., Archer, M. A. & Wayne, R. K. Transgressive segregation, adaptation and speciation. *Heredity (Edinb).* **83,** 363 (1999).
13. Soltis, P. S. & Soltis, D. E. The role of hybridization in plant speciation. *Annu. Rev. Plant Biol.* **60,** 561–588 (2009).
14. Coyne, J. A. & Orr, H. A. *Speciation. (Sinauer Associates Sunderland*. (MA, 2004).
15. Francillon-Vieillot, H., Arntzen, J. W. & Geraudie, J. Age, Growth and Longevity of

Sympatric Triturus cristatus, T. marmoratus and Their Hybrids (Amphibia, Urodela): A Skeletochronological Comparison. *J. Herpetol.* **24,** 13 (1990).

16. Arntzen, J. W., Jehle, R., Bardakci, F., Burke, T. & Wallis, G. P. Asymmetric Viability of Reciprocal-Cross Hybrids Between Crested and Marbled News (Triturus cristatus and T. marmoratus). *Evolution (N. Y).* **63,** 1191–1202 (2009).
17. Marques, D. A., Meier, J. I. & Seehausen, O. *A combinatorial view on speciation and adaptive radiation*. (Trends in ecology & evolution, 2019).
18. Mallet, J. Hybrid speciation. *Nature* **446,** 279–83 (2007).
19. Mavárez, J. & Linares, M. Homoploid hybrid speciation in animals. *Mol. Ecol.* **17,** 4181–4185 (2008).
20. Gross, B. & Rieseberg, L. H. The ecological genetics of homoploid hybrid speciation. *J. Hered.* **96,** 241–252 (2005).
21. Schumer, M., Rosenthal, G. G. & Andolfatto, P. How common is homoploid hybrid speciation? *Evolution (N. Y).* **68,** 1553–1560 (2014).
22. Mavárez, J. *et al.* Speciation by hybridization in Heliconius butterflies. *Nature* **441,** 868–871 (2006).
23. Wiens, J. J., Engstrom, T. N. & Chippindale, P. T. Rapid diversification, incomplete isolation, and the ``speciation clock'' in North {A}merican salamanders (genus Plethodon): testing the hybrid swarm hypothesis of rapid radiation. *Evolution (N. Y).* **60,** 2585–2603 (2006).
24. Keller, I. *et al.* Population genomic signatures of divergent adaptation, gene flow and hybrid speciation in the rapid radiation of Lake Victoria cichlid fishes. *Mol. Ecol.* **22,** 2848–2863 (2013).
25. Meier, J. I. *et al.* Ancient hybridization fuels rapid cichlid fish adaptive radiations. *Nat. Commun.* **8,** 14363 (2017).
26. Grant, P. R. & Grant, B. R. *No Title*. (Darwin's finches on Daphne Major island. (Princeton University Press, 2014).
27. MacLeod, A. *et al.* Hybridization masks speciation in the evolutionary history of the Galápagos marine iguana. *Proc. R. Soc. B* **282,** 20150 (2015).
28. Yakimowski, S. B. & Rieseberg, L. H. The role of homoploid hybridization in evolution: a century of studies synthesizing genetics and ecology. *Am. J. Bot.* **101,** 1247–1258 (2014).
29. Seehausen, O. & others. Genomics and the origin of species. *Nat. Rev. Genet.* **15,** 176–192 (2014).
30. Gompert, Z. & Buerkle, C. A. What, if anything, are hybrids: enduring truths and challenges associated with population structure and gene flow. *Evol. Appl.* **9,** 909–923 (2016).
31. Berkeley, U. of C. *Species by the numbers.* (2018). at <http://www.amphibiaweb.org/amphibian/speciesnums.html>
32. Pyron, R. A. Biogeographic analysis reveals ancient continental vicariance and recent oceanic dispersal in amphibians. *Syst. Biol.* **63,** 779–797 (2014).
33. Wiens, J. J. Speciation and Ecology Revisited: Phylogenetic Niche Conservatism and the Origin of Species. *Evolution (N. Y).* **58,** 193–197 (2004).
34. Kozak, K. H., Weisrock, D. W. & Larson, A. Rapid lineage accumulation in a non-adaptive radiation: Phylogenetic analysis of diversification rates in eastern North

American woodland salamanders (Plethodontidae: Plethodon). *Proc. R. Soc. B Biol. Sci.* **273,** 539–546 (2006).

35. Wake, D. B. Problems with Species: Patterns and Processes of Species Formation in Salamanders. *Ann. Missouri Bot. Gard.* **93,** 8–23 (2006).
36. Vences, M. & Wake, D. B. in *Amphibian Biology, Vol. 6, Systematics* (eds. Heatwole, H. H. & Tyler, M.) 2613–2669 (Surrey Beatty & Sons, Chipping Norton, Australia, 2007).
37. Wake, D. B. What Salamanders have Taught Us about Evolution. (2009). doi:10.1146/annurev.ecolsys.39.110707.173552
38. Blankers, T., Adams, D. C. & Wiens, J. J. Ecological radiation with limited morphological diversification in salamanders. *J. Evol. Biol.* **25,** 634–646 (2012).
39. Highton, R. Speciation in eastern North {A}merican salamanders of the genus Plethodon. *Annu. Rev. Ecol. Syst.* **26,** 579–600 (1995).
40. Highton, R. Geographic protein variation and speciation in the Plethodon dorsalis complex. *Herpetologica* **1997,** 345–356 (1997).
41. Highton, R. Geographic protein variation and speciation in the salamanders of the Plethodon cinereus group with the description of two new species. *Herpetologica* **1999,** 43–90 (1999).
42. Highton, R. & Peabody, R. in *The Biology of Plethodontid Salamanders.* (eds. Bruce, R. C., Jaeger, R. G. & Houck, L. D.) 31–93 (Kluwer Academic/Plenum Publishers, 2000).
43. Wake, D. B. & Jockusch, E. L. in *The Biology of Plethodontid Salamanders* 95–119 (Springer US, 2000). doi:10.1007/978-1-4615-4255-1_4
44. Patton, A. *et al.* A New Green Salamander in the Southern Appalachians: Evolutionary History of Aneides aeneus and Implications for Management and Conservation with the Description of a Cryptic Microendemic Species. *Copeia* **107,** (2019).
45. Rundell, R. J. & Price, T. D. Adaptive radiation, nonadaptive radiation, ecological speciation and nonecological speciation. *Trends Ecol. Evol.* **24,** 394–399 (2009).
46. Czekanski-Moir, J. E. & Rundell, R. J. The ecology of nonecological speciation and nonadaptive radiations. *Trends Ecol. Evol.* (2019).
47. Garcia-Paris, M., Good, D. A., Parra-Olea, G. & Wake, D. B. Biodiversity of Costa Rican salamanders: Implications of high levels of genetic differentiation and phylogeographic structure for species formation. *Proc. Natl. Acad. Sci.* **97,** 1640–1647 (2000).
48. Parra-Olea, G. & Wake, D. B. Extreme morphological and ecological homoplasy in tropical salamanders. *Proc. Natl. Acad. Sci. U. S. A.* **98,** 7888–7891 (2001).
49. Vieites, D. R., Min, M. S. & Wake, D. B. Rapid diversification and dispersal during periods of global warming by plethodontid salamanders. *Proc. Natl. Acad. Sci. U. S. A.* **104,** 19903–19907 (2007).
50. Parra-Olea, G., Windfield, J. C., Velo-Antón, G. & Zamudio, K. R. Isolation in habitat refugia promotes rapid diversification in a montane tropical salamander. *J. Biogeogr.* **39,** 353–370 (2012).
51. Bryson, R. W. *et al.* Phylogenomic insights into the diversification of salamanders in the Isthmura bellii group across the Mexican highlands México View project Phylogeography and contact zones in Ctenosaura pectinata View project Phylogenomic insights into the diversification of salamanders in the Isthmura bellii group across the Mexican highlands. *Artic. Mol. Phylogenetics Evol.* (2018). doi:10.1016/j.ympev.2018.03.024

52. Gillespie, R. G. *et al.* Comparing Adaptive Radiations Across Space, Time, and Taxa. *J. Hered.* **89154,** 14853 (2020).
53. Stamatakis, A. RAxML-VI-HPC: maximum likelihood-based phylogenetic analyses with thousands of taxa and mixed models. *Bioinformatics* **22,** 2688–2690 (2006).
54. Smith, S. A. & O'meara, B. C. treePL: divergence time estimation using penalized likelihood for large phylogenies. *Bioinformatics* **28,** 2689–2690 (2012).
55. Paradis, E., Claude, J. & Strimmer, K. APE: analyses of phylogenetics and evolution in {R} language. *Bioinformatics* **20,** 289–290 (2004).
56. Liu, F. G. R., Moler, P. E. & Miyamoto, M. M. Phylogeography of the salamander genus Pseudobranchus in the southeastern United States. *Mol. Phylogenet. Evol.* **39,** 149–159 (2006).
57. Fu, J. & Zeng, X. How many species are in the genus Batrachuperus? A phylogeographical analysis of the stream salamanders (family Hynobiidae) from southwestern China. *Mol. Ecol.* **17,** 1469–1488 (2008).
58. Yoshikawa, N., Matsui, M., Nishikawa, K., Misawa, Y. & Tanabe, S. Allozymic Variation in the Japanese Clawed Salamander, Onychodactylus japonicus (Amphibia: Caudata: Hynobiidae), with Special Reference to the Presence of Two Sympatric Genetic Types. *Zoolog. Sci.* **27,** 33–40 (2010).
59. Yoshikawa, N., Matsui, M. & Nishikawa, K. Genetic Structure and Cryptic Diversity of Onychodactylus japonicus (Amphibia, Caudata, Hynobiidae) in Northeastern Honshu, Japan, as Revealed by Allozymic Analysis. *Zoolog. Sci.* **29,** 229–237 (2012).
60. Yoshikawa, N. & Matsui, M. Two new Salamanders of the genus Onychodactylus from Eastern Honshu, Japan (Amphibia, Caudata, Hynobiidae). (2014). doi:10.11646/zootaxa.3866.1.3
61. Beaulieu, J. M. & O'Meara, B. C. Detecting hidden diversification shifts in models of trait dependent speciation and extinction. *Syst. Biol.* **65,** 583–601 (2016).
62. Stadler, T. How can we improve accuracy of macroevolutionary rate estimates? *Syst. Biol.* **62,** 321–329 (2013).
63. Beaulieu, J. M. & O'Meara, B. C. Extinction can be estimated from moderately sized molecular phylogenies. **69,** 1036–1043 (2015).
64. Rabosky, D. L. & Goldberg, E. E. Model inadequacy and mistaken inferences of trait-dependent speciation. *Syst. Biol.* **64,** 340–55 (2015).
65. Caetano, D. S., O'Meara, B. C. & Beaulieu, J. M. Hidden state models improve state-dependent diversification approaches, including biogeographical models. *Evolution (N. Y).* (2018).
66. Barraclough, T. G., Harvey, P. H. & Nee, S. Rate of rbcL gene sequence evolution and species diversification in flowering plants (angiosperms). Proceedings of the Royal Society of London. *Ser. B. Biol. Sci.* **263,** 589–591 (1996).
67. R Core Team. R: A Language and Environment for Statistical Computing. (2018). at <https://www.r-project.org/>
68. Hewitt, G. M. Some genetic consequences of ice ages, and their role in divergence and speciation. *Biol. J. Linn. Soc.* **58,** 247–276 (1996).
69. Kozak, K. H., Blaine, R. A. & Larson, A. Gene lineages and eastern North {A}merican palaeodrainage basins: phylogeography and speciation in salamanders of the Eurycea

bislineata species complex. *Mol. Ecol.* **15,** 191–207 (2006).
70. Shepard, D. B. & Burbrink, F. T. Phylogeographic and demographic effects of Pleistocene climatic fluctuations in a montane salamander, Plethodon fourchensis. *Mol. Ecol.* **18,** 2243–2262 (2009).
71. Walls, S. C. The role of climate in the dynamics of a hybrid zone in Appalachian salamanders. *Glob. Chang. Biol.* **15,** 1903–1910 (2009).
72. Rice, W. R. & Hostert, E. E. Laboratory experiments on speciation: what have we learned in 40 years? **47,** 1637–1653 (1993).
73. Servedio, M. R. & Noor, M. A. The role of reinforcement in speciation: theory and data. *Annu. Rev. Ecol. Evol. Syst.* **34,** 339–364 (2003).
74. Matute, D. R. & Ortiz-Barrientos, D. Speciation: The strength of natural selection driving reinforcement. *Curr. Biol.* **24,** 955–957 (2014).
75. Zamudio, K. R. & Wieczorek, A. M. Fine-scale spatial genetic structure and dispersal among spotted salamander (Ambystoma maculatum) breeding populations. *Mol. Ecol.* **16,** 257–274 (2007).
76. Emel, S. L. & Storfer, A. A decade of amphibian population genetic studies: synthesis and recommendations. *Conserv. Genet.* **13,** 1685–1689 (2012).
77. Meier, J. I. *et al.* The coincidence of ecological opportunity with hybridization explains rapid adaptive radiation in Lake Mweru cichlid fishes. *Nat. Commun.* **10,** (2019).
78. Rico, C. & Turner, G. F. Extreme microallopatric divergence in a cichlid species from Lake Malawi. *Mol. Ecol.* **11,** 1585–1590 (2002).
79. Church, S. A., Kraus, J. M., Mitchell, J. C., Church, D. R. & Taylor, D. R. Evidence for Multiple Pleistocene Refugia in the Postglacial Expansion of the Eastern Tiger salamander, Ambystoma Tigrinum Tigrinum. *Evolution (N. Y).* **57,** 372–383 (2003).
80. Crespi, E. J., Rissler, L. J. & Browne, R. A. Testing Pleistocene refugia theory: phylogeographical analysis of Desmognathus wrighti, a high-elevation salamander in the southern Appalachians. *Mol. Ecol.* **12,** 969–984 (2003).
81. Steele, C. A. & Storfer, A. Phylogeographic incongruence of codistributed amphibian species based on small differences in geographic distribution. *Mol. Phylogenet. Evol.* **43,** 468–479 (2007).
82. Zamudio, K. R. & Savage, W. K. Historical Isolation, Range Expansion, and Secondary Contact of Two Highly Divergent Mitochondrial Lineages in Spotted Salamanders (Ambystoma maculatum). *Evolution (N. Y).* **57,** 1631–1652 (2003).
83. Alexandrino, J., Froufe, E., Arntzen, J. W. & Ferrand, N. Genetic subdivision, glacial refugia and postglacial recolonization in the golden-striped salamander, Chioglossa lusitanica (Amphibia: Urodela). *Mol. Ecol.* **9,** 771–781 (2000).
84. Steinfartz, S., Veith, M. & Tautz, D. Mitochondrial sequence analysis of Salamandra taxa suggests old splits of major lineages and postglacial recolonizations of Central Europe from distinct source populations of Salamandra salamandra. *Mol. Ecol.* **9,** 397–410 (2000).
85. Mattoccia, M., Marta, S., Romano, A. & Sbordoni, V. Phylogeography of an Italian endemic salamander (genus Salamandrina): glacial refugia, postglacial expansions, and secondary contact. *Biol. J. Linn. Soc.* **104,** 903–992 (2011).
86. Kozak, K. H. & Wiens, J. J. Niche conservatism drives elevational diversity patterns in

Appalachian salamanders. *Am. Nat.* **176,** 40–54 (2010).

87. Parmesan, C. & Yohe, G. A globally coherent fingerprint of climate change impacts across natural systems. *Nature* **421,** 37–42 (2003).
88. Root, T. L., Hall, P., K. R., S. & S. H. Rosenzweig C. & Pounds, J. A. Fingerprints of global warming on wild animals and plants. *Nature* **421,** 57–60 (2003).
89. Kelly, A. E. & Goulden, M. L. Rapid shifts in plant distribution with recent climate change. *Proc. Natl. Acad. Sci.* **105,** 11823–11826 (2008).
90. Chen, I. C., Hill, J. K., Ohlemüller, R., Roy, D. B. & Thomas, C. D. Rapid range shifts of species associated with high levels of climate warming. *Science (80-. ).* **333,** 1024–1026 (2011).
91. Elsen, P. R. & Tingley, M. W. Global mountain topography and the fate of montane species under climate change. *Nat. Clim. Chang.* **5,** 772–776 (2015).
92. Garroway, C. J. *et al.* Climate change induced hybridization in flying squirrels. *Glob. Chang. Biol.* **16,** 113–121 (2010).
93. Hoffmann, A. A. & Sgrò, C. M. Climate change and evolutionary adaptation. *Nature* **470,** 479–85 (2011).
94. Chunco, A. J. Hybridization in a warmer world. *Ecol. Evol.* **4,** 2019–2031 (2014).
95. McQuillan, M. A. & Rice, A. M. Differential effects of climate and species interactions on range limits at a hybrid zone: potential direct and indirect impacts of climate change. *Ecol. Evol.* **5,** 5120–5137 (2015).
96. Pereira, R. J., Martínez-Solano, I. & Buckley, D. Hybridization during altitudinal range shifts: nuclear introgression leads to extensive cyto-nuclear discordance in the fire salamander. *Mol. Ecol.* **25,** 1551–1565 (2016).
97. Seehausen, O. Conservation: losing biodiversity by reverse speciation. *Curr. Biol.* **16,** 334-R337 (2006).
98. Seehausen, O., Takimoto, G., Roy, D. & Jokela, J. Speciation reversal and biodiversity dynamics with hybridization in changing environments. *Mol. Ecol.* **17,** 30–44 (2008).
99. Vonlanthen, P. & others. Eutrophication causes speciation reversal in whitefish adaptive radiations. *Nature* **482,** 357–362 (2012).
100. Hoskin, C. J., Higgie, M., McDonald, K. R. & Moritz, C. Reinforcement drives rapid allopatric speciation. *Nature* **437,** 1353–1356 (2005).
101. Pfennig, K. S. Reinforcement as an initiator of population divergence and speciation. *Curr. Zool.* **62,** 145–154 (2016).
102. Arnold, S. J., Reagan, N. L. & Verrell., P. A. Reproductive isolation and speciation in plethodontid salamanders. *Herpetologica* **49,** 216–228 (1993).
103. Reagan, N. L. *Evolution of sexual isolation in salamanders of the genus Plethodon (Doctoral dissertation)*. (University of Chicago, 1992).
104. Davis, M. P., Midford, P. E. & Maddison, W. Exploring power and parameter estimation of the BiSSE method for analyzing species diversification. *BMC Evol. Biol.* **13,** 38 (2013).
105. Freyman, W. A. & Höhna, S. Stochastic character mapping of state-dependent Diversification reveals the tempo of evolutionary decline in self-compatible Onagraceae lineages. *Syst. Biol.* **68,** 505–519 (2019).
106. Rabosky, D. L. Extinction rates should not be estimated from molecular phylogenies. *Evolution (N. Y).* **64,** 1816–1824 (2010).

107. Rabosky, D. L. Challenges in the estimation of extinction from molecular phylogenies: a response to Beaulieu and O'Meara. *Evolution (N. Y).* **70,** 218–228 (2016).
108. Kearney, M. Hybridization, glaciation and geographical parthenogenesis. *Trends Ecol. Evol.* **20,** 495–502 (2005).
109. Willis, B. L., van Oppen, M. J., Miller, D. J., Vollmer, S. V & Ayre, D. J. The role of hybridization in the evolution of reef corals. *Annu. Rev. Ecol. Evol. Syst* **37,** 489–517 (2006).
110. Schmid, M., Evans, B. J. & Bogart, J. P. Polyploidy in Amphibia. *Cytogenetic and Genome Research* **145,** 315–330 (2015).
111. Runemark, A., Vallejo-Marin, M. & Meier, J. I. Eukaryote hybrid genomes. *PLOS Genet.* **15,** e1008404 (2019).
112. Rabosky, D. L. & Goldberg, E. E. FiSSE: A simple nonparametric test for the effects of a binary character on lineage diversification rates. doi:10.1111/evo.13227
113. Jetz, W., Thomas, G., Joy, J., Hartmann, J. & Mooers, A. The global diversity of birds in space and time. *Nature* **491,** 444–448 (2012).
114. Belmaker, J. & Jetz, W. Relative roles of ecological and energetic constraints, diversification rates and region history on global species richness gradients. *Ecol. Lett.* **18,** 563–571 (2015).
115. *Spatial data download.* (2017). at <http://www.iucnredlist.org/technical-documents/spatial-data/>
116. Bivand, R., Keitt, T. & Rowlingson, B. rgdal: Bindings for the 'Geospatial' Data Abstraction Library. *R Packag. version* **1,** 2–13 (2017).
117. Schnute, J. T., Boers, N. & Haigh, R. PBSmapping: Mapping Fisheries Data and Spatial Analysis Tools. (2017).
118. Barbosa, A. M. fuzzysim: applying fuzzy logic to binary similarity indices in ecology. *Methods Ecol. Evol.* **6,** 853–858 (2015).
119. Hua, X. & Lanfear, R. The influence of non-random species sampling on macroevolutionary and macroecological inference from phylogenies. *Methods Ecol. Evol.* **9,** 1353–1362 (2018).
120. Jetz, W. & Pyron, R. A. The interplay of past diversification and evolutionary isolation with present imperilment across the amphibian tree of life. *Nat. Ecol. Evol.* **2,** 850 (2018).
121. Maddison, W. P. & FitzJohn, R. G. The unsolved challenge to phylogenetic correlation tests for categorical characters. *Syst. Biol.* **64,** 127–136 (2014).
122. Hahn, M. W. & Hibbins, M. S. A Three-Sample Test for Introgression. *Mol. Biol. Evol.* **36,** 2878–2882 (2019).
123. Green, R. E. & others. A draft sequence of the Neandertal genome. *Science (80-. ).* **328,** 710–722 (2010).
124. Durand, E. Y., Patterson, N., Reich, D. & Slatkin, M. Testing for ancient admixture between closely related populations. *Mol. Biol. Evol.* **28,** 2239–2252 (2011).
125. Pease, J. B. & Hahn, M. W. Detection and polarization of introgression in a five-taxon phylogeny. *Syst. Biol.* **64,** 651–662 (2015).
126. Martin, S. H., Davey, J. W. & Jiggins, C. D. Evaluating the use of ABBA--BABA statistics to locate introgressed loci. *Mol. Biol. Evol.* **32,** 244–257 (2014).
127. Zheng, Y. & Janke, A. Gene flow analysis method, the D-statistic, is robust in a wide

parameter space. *BMC Bioinformatics* **19,** 10 (2018).

128. Fukumoto, S., Ushimaru, A. & Minamoto, T. A basin-scale application of environmental DNA assessment for rare endemic species and closely related exotic species in rivers: a case study of giant salamanders in Japan. *J. Appl. Ecol.* **52,** 358–365 (2015).

129. Malyarchuk, B. A., Derenko, M. V & Denisova, G. A. Phylogenetic relationships among Asiatic salamanders of the genus *Salamandrella* based on variability of nuclear genes. *Russ. J. Genet.* **51,** 91–97 (2015).

130. Kawamura, T. Interspecific hybrids between *Hynobius nigrescens* and *Hynobius tokyoensis*. *J. Fac. Sci. Hokkaido Univ.* **13,** 248–252 (1957).

131. Steele, C. A., Baumsteiger, J. & Storfer, A. Polymorphic tetranucleotide microsatellites for Cope's giant salamander (*Dicamptodon copei*) and Pacific giant salamander (*Dicamptodon tenebrosus*). *Mol. Ecol. Resour.* **8,** 1071–1073 (2008).

132. Bogart, J. P., Bi, K., Fu, J., Noble, D. W. A. & Niedzwiecki, J. Unisexual salamanders (genus *Ambystoma*) present a new reproductive mode for eukaryotes. *Genome* **50,** 119–136 (2007).

133. Eastman, J. M., Niedzwiecki, J. N., Nadler, B. P. & Storfer, A. Duration and consistency of historical selection are correlated with adaptive trait evolution in the streamside salamander, *Ambystoma barbouri*. *Evolution.* **43,** 468–479 (2009).

134. Fitzpatrick, B. M. & Shaffer, H. B. Hybrid vigor between native and introduced salamanders raises new challenges for conservation. *Proc. Natl. Acad. Sci.* **104,** 15793–15798 (2007).

135. Weisrock, D. W., Shaffer, H. B., Storz, B. L., Storz, S. R. & Voss, S. R. Multiple nuclear gene sequences identify phylogenetic species boundaries in the rapidly radiating clade of Mexican ambystomatid salamanders. *Mol. Ecol.* **15,** 2489–2503 (2006).

136. Canestrelli, D., Bisconti, R. & Nascetti, G. Extensive unidirectional introgression between two salamander lineages of ancient divergence and its evolutionary implications. *Sci. Rep.* **4,** 6516 (2014).

137. Hauswaldt, J. S., Angelini, C., Pollok, A. & Steinfartz, S. Hybridization of two ancient salamander lineages: molecular evidence for endemic spectacled salamanders on the Apennine peninsula. *J. Zool.* **284,** 248–256 (2011).

138. Veith M., *et al.* Cracking the nut: Geographical adjacency of sister taxa supports vicariance in a polytomic salamander clade in the absence of node support. *Mol. Phylogenet. Evol.* **47,** 916–931 (2008).

139. Johannesen, J. *et al.* Distortion of symmetrical introgression in a hybrid zone: evidence for locus-specific selection and uni-directional range expansion. *J. Evol. Biol.* **19,** 705–716 (2006).

140. Escoriza, D., Gutiérrez-Rodríguez, J., Hassine, J. B. & Martínez-Solano, I. Genetic assessment of the threatened microendemic *Pleurodeles poireti* (Caudata, Salamandridae), with molecular evidence for hybridization with *Pleurodeles* nebulosus. *Conserv. Genet.* **17,** 1445–1458 (2016).

141. Phimmachak, S., Aowphol, A. & Stuart, B. L. Morphological and molecular variation in *Tylototriton* (Caudata: Salamandridae) in Laos, with description of a new species. *Zootaxa* **4006,** 285–310 (2015).

142. Davis, W. C. & Twitty, V. C. Courtship behavior and reproductive isolation in the species of *Taricha* (Amphibia, Caudata). *Copeia* **1964,** 601–610 (1964).
143. Hedgecock, D. & Ayala, F. J. Evolutionary divergence in the genus *Taricha* (Salamandridae). *Copeia* **1974,** 738–747 (1974).
144. Kuchta, S. R. & Tan, A. M. Isolation by distance and post-glacial range expansion in the rough-skinned newt, *Taricha granulosa*. *Mol. Ecol.* **14,** 225–244 (2005).
145. Kuchta, S. R. & Tan, A. M. Limited genetic variation across the range of the red-bellied newt, *Taricha rivularis*. *J. Herpetol.* **40,** 561–565 (2006).
146. Kuchta, S. R. Contact zones and species limits: hybridization between lineages of the California Newt, *Taricha torosa* in the southern Sierra Nevada. *Herpetologica* **63,** 332–350 (2007).
147. Johanet, A., Secondi, J. & Lemaire, C. Widespread introgression does not leak into allotopy in a broad sympatric zone. *Heredity.* **106,** 962 (2011).
148. Babik, W. *et al.* Phylogeography of two European newt species -- discordance between mtDNA and morphology. *Mol. Ecol.* **14,** 2475–2491 (2005).
149. van Riemsdijk, I. *et al.* The Near East as a cradle of biodiversity: a phylogeography of banded newts (genus *Ommatotriton*) reveals extensive inter-and intraspecific genetic differentiation. *Mol. Phylogenet. Evol.* **114,** 73–81 (2017).
150. Themudo, G., Nieman, A. M. & Arntzen, J. W. Is dispersal guided by the environment? A comparison of interspecific gene flow estimates among differentiated regions of a newt hybrid zone. *Mol. Ecol.* **21,** 5324–5335 (2012).
151. Schoorl, J. & Zuiderwijk, A. Ecological isolation in *Triturus cristatus* and *Triturus marmoratus* (Amphibia: Salamandridae). *Amphibia-Reptilia* **1,** 235–252 (1981).
152. Brede, E. G., Thorpe, R. S., Arntzen, J. W. & Langton, T. E. S. A morphometric study of a hybrid newt population (*Triturus cristatus*/*T. carnifex*): Beam Brook Nurseries. Surrey, UK. *Biol. J. of the Linn. Soc.* **70,** 685-295 (2000).
153. Wielstra, B. & Arntzen, J. W. Postglacial species displacement in *Triturus* newts deduced from asymmetrically introgressed mitochondrial DNA and ecological niche models. *BMC Evol. Biol.* **12,** 161 (2012).
154. Nelson, S. K., Niemiller, M. L. & Fitzpatrick, B. M. Co-occurrence and hybridization between *Necturus maculosus* and a heretofore unknown *Necturus* in the Southern Appalachians. *J. Herpetol.* **51,** 559–566 (2017).
155. Bonett, R. M., Chippindale, P. T., Moler, P. E., Van Devender, R. W. & Wake, D. B. Evolution of gigantism in amphiumid salamanders. *PLoSONE* **4,** (2009).
156. Nascetti, G., Cimmaruta, R., Lanza, B. & Bullini, L. Molecular taxonomy of European plethodontid salamanders (genus *Hydromantes*). *J. Herpetol.* **30,** 161–183 (1996).
157. Tilley, S. G. Review of: Morphologic and Genetic Studies of the European Plethodontid Salamanders: Taxonomic Inferences (Genus *Hydromantes*) by Benedetto Lanza, Vincenzo Caputo, Giuseppe Nascetti, and Luciano Bullini. *Copeia* **1998,** 526–528 (1998).
158. Jackman, T. R. Molecular and historical evidence for the introduction of clouded salamanders (genus *Aneides*) to Vancouver Island, British Columbia, Canada, from California. *Can. J. Zool.* **76,** 1570–1580 (1998).

159. Kozak, K. H. Sexual isolation and courtship behavior in salamanders of the *Eurycea bislineata* species complex, with comments on the evolution of the mental gland and pheromone delivery behavior in the Plethodontidae. *Southeast. Nat.* **2,** 281–292 (2003).
160. Bonett, R. M. Analysis of the contact zone between the dusky salamanders *Desmognathus fuscus fuscus* and *Desmognathus fuscus conanti* (Caudata: Plethodontidae). *Copeia* **105,** 344–355 (2002).
161. Highton, R., *et al.* Concurrent speciation in the eastern woodland salamanders (genus Plethodon): DNA sequences of the complete albumin nuclear and partial mitochondrial 12s genes. *Mol. Phyl. and Evol.* **63**, 278-290. (2012).
162. Highton, R. Geographic protein variation and speciation in the salamanders of the *Plethodon cinereus* group with the description of two new species. *Herpetologica* **1999,** 43–90 (1999).
163. Highton, R. Geographic protein variation and speciation in the *Plethodon* dorsalis complex. *Herpetologica* **1997,** 345–356 (1997).
164. Highton, R. Speciation in eastern North American salamanders of the genus *Plethodon. Annu. Rev. Ecol. Syst.* **26,** 579–600 (1995).
165. Highton, R. & Peabody, R. in *The Biology of Plethodontid Salamanders.* (eds. Bruce, R. C., Jaeger, R. G. & Houck, L. D.) 31–93 (Kluwer Academic/Plenum Publishers, 2000).
166. Niemiller, M. L., Fitzpatrick, B. M. & Miller, B. T. Recent divergence with gene flow in Tennessee cave salamanders (Plethodontidae: *Gyrinophilus*) inferred from gene genealogies. *Mol. Ecol.* **17,** 2258–2275 (2008).
167. Kuchta, S. R., Haughey, M., Wynn, A. H., Jacobs, J. F. & Highton, R. Ancient river systems and phylogeographical structure in the spring salamander, *Gyrinophilus porphyriticus*. *J. Biogeogr.* **43,** 639–652 (2016).
168. Smith, C. C. Problem of the Hybridization of the Red Cave Salamander, *Eurycea lucifuga* (Raf), and the Long-Tailed Salamander, *Eurycea longicauda melanopleura* (Green). *J. Ark. Acad. Sci.* **18,** 59–62 (1964).
169. Sweet, S. S. Secondary contact and hybridisation in the Texas cave salamanders *Eurycea neotenes* and *E. tridentifera*. *Copeia* **1984,** 428–441 (1984).
170. Guttman, S. I. & Karlin, A. A. Hybridisation of cryptic species of two-lined salamanders. (*Eurycea bislineata* complex). *Copeia* **1986,** 96–108 (1986).
171. Kozak, K. H. & Montanucci, R. R. Genetic variation across a contact zone between montane and lowland forms of the Two-lined salamander (*Eurycea bislineata*) species complex: a test of species limits. *Copeia* **2001,** 25–34 (2001).
172. Bendik, N. F., Meik, J. M., Gluesenkamp, A. G., Roelke, C. E. & Chippindale, P. T. Biogeography, phylogeny, and morphological evolution of central Texas cave and spring salamanders. *BMC Evol. Biol.* **13,** 201 (2013).
173. Jockusch, E. L., Martinez-Solano, I., Hansen, R. W. & Wake, D. B. Morphological and molecular diversification of slender salamanders (Caudata: Plethodontidae: *Batrachoseps*) in the southern Sierra Nevada of California with descriptions of two new species. *Zootaxa* **3190,** 30 (2012).
174. Jockusch, E. L. & Wake, D. B. Falling apart and merging: diversification of slender salamanders (Plethodontidae: *Batrachoseps*) in the American West. *Biol. J. Linn. Soc.* **76,** 361–391 (2002).

175. Jockusch, E. L., Yanev, K. P. and Wake, D. B. Molecular phylogenetic analysis of slender salamanders, genus Batrachoseps (Amphibia: *Plethodontidae*), from central coastal California with descriptions of four new species. *Herp. Mono.* 54-99. (2001).
176. Wake, D. B. & Lynch, J. F. Evolutionary relationships among Central American salamanders of the *Bolitoglossa franklini* group, with a description of a new species from Guatemala. *Herpetologica* **38,** 257–272 (1982).
177. Wake, D. B., Yang, S. Y. & Papenfuss, T. J. Natural hybridization and its evolutionary implications in Guatemalan plethodontid salamanders of the genus *Bolitoglossa*. *Herpetologica* **1980,** 335–345 (1980).

#### ***Supplementary Figure/Table Captions***

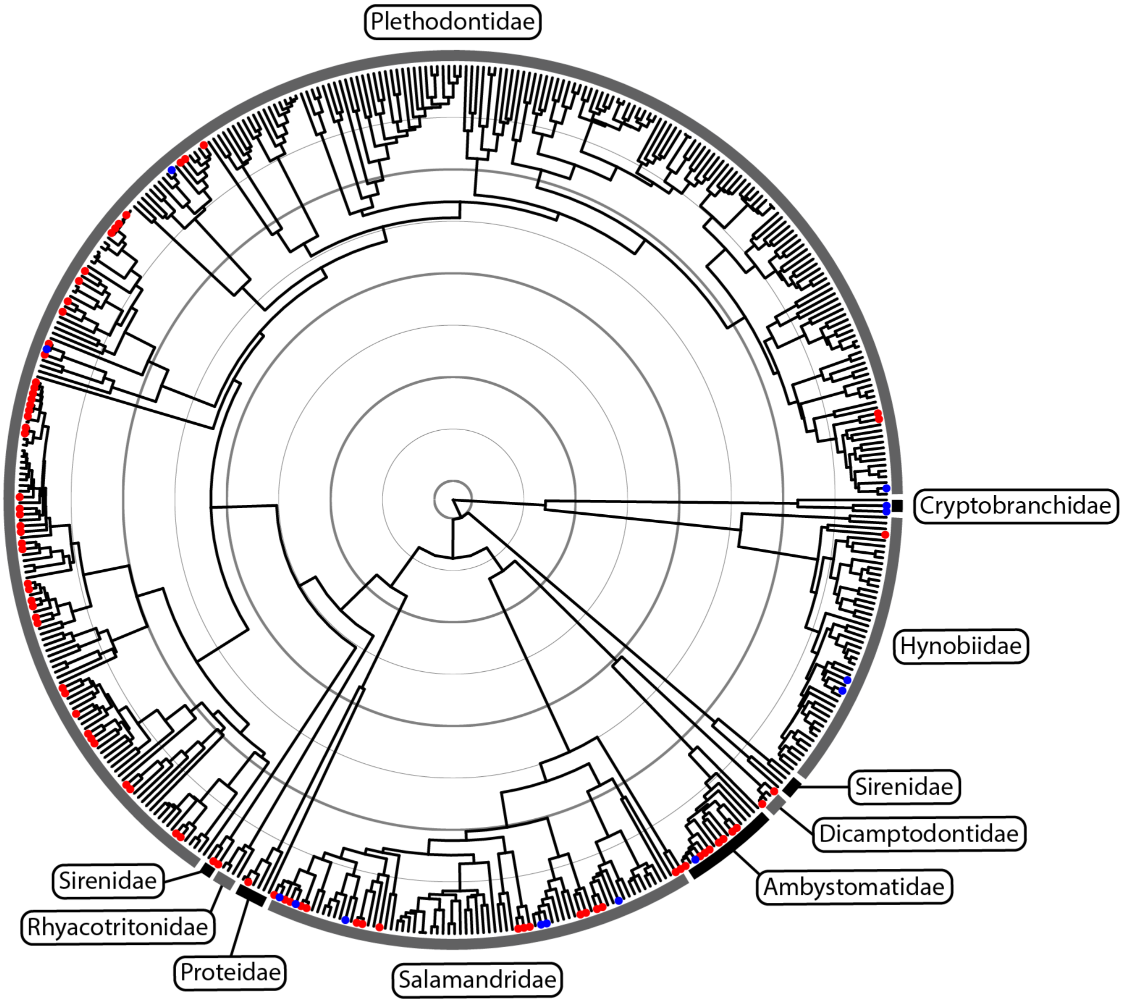


**Figure S1. Phylogeny used in analysis of complete dataset.** Salamanders were pruned from the time calibrated tree of Pyron (2014). Species observed to hybridize in nature according to the narrow dataset have red circles at the tips. Additional species included in the broad dataset are indicated by blue circles. Families are shown according to the external broken circle, each labeled with their respective names. Each internal circle corresponds to 25 million years.

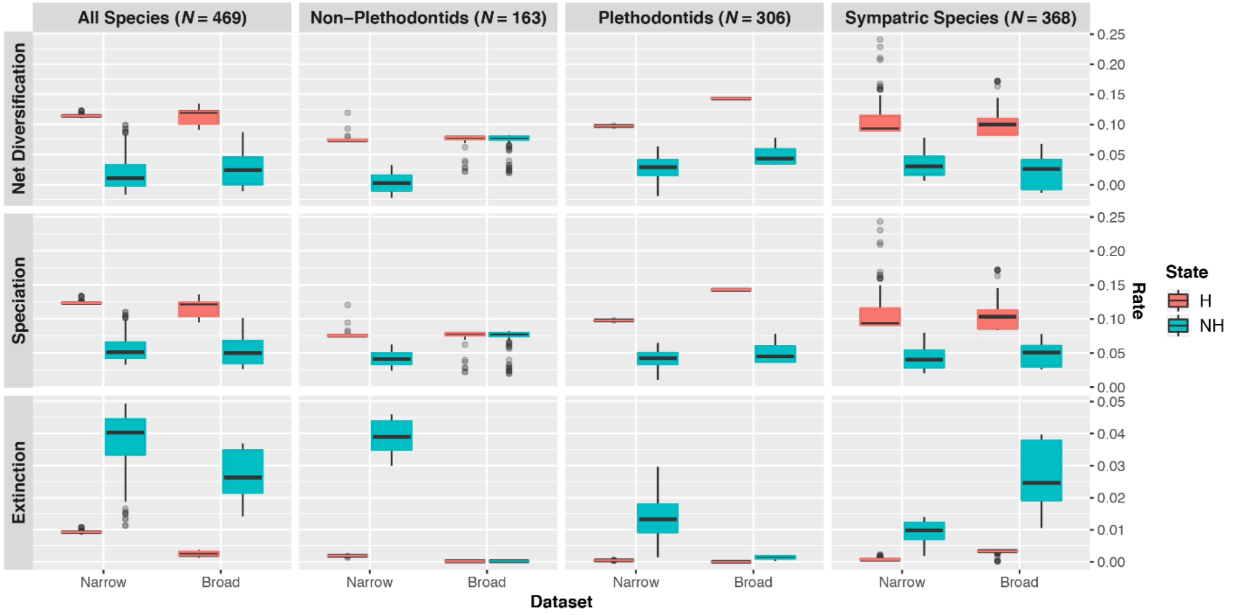


**Figure S2. Model-averaged lineage-specific diversification rate estimates at the tips of the phylogeny assuming we have sampled all extant hybridizing salamanders.** Results using different trees are displayed by column, whereas results for different parameters are displayed by row. Hybridizing lineages (H) are displayed in red, whereas non-hybridizing (NH) lineages are displayed in blue.

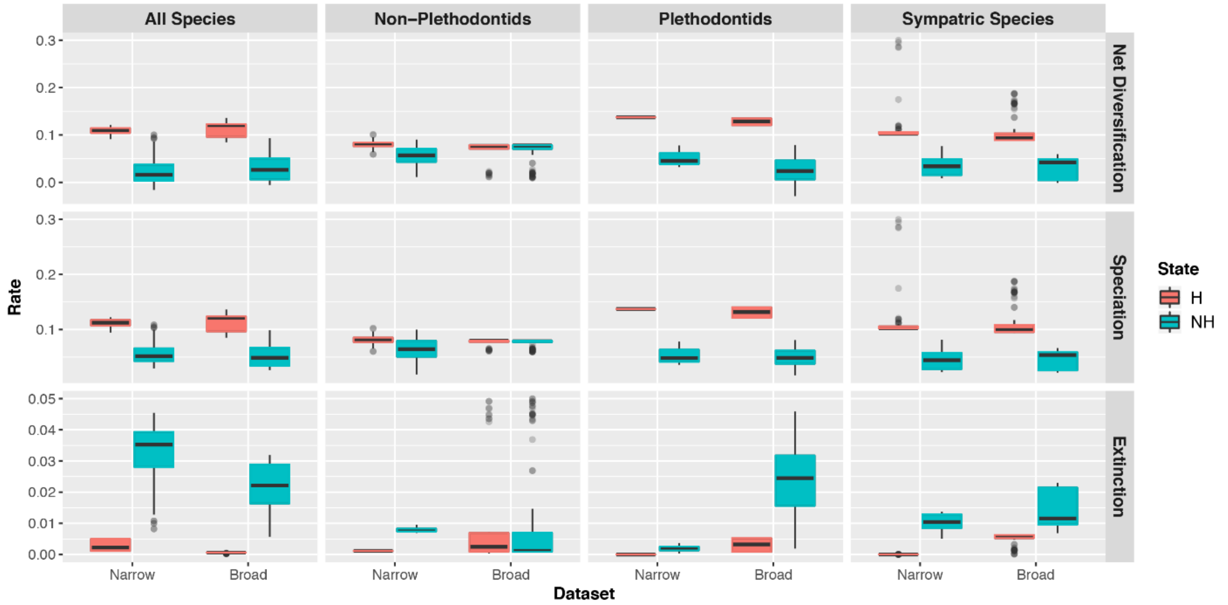


**Figure S3. Model-averaged lineage-specific diversification rate estimates at the tips of the phylogeny assuming the frequency of hybridization is equal to that observed in our data.** Results using different trees are displayed by column, whereas results for different parameters are displayed by row. Hybridizing lineages (H) are displayed in red, whereas non-hybridizing (NH) lineages are displayed in blue.

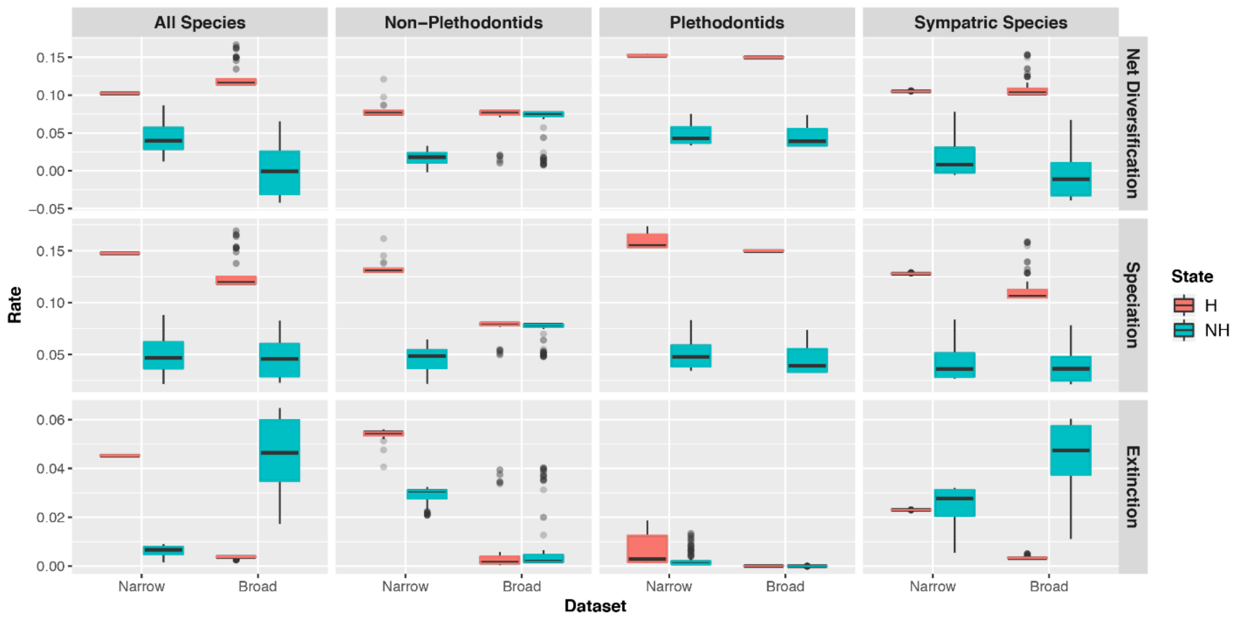


**Figure S4. Model-averaged lineage-specific diversification rate estimates at the tips of the phylogeny assuming 30% of species hybridize.** Results using different trees are displayed by column, whereas results for different parameters are displayed by row. Hybridizing lineages (H) are displayed in red, whereas non-hybridizing (NH) lineages are displayed in blue.

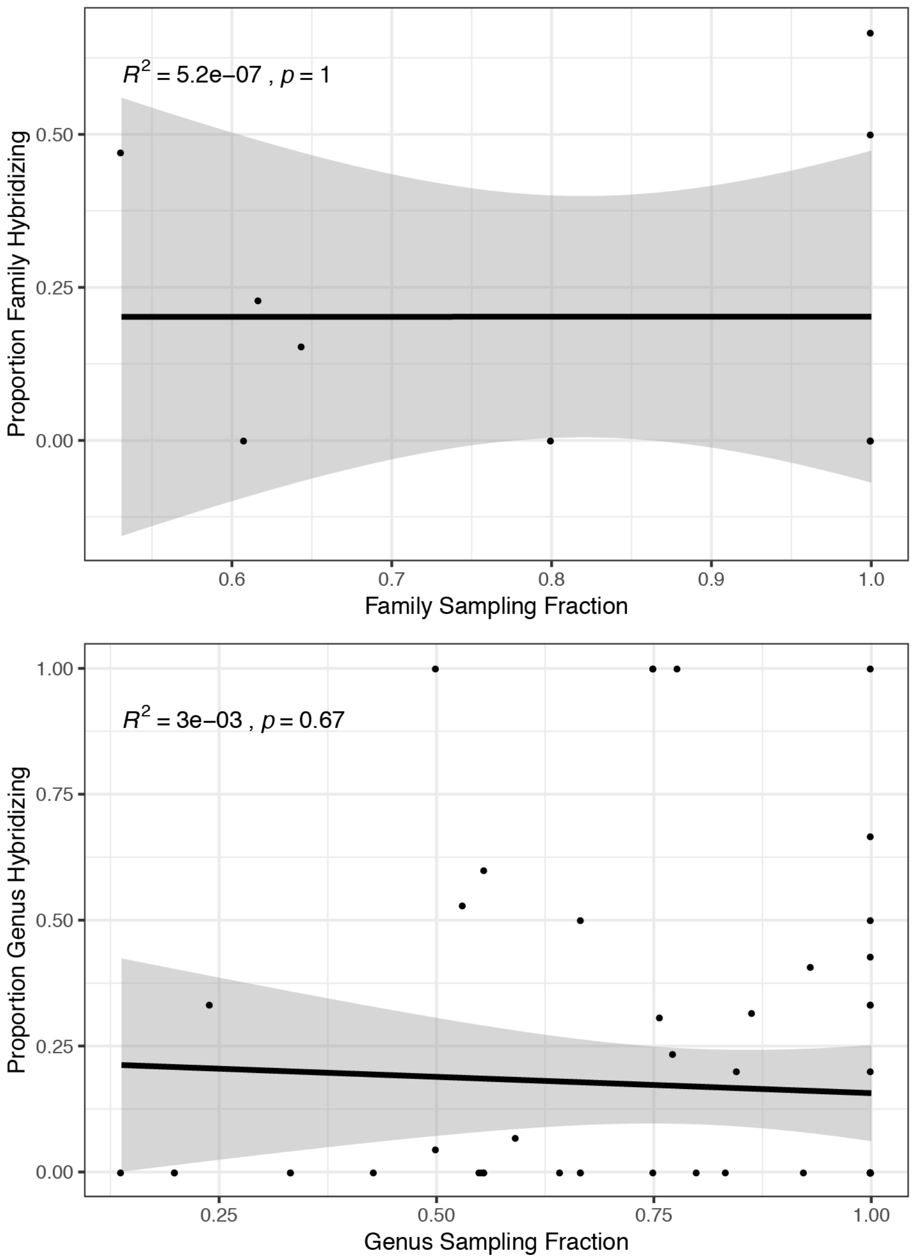


**Figure S5. Relationship between clade-specific sampling fraction and prevalence of hybridization.** The top plot shows results at the family level, whereas the bottom plot shows results at the genus level. No relationship is found for either dataset. Sampling fraction was defined as the proportion of extant diversity for each family/genus sampled in our phylogeny. Proportion family/genus hybridizing was defined as the proportion of extant family/genus diversity observed to hybridize in our dataset.

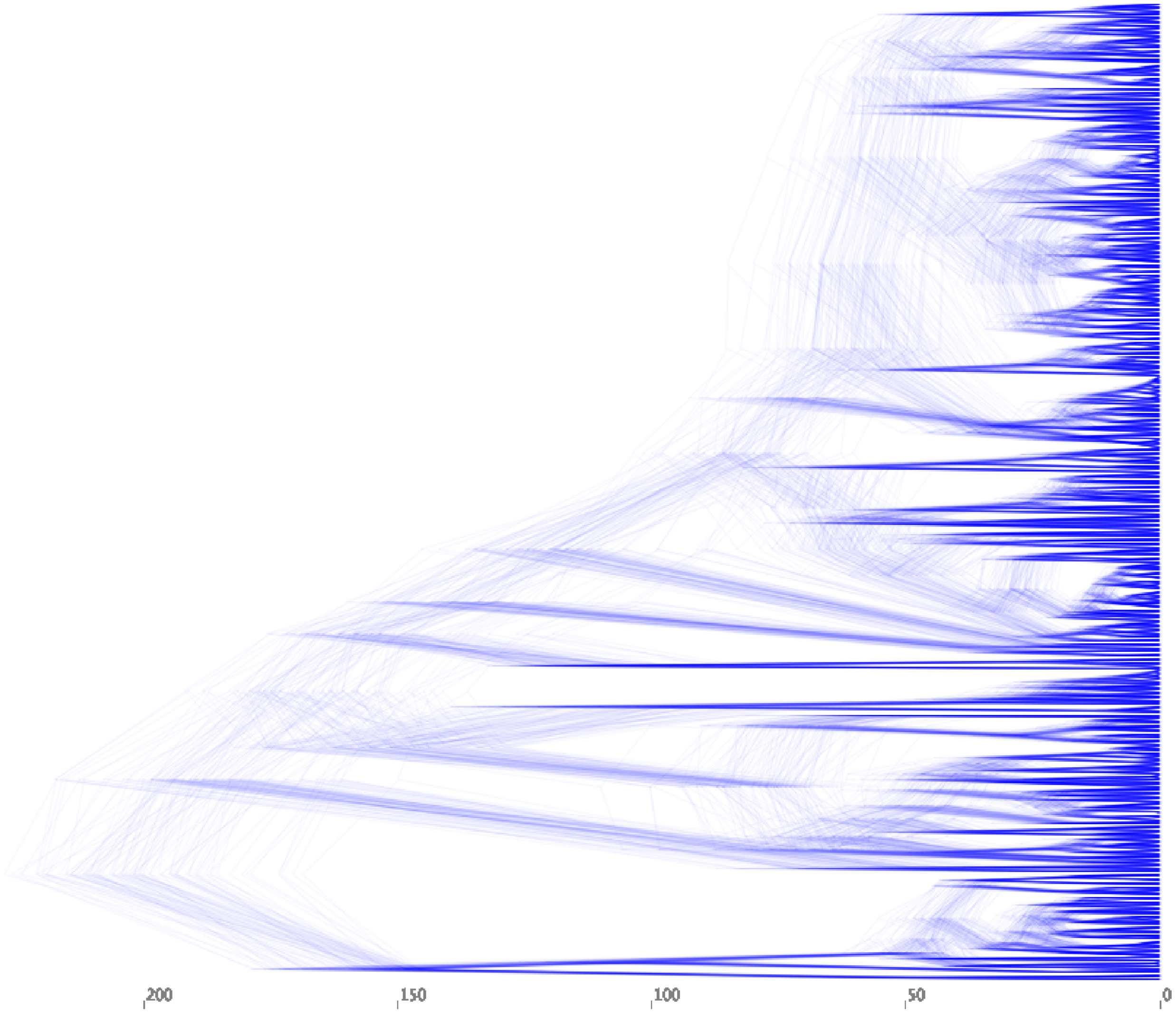


**Figure S6. Posterior distribution of 100 trees used to assess phylogenetic uncertainty.** Trees are obtained from Jetz and Pyron (2018).

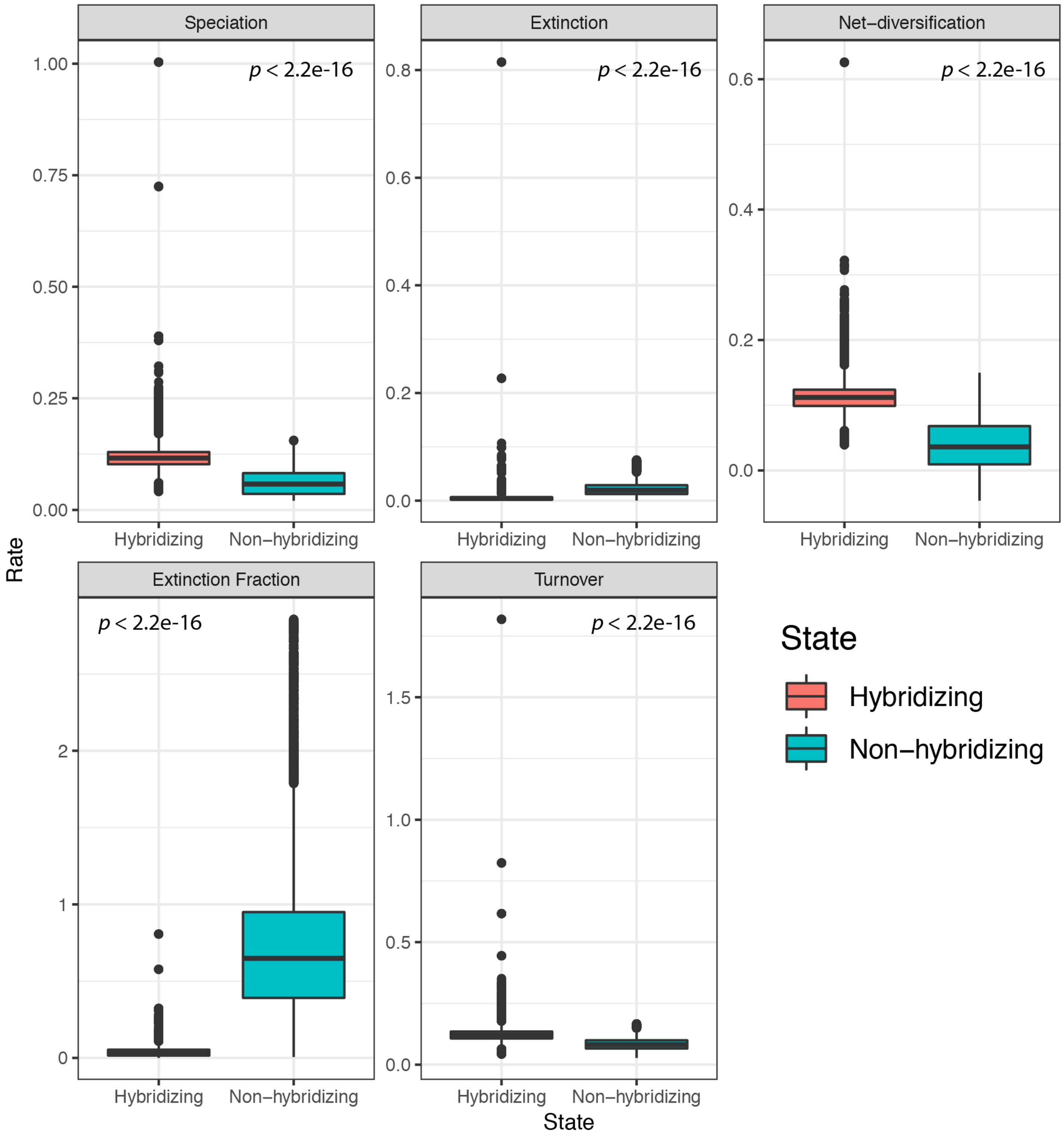


**Figure S7. Diversification rates obtained by analysis of the 100 trees obtained from Jetz and Pyron (2018).** Hybridizing lineages (H) are displayed in red, whereas non-hybridizing (NH) lineages are displayed in blue. P-values correspond to those obtained using a Welch's two-sample t-test using pooled model-averaged diversification rates across replicates.

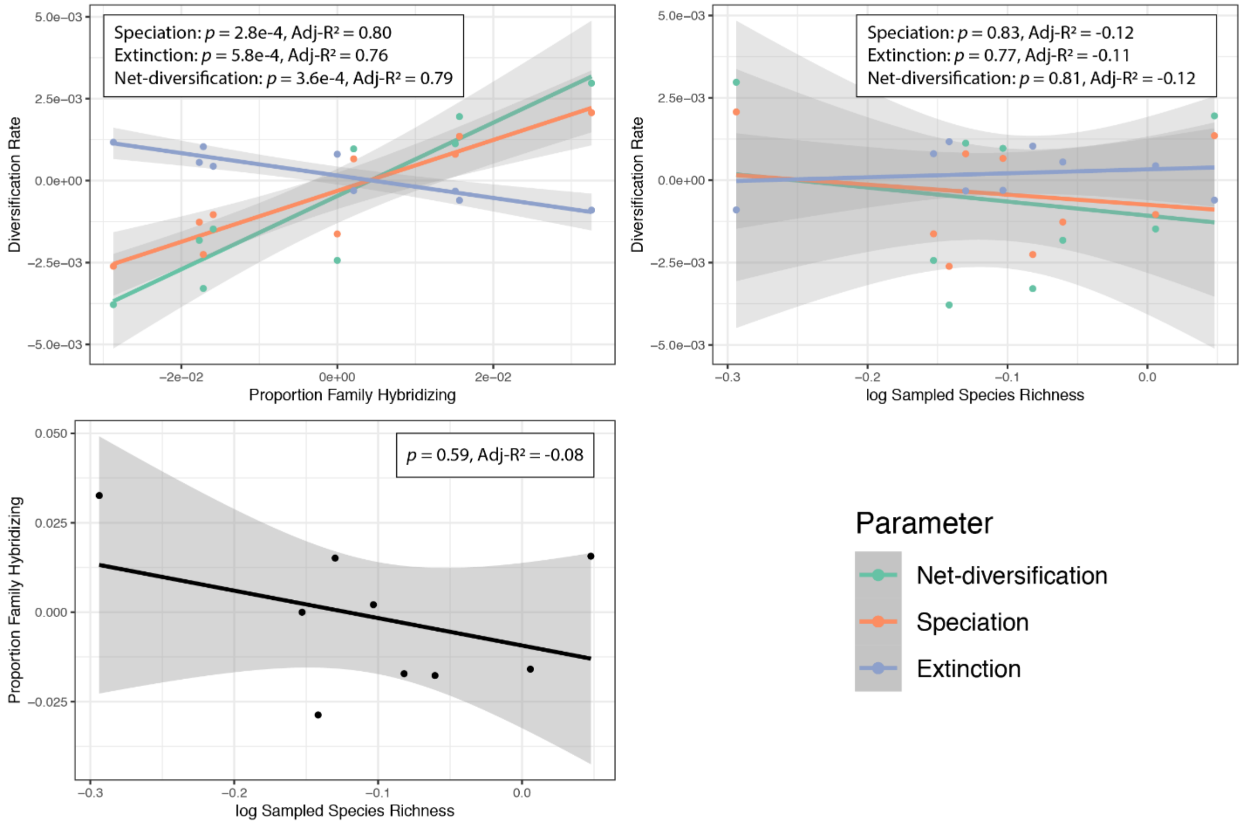


**Figure S8. Relationships between A) family-level diversification rate and proportion family hybridizing, B) family-level diversification rate and species richness, and C) proportion of family hybridizing and species richness.** Regressions and points are colored by rate (see inset legends).

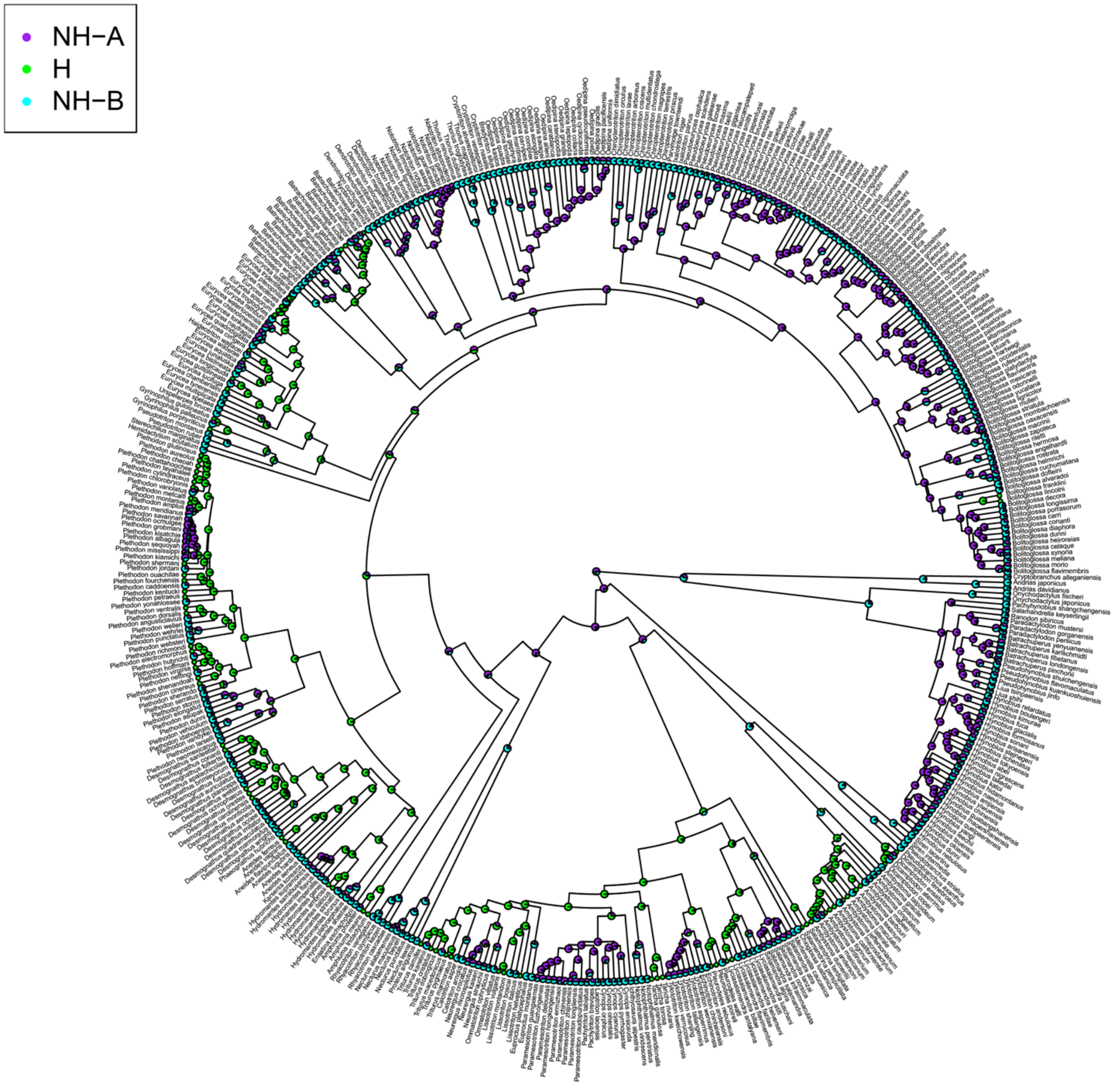


**Figure S9. Marginal ancestral reconstruction of character states according to the best-fit model as implemented in HiSSE using the dataset including all taxa.** Tip and node states represent their respective probabilities of belonging to each of the four states included in the full HiSSE model. The two hidden states are indicated by (**A**) and (**B**) respectively.

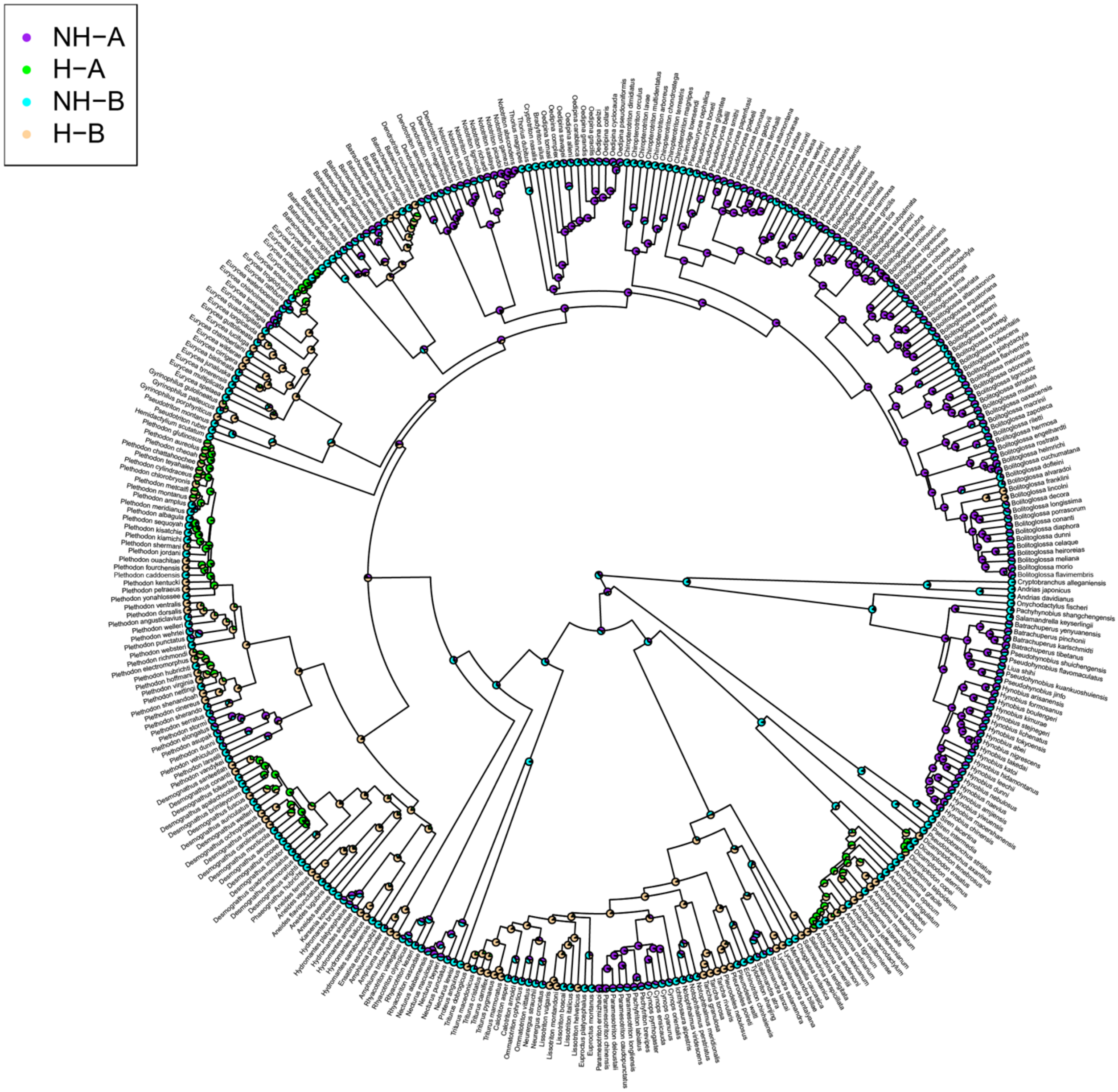


**Figure S10. Marginal ancestral reconstruction of character states according to the best-fit model as implemented in HiSSE using the dataset including only sympatric taxa.** Tip and node states represent their respective probabilities of belonging to each of the four states included in the full HiSSE model. The two hidden states are indicated by (**A**) and (**B**) respectively.

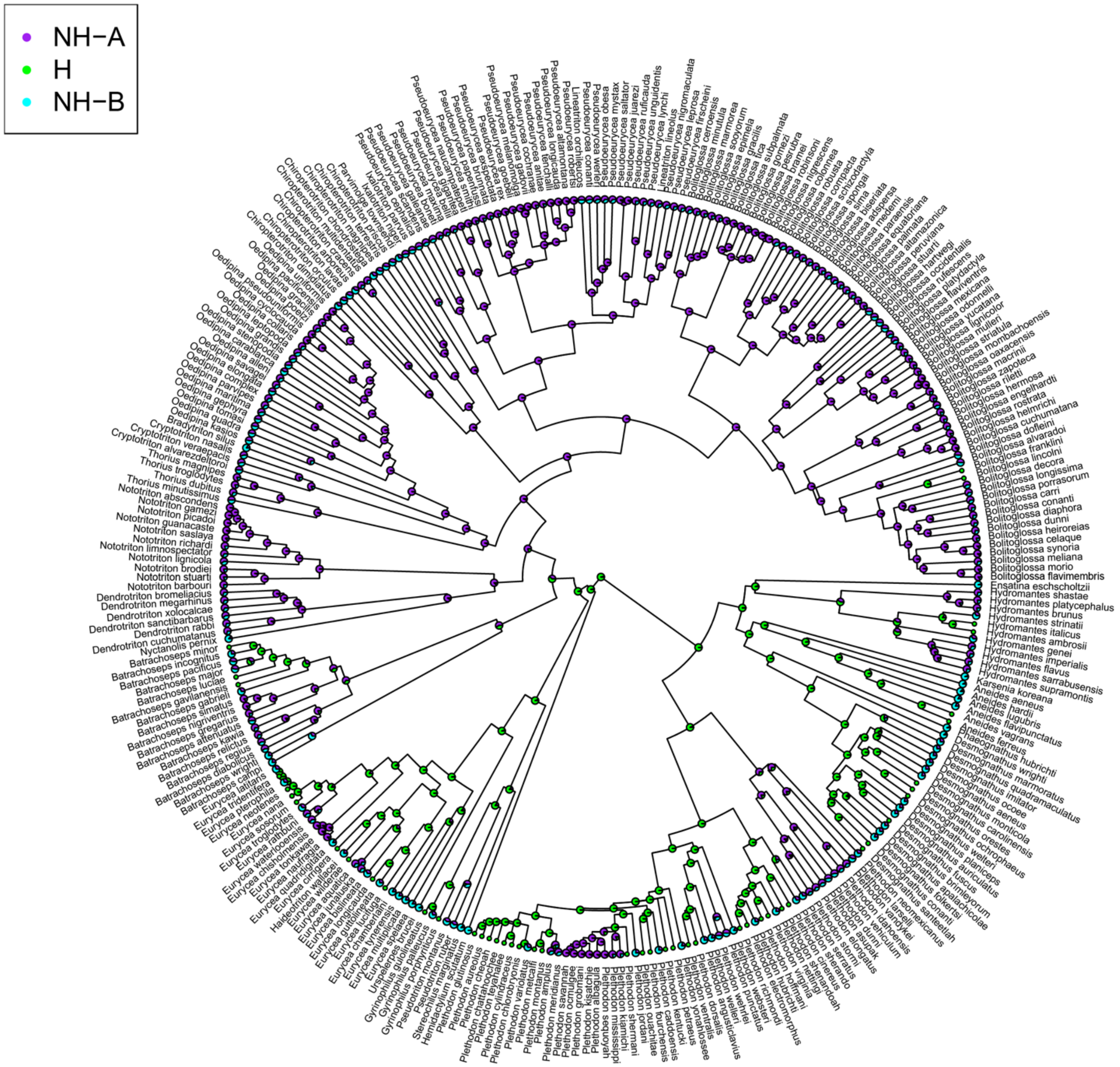


**Figure S11. Marginal ancestral reconstruction of character states according to the best-fit model as implemented in HiSSE using the dataset including only plethodontids.** Tip and node states represent their respective probabilities of belonging to each of the four states included in the full HiSSE model. The two hidden states are indicated by (**A**) and (**B**) respectively.

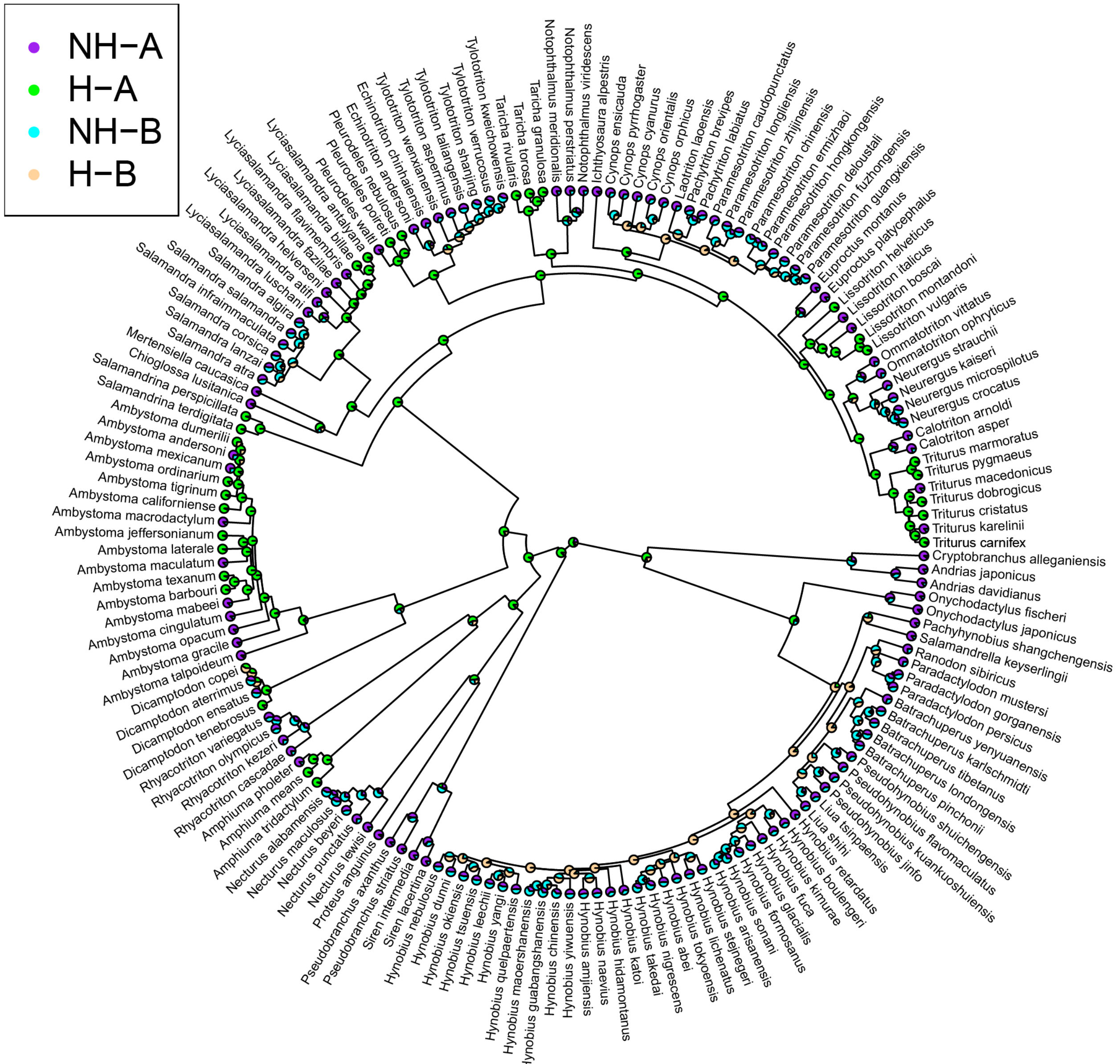


**Figure S12. Marginal ancestral reconstruction of character states according to the best-fit model as implemented in HiSSE using the dataset excluding plethodontids.** Tip and node states represent their respective probabilities of belonging to each of the four states included in the full HiSSE model. The two hidden states are indicated by (**A**) and (**B**) respectively.

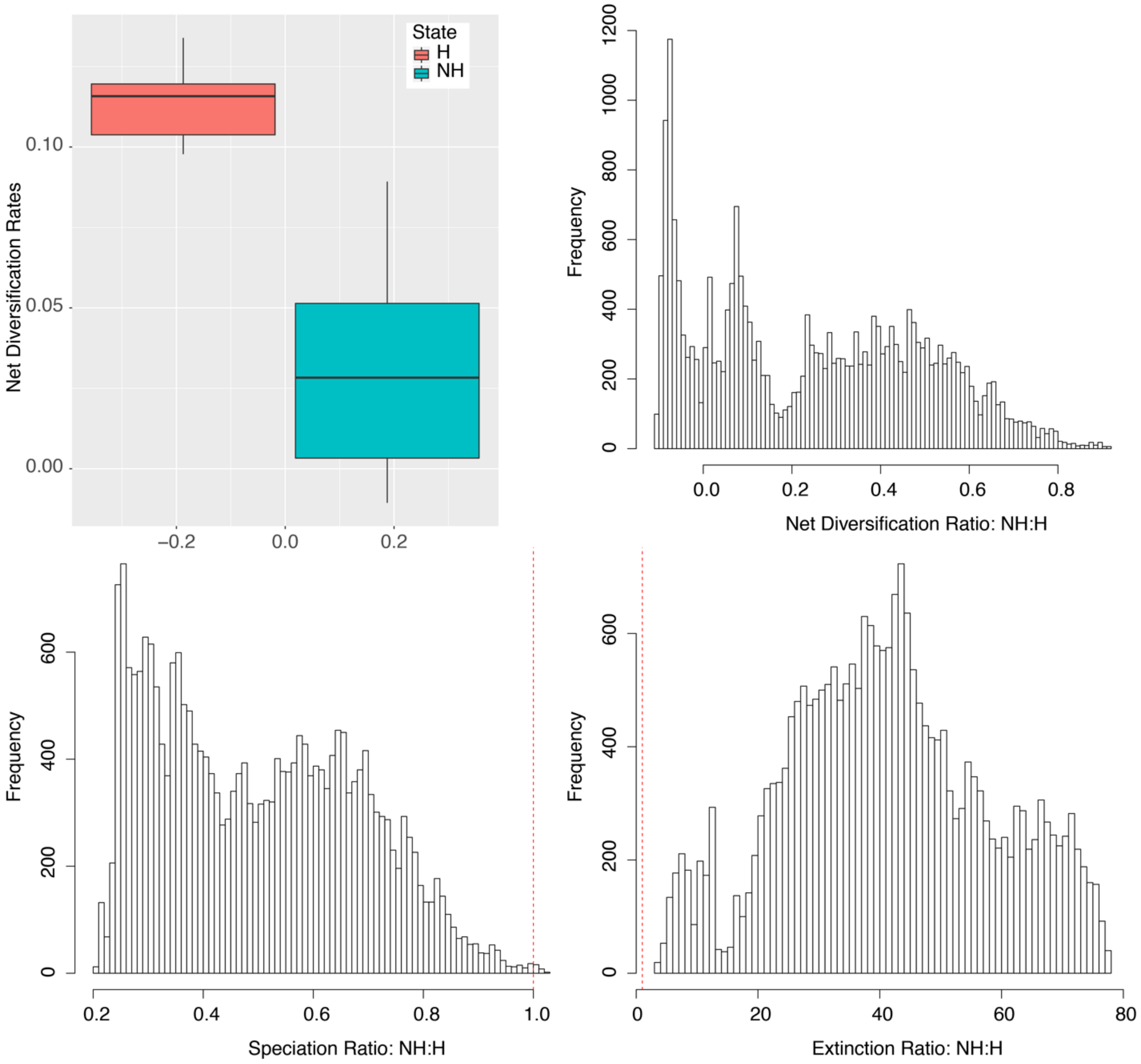


**Figure S13. Comparison of model averaged parameter estimates among character states excluding the *Glutinosus* group within genus *Plethodon* and using the narrow definition of hybridization.** Results reported here are those obtained when, assuming 20% of species hybridize. The box plots in the top left show model averaged net diversification rates for each state at the tips of the tree. The histograms illustrate the distributions of non-hybridizing to hybridizing lineages diversification rates as estimated at the tips of the phylogeny. A value > 1 corresponds to a comparison in which non-hybridizing lineages experience rates greater than those of hybridizing lineages and vice-versa. Dotted vertical lines are placed at 1, at which rates are equal among states.

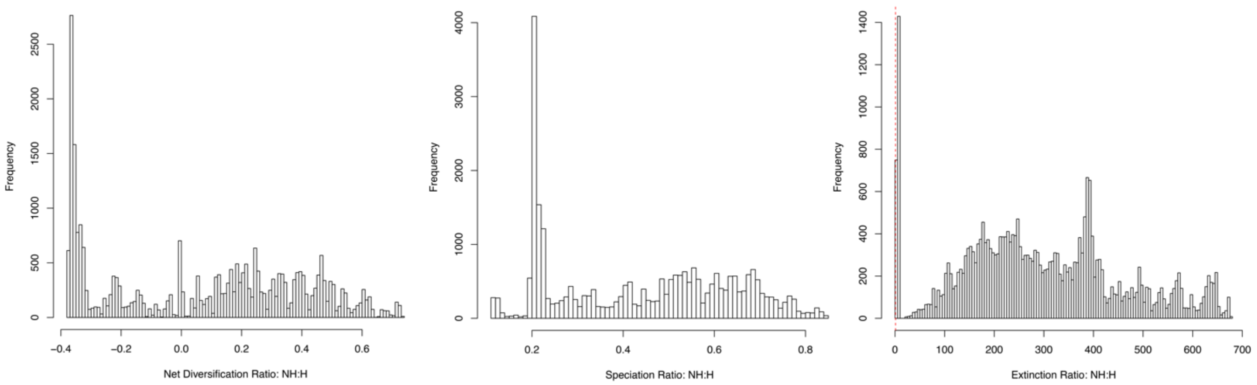


**Figure S13. Comparison of model averaged parameter estimates among character states excluding the *Glutinosus* group within genus *Plethodon* and using the broad definition of hybridization.** Results reported here are those obtained when, assuming 20% of species hybridize. Histograms illustrate the distributions of non-hybridizing to hybridizing lineages diversification rates as estimated at the tips of the phylogeny. A value > 1 corresponds to a comparison in which non-hybridizing lineages experience rates greater than those of hybridizing lineages and vice-versa. Dotted vertical lines are placed at 1, at which rates are equal among states.

| Cryptobranchidae | Narrow | Broad | Reference |
|---|---|---|---|
| *Andrias davidianus* | 0 | 1 | Fukumoto et al. 2015 |
| *Andrias japonicus* | 0 | 1 | Fukumoto et al. 2015 |
| **Hynobiidae** | | | |
| *Salamandrella keyserlingii* | 1 | 1 | Malyarchuk et al. 2015 |
| *Hynobius tokyoensis* | 0 | 1 | Kawamura & Toshijiro 1957 |
| *Hynobius nigrescens* | 0 | 1 | Kawamura & Toshijiro 1957 |
| **Dicamptodontidae** | | | |
| *Dicamptodon tenebrosus* | 1 | 1 | Steele et al. 2008 |
| *Dicamptodon copei* | 1 | 1 | Steele et al. 2008 |
| **Ambystomatidae** | | | |
| *Ambystoma jeffersonianum* | 1 | 1 | Bogart et al. 2007 |
| *Ambystoma laterale* | 1 | 1 | Bogart et al. 2007 |
| *Ambystoma texanum* | 1 | 1 | Eastman et al. 2009 |
| *Ambystoma barbouri* | 1 | 1 | Bogart et al. 2007; Eastman et al. 2009 |
| *Ambystoma californiense* | 1 | 1 | Fitzpatrick and Shaffer 2007 |
| *Ambystoma tigrinum* | 1 | 1 | Bogart et al. 2007 |
| *Ambystoma ordinarium* | 1 | 1 | Weisrock et al. 2006 |
| *Ambystoma mexicanum* | 0 | 1 | Brandon 1972 |
| *Ambystoma dumerilii* | 1 | 1 | Weisrock et al. 2006 |
| **Salamandridae** | | | |
| *Salamandrina perspicillata* | 1 | 1 | Canestrelli et al. 2014; Hauswaldt et al. 2011 |
| *Salamandrina terdigitata* | 1 | 1 | Canestrelli et al. 2014; Hauswaldt et al. 2011 |
| *Lyciasalamandra helverseni* | 0 | 1 | Veith et al. 2008 |
| *Lyciasalamandra antalyana* | 1 | 1 | Johannesen et al. 2006 |
| *Lyciasalamandra billae* | 1 | 1 | Johannesen et al. 2006 |
| *Pleurodeles nebulosus* | 1 | 1 | Escoriza et al. 2016 |
| *Pleurodeles poireti* | 1 | 1 | Escoriza et al. 2016 |
| *Tylototriton shanjing* | 0 | 1 | Phimmachak et al. 2015 |
| *Tylototriton verrucosus* | 0 | 1 | Phimmachak et al. 2015 |
| *Taricha rivularis* | 1 | 1 | Davis and Twitty 1964; Hedgecock and Ayala 1974; Kuchta and Tan 2005, 2006; Kuchta 2007 |
| *Taricha granulosa* | 1 | 1 | Davis and Twitty 1964; Hedgecock and Ayala 1974; Kuchta and Tan 2005, 2006; Kuchta 2007 |
| *Taricha torosa* | 1 | 1 | Davis and Twitty 1964; Hedgecock and Ayala 1974; Kuchta and Tan 2005, 2006; Kuchta 2007 |
| *Lissotriton helveticus* | 1 | 1 | Johanet et al. 2011 |
| *Lissotriton vulgaris* | 1 | 1 | Babik et al. 2005; Steinfartz et al. 2007 |
| *Lissotriton montandoni* | 1 | 1 | Babik et al. 2005; Steinfartz et al. 2007 |
| *Ommatotriton ophryticus* | 0 | 1 | van Riemsdijk et al. 2017 |
| *Triturus pygmaeus* | 1 | 1 | Themudo et al. 2012 |
| *Triturus marmoratus* | 1 | 1 | Schoorl and Zuiderwijk 1981; Brede et al. 2000; Babik et al. 2005 |
| *Triturus dobrogicus* | 1 | 1 | Schoorl and Zuiderwijk 1981; Brede et al. 2000; Babik et al. 2005 |
| *Triturus macedonicus* | 0 | 1 | Wielstra and Arntzen 2012 |
| *Triturus cristatus* | 1 | 1 | Schoorl and Zuiderwijk 1981; Brede et al. 2000; Babik et al. 2005 |
| *Triturus carnifex* | 1 | 1 | Schoorl and Zuiderwijk 1981; Brede et al. 2000; Babik et al. 2005 |
| *Triturus karelinii* | 0 | 1 | Wielstra and Arntzen 2012 |
| **Proteidae** | | | |
| *Necturus maculosus* | 0 | 1 | Nelson et al. 2017; Hybridization between N. maculosus and a new form (not yet described species) that is related to/sister to N. lewisi and N.punctatus. |
| **Amphiumidae** | | | |
| *Amphiuma tridactylum* | 1 | 1 | Bonett et al. 2009 |
| *Amphiuma means* | 1 | 1 | Bonett et al. 2009 |
| **Plethodontidae** | | | |
| *Hydromantes ambrosii* | 1 | 1 | Nascetti et al. 1996; Tilley 1998 |
| *Hydromantes italicus* | 1 | 1 | Nascetti et al. 1996; Tilley 1998 |
| *Aneides ferreus* | 1 | 1 | Jackman 1998 |
| *Aneides vagrans* | 1 | 1 | Jackman 1998 |
| *Desmognathus ochrophaeus* | 1 | 1 | Kozak 2003; Bonett 2002 |
| *Desmognathus orestes* | 1 | 1 | Kozak 2003; Bonett 2002 |
| *Desmognathus carolinensis* | 1 | 1 | Kozak 2003; Bonett 2002 |
| *Desmognathus fuscus* | 1 | 1 | Kozak 2003; Bonett 2002 |
| *Desmognathus santeetlah* | 1 | 1 | Kozak 2003; Bonett 2002 |
| *Desmognathus conanti* | 1 | 1 | Kozak 2003; Bonett 2002 |
| *Plethodon shenandoah* | 1 | 1 | Highton 2012 |
| *Plethodon cinereus* | 1 | 1 | Highton 1999 |
| *Plethodon hoffmani* | 1 | 1 | Highton 1999 |
| *Plethodon virginia* | 1 | 1 | Highton 1999 |
| *Plethodon richmondi* | 1 | 1 | Highton 1999 |
| *Plethodon electromorphus* | 1 | 1 | Highton 1999 |
| *Plethodon ventralis* | 1 | 1 | Highton 1997 |
| *Plethodon dorsalis* | 1 | 1 | Highton 1997 |
| *Plethodon yonahlossee* | 1 | 1 | Highton 1995; Highton and Peabody 2000; Wiens et al. 2006 |
| *Plethodon petraeus* | 1 | 1 | Highton 1995; Highton and Peabody 2000; Wiens et al. 2006 |
| *Plethodon kentucki* | 1 | 1 | Highton 1995; Highton and Peabody 2000; Wiens et al. 2006 |
| *Plethodon ouachitae* | 1 | 1 | Shepard et al. 2011 |
| *Plethodon fourchensis* | 1 | 1 | Shepard et al. 2011 |
| *Plethodon jordani* | 1 | 1 | Highton 1995; Highton and Peabody 2000; Wiens et al. 2006 |
| *Plethodon shermani* | 1 | 1 | Highton 1995; Highton and Peabody 2000; Wiens et al. 2006 |
| *Plethodon metcalfi* | 1 | 1 | Highton 1995; Highton and Peabody 2000; Wiens et al. 2006 |
| *Plethodon montanus* | 1 | 1 | Highton 1995; Highton and Peabody 2000; Wiens et al. 2006 |
| *Plethodon chlorobryonis* | 1 | 1 | Highton 1995; Highton and Peabody 2000; Wiens et al. 2006 |
| *Plethodon teyahalee* | 1 | 1 | Highton 1995; Highton and Peabody 2000; Wiens et al. 2006 |
| *Plethodon cylindraceus* | 1 | 1 | Highton 1995; Highton and Peabody 2000; Wiens et al. 2006 |
| *Plethodon chattahoochee* | 1 | 1 | Highton 1995; Highton and Peabody 2000; Wiens et al. 2006 |
| *Plethodon cheoah* | 1 | 1 | Highton 1995; Highton and Peabody 2000; Wiens et al. 2006 |
| *Plethodon glutinosus* | 1 | 1 | Highton 1995; Highton and Peabody 2000; Wiens et al. 2006 |
| *Plethodon aureolus* | 1 | 1 | Highton et al. 2012 |
| *Gyrinophilus porphyriticus* | 0 | 1 | Niemiller et al. 2008; Kuchta et al 2016 |
| *Gyrinophilus gulolineatus* | 0 | 1 | Niemiller et al. 2008; Kuchta et al 2016 |
| *Gyrinophilus palleucus* | 0 | 1 | Niemiller et al. 2008 |
| *Eurycea lucifuga* | 1 | 1 | Smith 1964 |
| *Eurycea longicauda* | 1 | 1 | Smith 1964 |
| *Eurycea wilderae* | 1 | 1 | Sweet 1984; Guttman and Karlin 1986; Kozak and Montanucci 2001 |
| *Eurycea cirrigera* | 1 | 1 | Sweet 1984; Guttman and Karlin 1986; Kozak and Montanucci 2001 |
| *Eurycea sosorum* | 1 | 1 | Bendik et al. 2013 |
| *Eurycea nana* | 1 | 1 | Bendik et al. 2013 |
| *Eurycea neotenes* | 1 | 1 | Sweet 1984; Guttman and Karlin 1986; Kozak and Montanucci 2001 |
| *Eurycea tridentifera* | 1 | 1 | Sweet 1984; Guttman and Karlin 1986; Kozak and Montanucci 2001 |
| *Batrachoseps simatus* | 1 | 1 | Jockusch et al. 2012 |
| *Batrachoseps gavilanensis* | 1 | 1 | Jockusch and Wake 2002 |
| *Batrachoseps luciae* | 1 | 1 | Jockusch et al. 2012; Jockusch et al., 2001 |
| *Batrachoseps minor* | 1 | 1 | Jockusch et al. 2001 |
| *Bolitoglossa lincolni* | 1 | 1 | Wake and Lynch 1982; Wake et al. 1980 |
| *Bolitoglossa franklini* | 1 | 1 | Wake and Lynch 1982; Wake et al. 1980 |

**Table S1: Species designation as hybridizable according to the narrow and broad datasets and corresponding literature.** Note this includes only those species included in our phylogeny

| Model | Trait-dependent Diversification | All Species | | | Sympatric Species | | | Plethodontids | | | Non-Plethodontids | | |
|---|---|---|---|---|---|---|---|---|---|---|---|---|---|
| | | *AICc* | *ΔAICc* | *Akaike Weights* | *AICc* | *ΔAICc* | *Akaike Weights* | *AICc* | *ΔAICc* | *Akaike Weights* | *AICc* | *ΔAICc* | *Akaike Weights* |
| BiSSE Null: One transition | No | 4173.37 | 182.74 | 0.000 | 3391.76 | 144.62 | 0.000 | 2635.67 | 83.12 | 0.000 | 1491.98 | 56.75 | 0.000 |
| BiSSE Null: Two transitions | No | 4090.55 | 99.92 | 0.000 | 3324.28 | 77.14 | 0.000 | 2577.00 | 24.46 | 0.000 | 1467.11 | 31.89 | 0.000 |
| CID-2: All transitions, all equal | No | 4117.81 | 127.17 | 0.000 | 3353.04 | 105.90 | 0.000 | 2630.00 | 77.45 | 0.000 | 1466.64 | 31.42 | 0.000 |
| CID-2: Three transitions, no double | No | 4021.40 | 30.77 | 0.000 | 3273.42 | 26.28 | 0.000 | 2562.99 | 10.44 | 0.004 | 1447.67 | 12.44 | 0.002 |
| CID-2: No double transitions, all equal | No | 4104.69 | 114.05 | 0.000 | 3340.20 | 93.06 | 0.000 | 2628.93 | 76.39 | 0.000 | 1462.82 | 27.59 | 0.000 |
| CID-4: Three transitions | No | 4011.10 | 20.47 | 0.000 | 3269.79 | 22.65 | 0.000 | 2560.74 | 8.19 | 0.013 | 1443.78 | 8.55 | 0.012 |
| CID-4: Equal transitions | No | 4087.30 | 96.67 | 0.000 | 3329.69 | 82.55 | 0.000 | 2618.14 | 65.59 | 0.000 | 1461.35 | 26.13 | 0.000 |
| BiSSE: All parameters free | Yes | 4061.35 | 70.72 | 0.000 | 3287.58 | 40.44 | 0.000 | 2558.34 | 5.80 | 0.044 | 1459.86 | 24.64 | 0.000 |
| HiSSE: No NH-B, all transitions | Yes | 4073.86 | 83.22 | 0.000 | 3300.24 | 53.10 | 0.000 | 2568.95 | 16.40 | 0.000 | 1493.21 | 57.99 | 0.000 |
| HiSSE: No NH-B, no double transitions | Yes | 4033.85 | 43.22 | 0.000 | 3277.22 | 30.08 | 0.000 | 2560.89 | 8.35 | 0.012 | 1463.94 | 28.71 | 0.000 |
| HiSSE: No H-B, all transitions | Yes | 3992.96 | 2.33 | 0.126 | 3292.19 | 45.05 | 0.000 | 2556.66 | 4.11 | 0.102 | 1454.11 | 18.88 | 0.000 |
| HiSSE: No H-B, no double transitions | Yes | **3990.63** | **0.00** | **0.402** | **3247.14** | **0.00** | **0.556** | **2552.55** | **0.00** | **0.793** | **1435.22** | **0.00** | **0.879** |
| HiSSE: All parameters free | Yes | 3994.05 | 3.42 | 0.073 | 3247.66 | 0.52 | 0.429 | 2563.94 | 11.40 | 0.003 | 1448.10 | 12.88 | 0.001 |
| HiSSE: No double transitions | Yes | 3990.65 | 0.01 | 0.399 | 3254.43 | 7.29 | 0.015 | 2559.15 | 6.60 | 0.029 | 1439.46 | 4.24 | 0.106 |

**Table S2: The 14 models fitted in our analysis using the narrow dataset assuming we have sampled all species that hybridize in nature**. Models varied in number of hidden states included as well as in the number of unique transitions among states permitted. HiSSE models contain hidden states, whereas BiSSE models do not. Double transitions are those in which two-state jumps occur (i.e. NH-**A** to H-B or vice versa). CID stands for character independent, thus CID-2 and CID-4 represent the two novel null models of trait-independent diversification that may be implemented in the HiSSE framework. Rows in bold indicate best-fit models.

| Model | Trait-dependent Diversification | All Species | | | Sympatric Species | | | Plethodontids | | | Non-Plethodontids | | |
|---|---|---|---|---|---|---|---|---|---|---|---|---|---|
| | | *AICc* | *ΔAICc* | *Akaike Weights* | *AICc* | *ΔAICc* | *Akaike Weights* | *AICc* | *ΔAICc* | *Akaike Weights* | *AICc* | *ΔAICc* | *Akaike Weights* |
| BiSSE Null: One transition | No | 4210.44 | 154.20 | 0.000 | 3443.09 | 122.04 | 0.000 | 2646.49 | 77.97 | 0.000 | 1515.40 | 35.86 | 0.000 |
| BiSSE Null: Two transitions | No | 4148.12 | 91.88 | 0.000 | 3393.87 | 72.82 | 0.000 | 2592.99 | 24.47 | 0.000 | 1502.54 | 23.00 | 0.000 |
| CID-2: All transitions, all equal | No | 4151.24 | 95.00 | 0.000 | 3401.87 | 80.82 | 0.000 | 2633.68 | 65.16 | 0.000 | 1489.38 | 9.84 | 0.005 |
| CID-2: Three transitions, no double | No | 4077.94 | 21.70 | 0.000 | 3342.23 | 21.18 | 0.000 | 2579.14 | 10.61 | 0.003 | 1483.82 | 4.27 | 0.081 |
| CID-2: No double transitions, all equal | No | 4139.34 | 83.10 | 0.000 | 3389.73 | 68.68 | 0.000 | 2639.53 | 71.01 | 0.000 | 1487.11 | 7.57 | 0.016 |
| CID-4: Three transitions | No | 4069.63 | 13.39 | 0.001 | 3341.48 | 20.43 | 0.000 | 2577.48 | 8.95 | 0.007 | **1479.54** | **0.00** | **0.683** |
| CID-4: Equal transitions | No | 4125.26 | 69.02 | 0.000 | 3381.94 | 60.89 | 0.000 | 2629.63 | 61.11 | 0.000 | 1489.02 | 9.47 | 0.006 |
| BiSSE: All parameters free | Yes | 4119.74 | 63.50 | 0.000 | 3357.57 | 36.52 | 0.000 | 2574.13 | 5.60 | 0.040 | 1510.44 | 30.90 | 0.000 |
| HiSSE: No NH-B, all transitions | Yes | 4132.25 | 76.01 | 0.000 | 3366.68 | 45.63 | 0.000 | 2580.13 | 11.60 | 0.002 | 1516.55 | 37.01 | 0.000 |
| HiSSE: No NH-B, no double transitions | Yes | 4089.76 | 33.52 | 0.000 | 3347.88 | 26.83 | 0.000 | 2576.99 | 8.47 | 0.010 | 1506.97 | 27.43 | 0.000 |
| HiSSE: No H-B, all transitions | Yes | 4058.84 | 2.60 | 0.144 | 3368.30 | 47.25 | 0.000 | **2568.52** | **0.00** | **0.658** | 1494.62 | 15.08 | 0.000 |
| HiSSE: No H-B, no double transitions | Yes | 4057.57 | 1.33 | 0.273 | 3325.01 | 3.96 | 0.087 | 2570.40 | 1.88 | 0.257 | 1482.56 | 3.02 | 0.151 |
| HiSSE: All parameters free | Yes | **4056.24** | **0.00** | **0.530** | **3321.05** | **0.00** | **0.630** | 2581.88 | 13.36 | 0.001 | 1489.90 | 10.36 | 0.004 |
| HiSSE: No double transitions | Yes | 4060.85 | 4.62 | 0.053 | 3322.65 | 1.60 | 0.283 | 2575.38 | 6.85 | 0.021 | 1484.58 | 5.03 | 0.055 |

**Table S3: The 14 models fitted in our analysis using the narrow dataset assuming we have sampled all species that hybridize in nature**. Models varied in number of hidden states included as well as in the number of unique transitions among states permitted. HiSSE models contain hidden states, whereas BiSSE models do not. Double transitions are those in which two-state jumps occur (i.e. NH-**A** to H-B or vice versa). CID stands for character independent, thus CID-2 and CID-4 represent the two novel null models of trait-independent diversification that may be implemented in the HiSSE framework. Rows in bold indicate best-fit models.

| Model | Trait-dependent Diversification | All Species | | | Sympatric Species | | | Plethodontids | | | Non-Plethodontids | | |
|---|---|---|---|---|---|---|---|---|---|---|---|---|---|
| | | *AICc* | *ΔAICc* | *Akaike Weights* | *AICc* | *ΔAICc* | *Akaike Weights* | *AICc* | *ΔAICc* | *Akaike Weights* | *AICc* | *ΔAICc* | *Akaike Weights* |
| BiSSE Null: One transition | No | 4169.69 | 181.78 | 0.000 | 3390.54 | 153.16 | 0.000 | 2637.46 | 94.86 | 0.000 | 1486.77 | 49.77 | 0.000 |
| BiSSE Null: Two transitions | No | 4093.52 | 105.61 | 0.000 | 3329.65 | 92.27 | 0.000 | 2582.39 | 39.79 | 0.000 | 1464.96 | 27.96 | 0.000 |
| CID-2: All transitions, all equal | No | 4112.63 | 124.72 | 0.000 | 3350.40 | 113.02 | 0.000 | 2623.60 | 81.00 | 0.000 | 1461.89 | 24.89 | 0.000 |
| CID-2: Three transitions, no double | No | 4023.51 | 35.60 | 0.000 | 3278.34 | 40.97 | 0.000 | 2565.84 | 23.24 | 0.000 | 1437.56 | 0.56 | 0.383 |
| CID-2: No double transitions, all equal | No | 4098.26 | 110.35 | 0.000 | 3336.68 | 99.30 | 0.000 | 2627.33 | 84.73 | 0.000 | 1458.20 | 21.20 | 0.000 |
| CID-4: Three transitions | No | 4018.18 | 30.27 | 0.000 | 3275.63 | 38.25 | 0.000 | 2567.99 | 25.39 | 0.000 | 1444.34 | 7.34 | 0.013 |
| CID-4: Equal transitions | No | 4093.95 | 106.04 | 0.000 | 3334.87 | 97.49 | 0.000 | 2624.46 | 81.86 | 0.000 | 1464.29 | 27.29 | 0.000 |
| BiSSE: All parameters free | Yes | 4047.43 | 59.52 | 0.000 | 3272.86 | 35.49 | 0.000 | 2552.10 | 9.50 | 0.008 | 1457.53 | 20.53 | 0.000 |
| HiSSE: No NH-B, all transitions | Yes | 4040.95 | 53.04 | 0.000 | 3284.37 | 47.00 | 0.000 | 2564.89 | 22.29 | 0.000 | 1471.08 | 34.07 | 0.000 |
| HiSSE: No NH-B, no double transitions | Yes | 4023.47 | 35.55 | 0.000 | 3268.31 | 30.94 | 0.000 | 2559.83 | 17.23 | 0.000 | 1446.69 | 9.69 | 0.004 |
| HiSSE: No H-B, all transitions | Yes | 4059.91 | 71.99 | 0.000 | 3246.77 | 9.39 | 0.008 | 2546.72 | 4.12 | 0.112 | 1440.89 | 3.88 | 0.073 |
| HiSSE: No H-B, no double transitions | Yes | 3988.53 | 0.62 | 0.380 | 3251.58 | 14.21 | 0.001 | **2542.60** | **0.00** | **0.875** | 1443.75 | 6.75 | 0.017 |
| HiSSE: All parameters free | Yes | **3987.91** | **0.00** | **0.518** | 3241.24 | 3.86 | 0.125 | 2556.65 | 14.05 | 0.001 | 1446.79 | 9.78 | 0.004 |
| HiSSE: No double transitions | Yes | 3991.16 | 3.24 | 0.102 | **3237.38** | **0.00** | **0.866** | 2552.92 | 10.32 | 0.005 | **1437.00** | **0.00** | **0.506** |

**Table S4: The 14 models fitted in our analysis using the narrow dataset assuming the prevalence of hybridization in nature is equal to that which we observe in our data**. Models varied in number of hidden states included as well as in the number of unique transitions among states permitted. HiSSE models contain hidden states, whereas BiSSE models do not. Double transitions are those in which two-state jumps occur (i.e. NH-**A** to H-B or vice versa). CID stands for character independent, thus CID-2 and CID-4 represent the two novel null models of trait-independent diversification that may be implemented in the HiSSE framework. Rows in bold indicate best-fit models.

| Model | Trait-dependent Diversification | All Species | | | Sympatric Species | | | Plethodontids | | | Non-Plethodontids | | |
|---|---|---|---|---|---|---|---|---|---|---|---|---|---|
| | | *AICc* | *ΔAICc* | *Akaike Weights* | *AICc* | *ΔAICc* | *Akaike Weights* | *AICc* | *ΔAICc* | *Akaike Weights* | *AICc* | *ΔAICc* | *Akaike Weights* |
| BiSSE Null: One transition | No | 4209.37 | 162.16 | 0.000 | 3443.21 | 132.92 | 0.000 | 2649.83 | 99.41 | 0.000 | 1510.60 | 36.34 | 0.000 |
| BiSSE Null: Two transitions | No | 4152.32 | 105.11 | 0.000 | 3400.60 | 90.31 | 0.000 | 2599.40 | 48.98 | 0.000 | 1500.97 | 26.72 | 0.000 |
| CID-2: All transitions, all equal | No | 4149.73 | 102.52 | 0.000 | 3401.32 | 91.03 | 0.000 | 2641.06 | 90.65 | 0.000 | 1488.27 | 14.02 | 0.001 |
| CID-2: Three transitions, no double | No | 4081.99 | 34.78 | 0.000 | 3348.52 | 38.22 | 0.000 | 2582.60 | 32.19 | 0.000 | **1474.25** | **0.00** | **0.932** |
| CID-2: No double transitions, all equal | No | 4137.14 | 89.93 | 0.000 | 3389.12 | 78.82 | 0.000 | 2639.06 | 88.65 | 0.000 | 1485.78 | 11.53 | 0.003 |
| CID-4: Three transitions | No | 4076.88 | 29.67 | 0.000 | 3347.10 | 36.81 | 0.000 | 2585.01 | 34.59 | 0.000 | 1480.35 | 6.10 | 0.044 |
| CID-4: Equal transitions | No | 4132.20 | 84.99 | 0.000 | 3387.42 | 77.13 | 0.000 | 2636.50 | 86.08 | 0.000 | 1490.92 | 16.66 | 0.000 |
| BiSSE: All parameters free | Yes | 4098.88 | 51.67 | 0.000 | 3340.46 | 30.16 | 0.000 | 2567.40 | 16.98 | 0.000 | 1493.79 | 19.54 | 0.000 |
| HiSSE: No NH-B, all transitions | Yes | 4110.92 | 63.71 | 0.000 | 3353.08 | 42.79 | 0.000 | 2580.19 | 29.77 | 0.000 | 1512.23 | 37.98 | 0.000 |
| HiSSE: No NH-B, no double transitions | Yes | 4077.76 | 30.55 | 0.000 | 3336.59 | 26.29 | 0.000 | 2575.86 | 25.44 | 0.000 | 1485.53 | 11.28 | 0.003 |
| HiSSE: No H-B, all transitions | Yes | 4057.10 | 9.89 | 0.006 | 3333.84 | 23.54 | 0.000 | 2562.23 | 11.81 | 0.003 | 1493.27 | 19.02 | 0.000 |
| HiSSE: No H-B, no double transitions | Yes | 4054.97 | 7.76 | 0.017 | 3325.33 | 15.03 | 0.001 | **2557.05** | **6.63** | **0.035** | 1482.48 | 8.23 | 0.015 |
| HiSSE: All parameters free | Yes | **4047.21** | **0.00** | **0.847** | 3315.56 | 5.26 | 0.067 | 2569.97 | 19.55 | 0.000 | 1487.54 | 13.28 | 0.001 |
| HiSSE: No double transitions | Yes | 4050.96 | 3.75 | 0.130 | **3310.29** | **0.00** | **0.932** | 2550.42 | 0.00 | 0.962 | 1504.32 | 30.06 | 0.000 |

**Table S5: The 14 models fitted in our analysis using the broad dataset assuming the prevalence of hybridization in nature is equal to that which we observe in our data**. Models varied in number of hidden states included as well as in the number of unique transitions among states permitted. HiSSE models contain hidden states, whereas BiSSE models do not. Double transitions are those in which two-state jumps occur (i.e. NH-A to H-B or vice versa). CID stands for character independent, thus CID-2 and CID-4 represent the two novel null models of trait-independent diversification that may be implemented in the HiSSE framework. Rows in bold indicate best-fit model.

| Model | Trait-dependent Diversification | All Species | | | Sympatric Species | | | Plethodontids | | | Non-Plethodontids | | |
|---|---|---|---|---|---|---|---|---|---|---|---|---|---|
| | | *AICc* | *ΔAICc* | *Akaike Weights* | *AICc* | *ΔAICc* | *Akaike Weights* | *AICc* | *ΔAICc* | *Akaike Weights* | *AICc* | *ΔAICc* | *Akaike Weights* |
| BiSSE Null: One transition | No | 4170.31 | 190.81 | 0.000 | 3391.55 | 150.94 | 0.000 | 2644.86 | 93.89 | 0.000 | 1486.77 | 42.49 | 0.000 |
| BiSSE Null: Two transitions | No | 4098.94 | 119.44 | 0.000 | 3334.63 | 94.01 | 0.000 | 2589.79 | 38.82 | 0.000 | 1464.96 | 20.68 | 0.000 |
| CID-2: All transitions, all equal | No | 4110.14 | 130.64 | 0.000 | 3349.15 | 108.54 | 0.000 | 2632.11 | 81.14 | 0.000 | 1466.48 | 22.20 | 0.000 |
| CID-2: Three transitions, no double | No | 4027.61 | 48.11 | 0.000 | 3281.92 | 41.31 | 0.000 | 2570.98 | 20.01 | 0.000 | 1454.05 | 9.77 | 0.005 |
| CID-2: No double transitions, all equal | No | 4096.81 | 117.31 | 0.000 | 3336.35 | 95.74 | 0.000 | 2633.89 | 82.91 | 0.000 | 1464.02 | 19.74 | 0.000 |
| CID-4: Three transitions | No | 4022.21 | 42.71 | 0.000 | 3278.06 | 37.45 | 0.000 | 2574.33 | 23.36 | 0.000 | 1449.92 | 5.64 | 0.039 |
| CID-4: Equal transitions | No | 4097.25 | 117.74 | 0.000 | 3338.06 | 97.45 | 0.000 | 2633.18 | 82.20 | 0.000 | 1469.52 | 25.24 | 0.000 |
| BiSSE: All parameters free | Yes | 4039.81 | 60.31 | 0.000 | 3268.29 | 27.68 | 0.000 | 2563.16 | 12.19 | 0.001 | 1458.20 | 13.92 | 0.001 |
| HiSSE: No NH-B, no double transitions | Yes | 4020.58 | 41.08 | 0.000 | 3266.29 | 25.68 | 0.000 | 2568.96 | 17.99 | 0.000 | 1459.34 | 15.06 | 0.000 |
| HiSSE: No NH-B, all transitions | Yes | 4038.08 | 58.58 | 0.000 | 3279.81 | 39.19 | 0.000 | 2575.96 | 24.98 | 0.000 | 1471.11 | 26.83 | 0.000 |
| HiSSE: No H-B , no double transitions | Yes | **3979.50** | **0.00** | **0.715** | 3274.05 | 33.44 | 0.000 | **2550.97** | **0.00** | **0.523** | 1448.35 | 4.07 | 0.085 |
| HiSSE: No H-B, all transitions | Yes | 3990.73 | 11.22 | 0.003 | 3249.90 | 9.29 | 0.009 | 2551.17 | 0.19 | 0.475 | 1446.57 | 2.29 | 0.206 |
| HiSSE: All parameters free | Yes | 3986.74 | 7.24 | 0.019 | **3240.61** | **0.00** | **0.980** | 2565.34 | 14.36 | 0.000 | 1451.73 | 7.45 | 0.016 |
| HiSSE: No double transitions | Yes | 3981.50 | 2.00 | 0.263 | 3249.66 | 9.05 | 0.011 | 2565.26 | 14.29 | 0.000 | **1444.28** | **0.00** | **0.648** |

**Table S6: The 14 models fitted in our analysis using the narrow dataset assuming 20% of species hybridize.** Models varied in number of hidden states included as well as in the number of unique transitions among states permitted. HiSSE models contain hidden states, whereas BiSSE models do not. Double transitions are those in which two-state jumps occur (i.e. NH-**A** to H-B or vice versa). CID stands for character independent, thus CID-2 and CID-4 represent the two novel null models of trait-independent diversification that may be implemented in the HiSSE framework. Rows in bold indicate best-fit models.

| Model | Trait-dependent Diversification | All Species | | | Sympatric Species | | | Plethodontids | | | Non-plethodontids | | |
|---|---|---|---|---|---|---|---|---|---|---|---|---|---|
| | | *AICc* | *ΔAICc* | *Akaike Weights* | *AICc* | *ΔAICc* | *Akaike Weights* | *AICc* | *ΔAICc* | *Akaike Weights* | *AICc* | *ΔAICc* | *Akaike Weights* |
| BiSSE Null: One transition | No | 4209.55 | 162.81 | 0.000 | 3442.40 | 123.97 | 0.000 | 2652.92 | 98.52 | 0.000 | 1513.34 | 39.60 | 0.000 |
| BiSSE Null: Two transitions | No | 4152.91 | 106.17 | 0.000 | 3395.12 | 76.69 | 0.000 | 2604.52 | 50.12 | 0.000 | 1501.47 | 27.74 | 0.000 |
| CID-2: All transitions, all equal | No | 4149.59 | 102.86 | 0.000 | 3401.56 | 83.14 | 0.000 | 2636.08 | 81.68 | 0.000 | 1488.83 | 15.10 | 0.000 |
| CID-2: Three transitions, no double | No | 4082.41 | 35.67 | 0.000 | 3343.39 | 24.96 | 0.000 | 2588.50 | 34.10 | 0.000 | **1473.74** | **0.00** | **0.949** |
| CID-2: No double transitions, all equal | No | 4137.09 | 90.36 | 0.000 | 3388.86 | 70.43 | 0.000 | 2638.55 | 84.14 | 0.000 | 1486.23 | 12.49 | 0.002 |
| CID-4: Three transitions | No | 4075.91 | 29.18 | 0.000 | 3355.39 | 36.96 | 0.000 | 2584.89 | 30.48 | 0.000 | 1480.55 | 6.82 | 0.031 |
| CID-4: Equal transitions | No | 4144.60 | 97.86 | 0.000 | 3395.84 | 77.41 | 0.000 | 2640.87 | 86.46 | 0.000 | 1490.21 | 16.48 | 0.000 |
| BiSSE: All parameters free | Yes | 4097.73 | 50.99 | 0.000 | 3350.59 | 32.16 | 0.000 | 2565.14 | 10.74 | 0.005 | 1510.75 | 37.01 | 0.000 |
| HiSSE: No NH-B, all transitions | Yes | 4077.04 | 30.31 | 0.000 | 3343.36 | 24.94 | 0.000 | 2573.61 | 19.21 | 0.000 | 1489.37 | 15.63 | 0.000 |
| HiSSE: No NH-B, no double transitions | Yes | 4076.68 | 29.94 | 0.000 | 3360.11 | 41.69 | 0.000 | 2577.93 | 23.53 | 0.000 | 1516.29 | 42.56 | 0.000 |
| HiSSE: No H-B, all transitions | Yes | 4049.90 | 3.16 | 0.130 | 3355.30 | 36.87 | 0.000 | 2564.85 | 10.45 | 0.005 | 1492.63 | 18.89 | 0.000 |
| HiSSE: No H-B, no double transitions | Yes | 4054.96 | 8.22 | 0.010 | 3329.95 | 11.52 | 0.002 | **2554.40** | **0.00** | **0.970** | 1482.37 | 8.64 | 0.013 |
| HiSSE: All parameters free | Yes | **4046.74** | **0.00** | **0.631** | 3319.57 | 1.14 | 0.361 | 2566.55 | 12.15 | 0.002 | 1488.73 | 15.00 | 0.001 |
| HiSSE: No double transitions | Yes | 4048.76 | 2.02 | 0.229 | **3318.43** | **0.00** | **0.637** | 2562.33 | 7.93 | 0.018 | 1484.90 | 11.17 | 0.004 |

**Table S7: The 14 models fitted in our analysis using the broad dataset assuming 20% of species hybridize.** Models varied in number of hidden states included as well as in the number of unique transitions among states permitted. HiSSE models contain hidden states, whereas BiSSE models do not. Double transitions are those in which two-state jumps occur (i.e. NH-A to H-B or vice versa). CID stands for character independent, thus CID-2 and CID-4 represent the two novel null models of trait-independent diversification that may be implemented in the HiSSE framework. Rows in bold indicate best-fit model.

| Model | Trait-dependent Diversification | All Species | | | Sympatric Species | | | Plethodontids | | | Non-Plethodontids | | |
|---|---|---|---|---|---|---|---|---|---|---|---|---|---|
| | | *AICc* | *ΔAICc* | *Akaike Weights* | *AICc* | *ΔAICc* | *Akaike Weights* | *AICc* | *ΔAICc* | *Akaike Weights* | *AICc* | *ΔAICc* | *Akaike Weights* |
| BiSSE Null: One transition | No | 4175.19 | 196.80 | 0.000 | 3393.81 | 162.88 | 0.000 | 2648.43 | 108.72 | 0.000 | 1482.50 | 40.69 | 0.000 |
| BiSSE Null: Two transitions | No | 4117.89 | 139.50 | 0.000 | 3343.74 | 112.81 | 0.000 | 2604.82 | 65.11 | 0.000 | 1468.44 | 26.63 | 0.000 |
| CID-2: All transitions, all equal | No | 4106.77 | 128.38 | 0.000 | 3347.79 | 116.87 | 0.000 | 2634.37 | 94.66 | 0.000 | 1457.13 | 15.32 | 0.000 |
| CID-2: Three transitions, no double | No | 4047.25 | 68.86 | 0.000 | 3292.72 | 61.79 | 0.000 | 2580.60 | 40.89 | 0.000 | 1448.30 | 6.49 | 0.024 |
| CID-2: No double transitions, all equal | No | 4097.28 | 118.89 | 0.000 | 3337.17 | 106.24 | 0.000 | 2628.82 | 89.11 | 0.000 | 1455.57 | 13.76 | 0.001 |
| CID-4: Three transitions | No | 4006.14 | 27.75 | 0.000 | 3282.41 | 51.48 | 0.000 | 2546.08 | 6.37 | 0.038 | 1448.71 | 6.90 | 0.019 |
| CID-4: Equal transitions | No | 4083.10 | 104.71 | 0.000 | 3328.09 | 97.16 | 0.000 | 2606.16 | 66.44 | 0.000 | 1472.13 | 30.32 | 0.000 |
| BiSSE: All parameters free | Yes | 4046.68 | 68.29 | 0.000 | 3264.66 | 33.73 | 0.000 | 2549.85 | 10.14 | 0.006 | 1457.41 | 15.60 | 0.000 |
| HiSSE: No NH-B, all transitions | Yes | 4059.20 | 80.81 | 0.000 | 3276.06 | 45.13 | 0.000 | 2658.06 | 118.35 | 0.000 | 1469.41 | 27.60 | 0.000 |
| HiSSE: No NH-B, no double transitions | Yes | 4020.54 | 42.16 | 0.000 | 3265.99 | 35.06 | 0.000 | 2557.47 | 17.76 | 0.000 | 1459.32 | 17.51 | 0.000 |
| HiSSE: No H-B, all transitions | Yes | **3978.39** | **0.00** | **0.862** | 3242.87 | 11.95 | 0.003 | 2547.05 | 7.34 | 0.023 | 1459.10 | 17.29 | 0.000 |
| HiSSE: No H-B, no double transitions | Yes | 3982.24 | 3.86 | 0.125 | **3230.93** | **0.00** | **0.991** | **2539.71** | **0.00** | **0.915** | 1443.25 | 1.44 | 0.298 |
| HiSSE: All parameters free | Yes | 3986.85 | 8.46 | 0.013 | 3240.96 | 10.03 | 0.007 | 2547.83 | 8.12 | 0.016 | 1446.99 | 5.18 | 0.046 |
| HiSSE: No double transitions | Yes | 4000.55 | 22.17 | 0.000 | 3250.83 | 19.90 | 0.000 | 2552.34 | 12.63 | 0.002 | **1441.81** | **0.00** | **0.612** |

**Table S8: The 14 models fitted in our analysis using the narrow dataset assuming 30% of species hybridize**. Models varied in number of hidden states included as well as in the number of unique transitions among states permitted. HiSSE models contain hidden states, whereas BiSSE models do not. Double transitions are those in which two-state jumps occur (i.e. NH-**A** to H-B or vice versa). CID stands for character independent, thus CID-2 and CID-4 represent the two novel null models of trait-independent diversification that may be implemented in the HiSSE framework. Rows in bold indicate best-fit models.

| Model | Trait-dependent Diversification | All Species | | | Sympatric Species | | | Plethodontids | | | Non-Plethodontids | | |
|---|---|---|---|---|---|---|---|---|---|---|---|---|---|
| | | *AICc* | *ΔAICc* | *Akaike Weights* | *AICc* | *ΔAICc* | *Akaike Weights* | *AICc* | *ΔAICc* | *Akaike Weights* | *AICc* | *ΔAICc* | *Akaike Weights* |
| BiSSE Null: One transition | No | 4209.37 | 162.16 | 0.000 | 3443.21 | 132.92 | 0.000 | 2649.83 | 99.41 | 0.000 | 1510.60 | 36.34 | 0.000 |
| BiSSE Null: Two transitions | No | 4152.32 | 105.11 | 0.000 | 3400.60 | 90.31 | 0.000 | 2599.40 | 48.98 | 0.000 | 1500.97 | 26.72 | 0.000 |
| CID-2: All transitions, all equal | No | 4149.73 | 102.52 | 0.000 | 3401.32 | 91.03 | 0.000 | 2641.06 | 90.65 | 0.000 | 1488.27 | 14.02 | 0.001 |
| CID-2: Three transitions, no double | No | 4081.99 | 34.78 | 0.000 | 3348.52 | 38.22 | 0.000 | 2582.60 | 32.19 | 0.000 | **1474.25** | **0.00** | **0.932** |
| CID-2: No double transitions, all equal | No | 4137.14 | 89.93 | 0.000 | 3389.12 | 78.82 | 0.000 | 2639.06 | 88.65 | 0.000 | 1485.78 | 11.53 | 0.003 |
| CID-4: Three transitions | No | 4076.88 | 29.67 | 0.000 | 3347.10 | 36.81 | 0.000 | 2585.01 | 34.59 | 0.000 | 1480.35 | 6.10 | 0.044 |
| CID-4: Equal transitions | No | 4132.20 | 84.99 | 0.000 | 3387.42 | 77.13 | 0.000 | 2636.50 | 86.08 | 0.000 | 1490.92 | 16.66 | 0.000 |
| BiSSE: All parameters free | Yes | 4098.88 | 51.67 | 0.000 | 3340.46 | 30.16 | 0.000 | 2567.40 | 16.98 | 0.000 | 1493.79 | 19.54 | 0.000 |
| HiSSE: No NH-B, all transitions | Yes | 4110.92 | 63.71 | 0.000 | 3353.08 | 42.79 | 0.000 | 2580.19 | 29.77 | 0.000 | 1512.23 | 37.98 | 0.000 |
| HiSSE: No NH-B, no double transitions | Yes | 4077.76 | 30.55 | 0.000 | 3336.59 | 26.29 | 0.000 | 2575.86 | 25.44 | 0.000 | 1485.53 | 11.28 | 0.003 |
| HiSSE: No H-B, all transitions | Yes | 4057.10 | 9.89 | 0.006 | 3333.84 | 23.54 | 0.000 | 2562.23 | 11.81 | 0.003 | 1493.27 | 19.02 | 0.000 |
| HiSSE: No H-B, no double transitions | Yes | 4054.97 | 7.76 | 0.017 | 3325.33 | 15.03 | 0.001 | 2557.05 | 6.63 | 0.035 | 1482.48 | 8.23 | 0.015 |
| HiSSE: All parameters free | Yes | **4047.21** | **0.00** | **0.847** | 3315.56 | 5.26 | 0.067 | 2569.97 | 19.55 | 0.000 | 1487.54 | 13.28 | 0.001 |
| HiSSE: No double transitions | Yes | 4050.96 | 3.75 | 0.130 | **3310.29** | **0.00** | **0.932** | **2550.42** | **0.00** | **0.962** | 1504.32 | 30.06 | 0.000 |

**Table S9: The 14 models fitted in our analysis using the broad dataset assuming 30% of species hybridize**. Models varied in number of hidden states included as well as in the number of unique transitions among states permitted. HiSSE models contain hidden states, whereas BiSSE models do not. Double transitions are those in which two-state jumps occur (i.e. NH-A to H-B or vice versa). CID stands for character independent, thus CID-2 and CID-4 represent the two novel null models of trait-independent diversification that may be implemented in the HiSSE framework. Rows in bold indicate best-fit model.

| Definition of Hybridization | Dataset | Model | Akaike Weight | Speciation | | | | Extinction | | | | Net Diversification | | | |
|---|---|---|---|---|---|---|---|---|---|---|---|---|---|---|---|
| | | | | *NH-A* | *NH-B* | *H-A* | *H-B* | *NH-A* | *NH-B* | *H-A* | *H-B* | *NH-A* | *NH-B* | *H-A* | *H-B* |
| **Narrow** | **All Species** | **HiSSE: No H-B, no double transitions** | **0.402** | **0.150** | **0.026** | **0.098** | **NA** | **3.09E-10** | **2.02E-10** | **0.078** | **NA** | **0.150** | **2.58E-2** | **0.021** | **NA** |
| | | HiSSE: No double transitions | 0.399 | 0.112 | 0.029 | 0.093 | 0.112 | 2.49E-9 | 0.028 | 1.91E-10 | 2.32E-10 | 0.112 | 1.44E-3 | 0.093 | 0.112 |
| | | HiSSE: No H-B, all transitions | 0.126 | 0.023 | 0.126 | 0.104 | NA | 0.015 | 0.037 | 2.15E-10 | NA | 0.008 | 8.91E-2 | 0.104 | NA |
| | | HiSSE: All parameters free | 0.073 | 0.118 | 0.032 | 0.132 | 0.051 | 2.44E-10 | 0.034 | 2.72E-10 | 1.05E-10 | 0.118 | -1.80E-3 | 0.132 | 0.051 |
| | **Sympatric Species** | **HiSSE: No H-B, no double transitions** | **0.556** | **0.124** | **0.021** | **0.093** | **NA** | **2.56E-10** | **0.064** | **1.91E-10** | **NA** | **0.124** | **-4.23E-2** | **0.093** | **NA** |
| | | HiSSE: All parameters free | 0.429 | 0.099 | 0.029 | 0.064 | 0.210 | 2.04E-10 | 0.033 | 1.32E-10 | 4.42E-10 | 0.099 | -3.50E-3 | 0.064 | 0.210 |
| | **Plethodontids** | **HiSSE: No H-B, no double transitions** | **0.793** | **0.222** | **0.059** | **0.058** | **NA** | **4.58E-10** | **1.21E-10** | **1.19E-10** | **NA** | **0.222** | **5.88E-2** | **0.058** | **NA** |
| | | HiSSE: No H-B, all transitions | 0.102 | 0.001 | 0.067 | 0.112 | NA | 0.003 | 1.37E-10 | 2.30E-10 | NA | -0.002 | 6.66E-2 | 0.112 | NA |
| | **Non-plethodontids** | **HiSSE: No H-B, no double transitions** | **0.879** | **0.157** | **0.032** | **0.069** | **NA** | **7.43E-04** | **0.096** | **0.001** | **NA** | **0.156** | **-6.37E-2** | **0.068** | **NA** |
| | | HiSSE: All parameters free | 0.106 | 0.128 | 0.026 | 0.066 | 0.101 | 2.64E-10 | 0.024 | 1.37E-10 | 0.001 | 0.128 | 2.80E-3 | 0.066 | 0.100 |
| **Broad** | **All Species** | **HiSSE: All parameters free** | **0.530** | **0.030** | **0.117** | **0.128** | **0.058** | **0.035** | **2.65E-10** | **2.40E-10** | **1.29E-10** | **-0.005** | **0.117** | **0.128** | **0.058** |
| | | HiSSE: No H-B, no double transitions | 0.273 | 0.144 | 0.023 | 0.093 | NA | 2.98E-10 | 0.068 | 1.92E-10 | NA | 0.144 | -0.045 | 0.093 | NA |
| | | HiSSE: No H-B, all transitions | 0.144 | 0.125 | 0.022 | 0.110 | NA | 0.048 | 0.023 | 0.001 | NA | 0.077 | 0.000 | 0.109 | NA |
| | | HiSSE: No double transitions | 0.053 | 0.026 | 0.137 | 0.087 | 0.248 | 0.017 | 0.059 | 1.79E-10 | 4.07E-3 | 0.009 | 0.078 | 0.087 | 0.244 |
| | **Sympatric Species** | **HiSSE: All parameters free** | **0.630** | **0.116** | **0.020** | **0.171** | **0.052** | **2.38E-10** | **0.060** | **3.52E-8** | **0.003** | **0.116** | **-0.040** | **0.171** | **0.049** |
| | | HiSSE: No double transitions | 0.283 | 0.027 | 0.091 | 0.119 | 0.063 | 0.044 | 0.006 | 2.46E-10 | 0.027 | -0.016 | 0.085 | 0.119 | 0.036 |
| | | HiSSE: No H-B, no double transitions | 0.087 | 0.121 | 0.019 | 0.087 | NA | 2.49E-10 | 5.66E-2 | 1.80E-10 | NA | 0.121 | -0.038 | 0.087 | NA |
| | **Plethodontids** | **HiSSE: No H-B, all transitions** | **0.658** | **0.005** | **0.074** | **0.120** | **NA** | **0.016** | **1.52E-10** | **2.65E-10** | **NA** | -0.011 | 0.073 | 0.120 | NA |
| | | HiSSE: No H-B, no double transitions | 0.257 | 0.190 | 0.057 | 0.078 | NA | 3.91E-10 | 1.17E-10 | 1.61E-10 | NA | 0.190 | 0.057 | 0.078 | NA |
| | **Non-plethodontids** | **CID-4: Three transitions** | **0.683** | **Below** | **Below** | **Below** | **Below** | **Below** | **Below** | **Below** | **Below** | **Below** | **Below** | **Below** | **Below** |
| | | HiSSE: No H-B, no double transitions | 0.151 | 0.112 | 0.012 | 0.054 | NA | 2.31E-10 | 1.30E-5 | 1.12E-10 | NA | 0.112 | 0.011 | 0.054 | NA |
| | | CID-2: Three transitions, no double | 0.081 | 0.057 | 0.104 | 0.057 | 0.104 | 0.053 | 0.036 | 0.053 | 0.036 | 0.004 | 0.068 | 0.004 | 0.068 |
| | | HiSSE: No double transitions | 0.055 | 0.021 | 0.095 | 0.111 | 0.051 | 0.017 | 1.95E-10 | 0.083 | 0.033 | 0.005 | 0.094 | 0.028 | 0.017 |
| | | **CID-4: Three transitions** | **0.683** | *A* | *B* | *C* | *D* | *A* | *B* | *C* | *D* | *A* | *B* | *C* | *D* |
| | | | | **0.097** | **0.013** | **0.079** | **0.013** | **1.99E-10** | **2.66E-11** | **1.63E-10** | **2.66E-11** | **9.67E-02** | **1.29E-02** | **7.90E-02** | **1.29E-02** |

**Table S10: Best fit models assuming we have sampled all species that hybridize in nature**. Included are models that received >5% Akaike weights for their respective analyses. For each dataset, the best fit model is bold. Maximum-likelihood parameter estimates for speciation, extinction, and net diversification are reported. Non-hybridizing is abbreviated as NH, Hybridizing as H; A and B indicate the two hidden states. CID-4 is shown separately, as there are four hidden states included in the model, each distributed without respect to the sampled trait.

| Definition of Hybridization | Dataset | Model | Akaike Weight | Speciation | | | | Extinction | | | | Net Diversification | | | |
|---|---|---|---|---|---|---|---|---|---|---|---|---|---|---|---|
| | | | | *NH-A* | *NH-B* | *H-A* | *H-B* | *NH-A* | *NH-B* | *H-A* | *H-B* | *NH-A* | *NH-B* | *H-A* | *H-B* |
| Narrow | **All Species** | **HiSSE: All parameters free** | **0.518** | **0.111** | **0.030** | **0.135** | **0.064** | **2.29E-10** | **0.030** | **0.001** | **0.001** | **0.111** | **-4.00E-4** | **0.134** | **0.063** |
| | | HiSSE: No H-B, no double transitions | 0.380 | 0.140 | 0.023 | 0.102 | NA | 2.89E-10 | 0.068 | 1.51E-4 | NA | 0.140 | -0.045 | 0.102 | NA |
| | | HiSSE: No double transitions | 0.102 | 0.027 | 0.108 | 0.133 | 0.135 | 0.057 | 0.014 | 0.004 | 0.048 | -0.030 | 0.094 | 0.129 | 0.087 |
| | **Sympatric Species** | **HiSSE: No double transitions** | **0.866** | **0.086** | **0.020** | **0.312** | **0.102** | **0.004** | **0.013** | **6.43E-10** | **2.10E-10** | **0.082** | **0.007** | **0.312** | **0.102** |
| | | HiSSE: All parameters free | 0.125 | 0.024 | 0.090 | 0.250 | 0.074 | 0.022 | 1.86E-10 | 5.97E-10 | 1.53E-10 | 0.002 | 0.090 | 0.250 | 0.074 |
| | **Plethodontids** | **HiSSE: No H-B, no double transitions** | **0.875** | **0.040** | **0.086** | **0.139** | **NA** | **0.002** | **1.77E-10** | **2.86E-10** | **NA** | **0.038** | **0.086** | **0.139** | **NA** |
| | | HiSSE: No H-B, all transitions | 0.112 | 0.006 | 0.072 | 0.128 | NA | 0.017 | 1.48E-10 | 2.65E-10 | NA | -0.011 | 0.072 | 0.128 | NA |
| | **Non-plethodontids** | **HiSSE: No double transitions** | **0.506** | **0.124** | **0.023** | **0.072** | **0.110** | **2.56E-10** | **0.018** | **0.001** | **0.004** | **0.124** | **0.005** | **0.070** | **0.106** |
| | | CID-2: Three transitions, no double | 0.383 | 0.083 | 0.014 | 0.083 | 0.014 | 1.71E-10 | 2.87E-11 | 1.71E-10 | 2.87E-11 | 0.083 | 0.014 | 0.083 | 0.014 |
| | | HiSSE: No H-B, all transitions | 0.073 | 0.124 | 0.023 | 0.072 | 0.110 | 2.56E-10 | 0.018 | 0.001 | 0.004 | 0.124 | 0.005 | 0.070 | 0.106 |
| Broad | **All Species** | **HiSSE: All parameters free** | **0.847** | **0.026** | **0.107** | **0.065** | **0.125** | **0.025** | **2.21E-10** | **1.33E-10** | **1.66E-4** | **0.001** | **0.107** | **0.064** | **0.125** |
| | | HiSSE: No double transitions | 0.130 | 0.025 | 0.090 | 0.124 | 0.241 | 0.075 | 0.020 | 0.004 | 4.96E-10 | -0.050 | 0.070 | 0.120 | 0.241 |
| | **Sympatric Species** | **HiSSE: No double transitions** | **0.932** | **0.022** | **0.067** | **0.094** | **0.193** | **0.023** | **0.007** | **0.006** | **3.98E-10** | **-0.001** | **0.060** | **0.088** | **0.193** |
| | | HiSSE: All parameters free | 0.067 | 0.021 | 0.085 | 0.146 | 0.064 | 0.025 | 1.76E-10 | 3.00E-10 | 0.009 | -0.004 | 0.085 | 0.146 | 0.055 |
| | **Plethodontids** | **HiSSE: No double transitions** | **0.962** | **0.016** | **0.085** | **0.117** | **0.141** | **0.048** | **1.76E-10** | **2.41E-10** | **0.006** | **-0.032** | **0.085** | **0.117** | **0.135** |
| | **Non-plethodontids** | **CID-2: Three transitions, no double** | **0.932** | **0.061** | **0.080** | **0.061** | **0.080** | **0.056** | **1.65E-10** | **0.056** | **1.65E-10** | **0.005** | **0.080** | **0.005** | **0.080** |

**Table S11: Best fit models assuming the prevalence of hybridization in nature is equal to that observed in our data.** Included are models that received >5% Akaike weights for their respective analyses. For each dataset, the best fit model is bold. Maximum-likelihood parameter estimates for speciation, extinction, and net diversification are reported. Non-hybridizing is abbreviated as NH, Hybridizing as H; A and B indicate the two hidden states.

| Definition of Hybridization | Dataset | Model | Akaike Weight | Speciation | | | | Extinction | | | | Net Diversification | | | |
|---|---|---|---|---|---|---|---|---|---|---|---|---|---|---|---|
| | | | | *NH-A* | *NH-B* | *H-A* | *H-B* | *NH-A* | *NH-B* | *H-A* | *H-B* | *NH-A* | *NH-B* | *H-A* | *H-B* |
| **Narrow** | **All Species** | **HiSSE: No H-B, all transitions** | **0.862** | **0.018** | **0.093** | **0.146** | **NA** | **0.006** | **6.16E-4** | **0.043** | **NA** | **0.018** | **0.092** | **0.103** | **NA** |
| | | HiSSE: No H-B, no double transitions | 0.125 | 0.028 | 0.110 | 0.161 | NA | 0.029 | 0.001 | 6.23E-2 | NA | 0.028 | 1.09E-1 | 0.099 | NA |
| | **Sympatric Species** | **HiSSE: No H-B, no double transitions** | **0.991** | **0.267** | **0.095** | **0.128** | **NA** | **0.032** | **0.000** | **0.023** | **NA** | **0.267** | **0.095** | **0.105** | **NA** |
| | **Plethodontids** | **HiSSE: No H-B, no double transitions** | **0.915** | **0.035** | **0.082** | **0.156** | **NA** | **7.28E-11** | **1.69E-10** | **3.22E-10** | **NA** | **0.035** | **0.082** | **0.156** | **NA** |
| | **Non-plethodontids** | **HiSSE: No double transitions** | **0.612** | **0.009** | **0.051** | **0.146** | **0.234** | **0.028** | **0.049** | **0.087** | **0.043** | **0.009** | **0.002** | **0.059** | **0.191** |
| | | HiSSE: No H-B, no double transitions | 0.298 | 0.097 | 0.012 | 0.098 | NA | 2.00E-10 | 0.003 | 0.001 | NA | 0.097 | 0.009 | 0.097 | NA |
| **Broad** | **All Species** | **HiSSE: No double transitions** | **0.993** | **0.022** | **0.089** | **0.116** | **0.171** | **0.066** | **0.012** | **0.004** | **0.002** | **0.022** | **0.077** | **0.112** | **0.169** |
| | **Sympatric Species** | **HiSSE: All parameters free** | **0.999** | **0.020** | **0.090** | **0.160** | **0.102** | **0.061** | **0.001** | **0.005** | **0.003** | **0.020** | **0.088** | **0.155** | **0.099** |
| | **Plethodontids** | **HiSSE: No H-B, no double transitions** | **0.979** | **0.033** | **0.080** | **0.150** | **NA** | **0.000** | **3.09E-10** | **1.65E-10** | **NA** | **0.033** | **0.080** | **0.150** | **NA** |
| | **Non-plethodontids** | **CID-2: Three transitions, no double** | **0.958** | **0.047** | **0.080** | **0.047** | **0.080** | **0.041** | **1.65E-10** | **0.041** | **1.65E-10** | **0.047** | **0.080** | **0.006** | **0.080** |

**Table S12: Best fit models assuming 30% of species hybridize.** Included are models that received >5% Akaike weights for their respective analyses. For each dataset, the best fit model is bold. Maximum-likelihood parameter estimates for speciation, extinction, and net diversification are reported. Non-hybridizing is abbreviated as NH, Hybridizing as H; A and B indicate the two hidden states.

| Tree | Frequency of Hybridization | Dataset | Speciation λ | | Extinction μ | | Net Diversification $r = \lambda - \mu$ | | Extinction Fraction $\varepsilon = \mu \div \lambda$ | | Turnover $\tau = \lambda + \mu$ | |
|---|---|---|---|---|---|---|---|---|---|---|---|---|
| | | | H | NH | H | NH | H | NH | H | NH | H | NH |
| All Species | Fully-sampled (11% N, 13% B) | N | **0.101 ± 5e-04** | 0.0588 ± 1.928e-03 | 4.47e-10 ± 1.262e-11 | **0.035 ± 7.68e-04** | **0.101 ± 5e-04** | 0.0237 ± 2.68e-03 | 4.45e-09 ± 1.314e-10 | **0.703 ± 0.0352** | 0.101 ± 5e-04 | 0.0938 ± 1.198e-03 |
| | | B | **0.103 ± 2.22e-03** | 0.0572 ± 2.02e-03 | 1.38e-04 ± 8.08e-06 | **0.033 ± 5.52e-04** | **0.103 ± 2.22e-03** | 0.0242 ± 2.56e-03 | 1.34e-03 ± 5.82e-05 | **0.677 ± 0.032** | 0.103 ± 2.24e-03 | 0.0903 ± 1.494e-03 |
| | Emprical | N | **0.111 ± 1.82e-3** | 0.0549 ± 1.718e-3 | 2.9e-3 ± 4.14e-4 | **0.0335 ± 8.08e-4** | **0.108 ± 1.832e-3** | 0.0214 ± 2.52e-3 | 0.0262 ± 3.74e-3 | **0.715 ± 0.036** | **0.114 ± 1.898e-3** | 0.0884 ± 9.18e-4 |
| | | B | **0.114 ± 3.46e-3** | 0.0517 ± 1.97e-3 | 5.73e-4 ± 2.52e-5 | **0.022 ± 7.36e-4** | **0.113 ± 3.46e-3** | 0.0297 ± 2.7e-3 | 5.19e-3 ± 2.82e-4 | **0.541 ± 0.0346** | **0.114 ± 3.46e-3** | 0.0737 ± 1.242e-3 |
| | 20% | N | **0.124 ± 5.9e-4** | 0.0552 ± 1.696e-3 | 9.39e-3 ± 8.62e-5 | **0.0383 ± 8.36e-4** | **0.115 ± 5.04e-4** | 0.017 ± 2.54e-3 | 0.0755 ± 3.2e-4 | **0.794 ± 0.0356** | **0.134 ± 6.76e-4** | 0.0935 ± 8.6e-4 |
| | | B | **0.117 ± 2.72e-3** | 0.0528 ± 2.04e-3 | 2.39e-3 ± 1.55e-4 | **0.0273 ± 7.24e-4** | **0.115 ± 2.88e-3** | 0.0255 ± 2.74e-3 | 0.0214 ± 1.95e-3 | **0.647 ± 0.039** | **0.12 ± 2.58e-3** | 0.0801 ± 1.368e-3 |
| | 30% | N | **0.148 ± 9.74e-5** | 0.0486 ± 1.696e-3 | **0.0453 ± 5.76e-5** | 6.38e-3 ± 1.85e-4 | **0.102 ± 3.98e-5** | 0.0422 ± 1.878e-3 | 0.307 ± 1.874e-4 | 0.164 ± 0.01032 | **0.193 ± 1.55e-4** | 0.055 ± 1.512e-3 |
| | | B | **0.127 ± 3.36e-3** | 0.046 ± 1.73e-3 | 3.64e-3 ± 9.66e-5 | **0.0463 ± 1.374e-3** | **0.124 ± 3.46e-3** | -3.1e-4 ± 3.1e-3 | 0.0293 ± 1.298e-3 | **1.29 ± 0.0838** | **0.131 ± 3.26e-3** | 0.0923 ± 3.56e-4 |
| Sympatric Species | Fully-sampled (11% N, 13% B) | N | **0.0944 ± 2.96e-03** | 0.0486 ± 1.632e-03 | 6.65e-10 ± 1.856e-11 | **0.0367 ± 8.96e-04** | **0.0944 ± 2.96e-03** | 0.0119 ± 2.52e-03 | 7.08e-09 ± 1.684e-10 | **0.855 ± 0.043** | 0.0944 ± 2.96e-03 | 0.0853 ± 7.46e-04 |
| | | B | **0.0983 ± 6.12e-03** | 0.0445 ± 1.89e-03 | 3.61e-03 ± 7.06e-04 | **0.0413 ± 1.164e-03** | **0.0947 ± 6.74e-03** | 3.22e-03 ± 3.06e-03 | 0.052 ± 0.01222 | **1.12 ± 0.069** | 0.102 ± 5.5e-03 | 0.0859 ± 7.26e-04 |
| | Emprical | N | **0.115 ± 9.82e-3** | 0.0444 ± 1.892e-3 | 3.19e-5 ± 2.06e-11 | **0.0105 ± 2.76e-4** | **0.115 ± 9.82e-3** | 0.034 ± 2.16e-3 | 2.94e-4 ± 1.088e-5 | **0.293 ± 0.01934** | **0.115 ± 9.82e-3** | 0.0549 ± 1.616e-3 |
| | | B | **0.113 ± 6.08e-3** | 0.045 ± 1.898e-3 | 5.03e-3 ± 4.02e-4 | **0.0146 ± 6.82e-4** | **0.108 ± 6.48e-3** | 0.0304 ± 2.58e-3 | 0.0501 ± 4.6e-3 | **0.441 ± 0.0402** | **0.118 ± 5.68e-3** | 0.0596 ± 1.216e-3 |
| | 20% | N | **0.111 ± 8.14e-3** | 0.0419 ± 1.728e-3 | 8.57e-4 ± 8.68e-5 | **9.45e-3 ± 3.5e-4** | **0.11 ± 8.04e-3** | 0.0324 ± 2.08e-3 | 7.54e-3 ± 1.53e-4 | **0.283 ± 0.0202** | **0.112 ± 8.22e-3** | 0.0513 ± 1.378e-3 |
| | | B | **0.106 ± 4.6e-3** | 0.047 ± 1.854e-3 | 3.02e-3 ± 1.644e-4 | **0.0273 ± 1.102e-3** | **0.103 ± 4.74e-3** | 0.0197 ± 2.96e-3 | 0.0306 ± 2.04e-3 | **0.732 ± 0.0568** | **0.109 ± 4.46e-3** | 0.0744 ± 7.58e-4 |
| | 30% | N | **0.128 ± 6.38e-5** | 0.0407 ± 1.588e-3 | 0.0231 ± 4.62e-7 | 0.0255 ± 7.4e-4 | **0.105 ± 6.42e-5** | 0.0153 ± 2.32e-3 | 0.18 ± 9.28e-5 | **0.736 ± 0.0406** | **0.151 ± 6.34e-5** | 0.0662 ± 8.46e-4 |
| | | B | **0.114 ± 3.24e-3** | 0.0382 ± 1.694e-3 | 3.53e-3 ± 1.108e-4 | **0.0457 ± 1.462e-3** | **0.11 ± 3.14e-3** | -7.46e-3 ± 3.16e-3 | 0.0311 ± 7.14e-5 | **1.48 ± 0.0972** | **0.117 ± 3.36e-3** | 0.0839 ± 2.32e-4 |
| Plethodontids | Fully-sampled (11% N, 13% B) | N | 0.0687 ± 3.5e-04 | 0.0671 ± 2.56e-03 | 4.27e-04 ± 5.48e-05 | 2.7e-04 ± 1.002e-04 | 0.0682 ± 4.06e-04 | 0.0668 ± 2.56e-03 | 6.27e-03 ± 8.22e-04 | 4.31e-03 ± 3.8e-04 | 0.0691 ± 2.96e-04 | 0.0674 ± 2.54e-03 |
| | | B | **0.108 ± 2.44e-04** | 0.0573 ± 1.342e-03 | 4.57e-04 ± 5.02e-05 | 3.28e-03 ± 2.78e-04 | **0.108 ± 2.94e-04** | 0.054 ± 1.586e-03 | 4.23e-03 ± 4.7e-04 | **0.0679 ± 8.32e-03** | **0.109 ± 1.948e-04** | 0.0606 ± 1.112e-03 |
| | Emprical | N | **0.137 ± 2.72e-5** | 0.0521 ± 1.524e-3 | 2.31e-6 ± 4.24e-7 | **1.93e-3 ± 1.002e-4** | **0.137 ± 2.76e-5** | 0.0502 ± 1.62e-3 | 1.69e-5 ± 3.1e-6 | **0.0426 ± 3.08e-3** | **0.137 ± 2.7e-5** | 0.0541 ± 1.43e-3 |
| | | B | **0.131 ± 2.3e-3** | 0.0484 ± 2.14e-3 | 3.13e-3 ± 5.38e-4 | **0.0245 ± 1.448e-3** | **0.128 ± 1.77e-3** | 0.0239 ± 3.6e-3 | 0.0231 ± 3.72e-3 | **0.727 ± 0.0854** | **0.134 ± 2.84e-3** | 0.0729 ± 6.94e-4 |
| | 20% | N | **0.0979 ± 5.5e-6** | 0.0413 ± 1.478e-3 | 4.26e-4 ± 1.204e-8 | **0.0137 ± 7.66e-4** | **0.0975 ± 5.48e-6** | 0.0276 ± 2.24e-3 | 4.35e-3 ± 1.63e-7 | **0.435 ± 0.0512** | **0.0984 ± 5.5e-6** | 0.055 ± 7.12e-4 |
| | | B | **0.143 ± 1.788e-4** | 0.0491 ± 1.638e-3 | 3.48e-6 ± 1.334e-6 | **1.4e-3 ± 6.66e-5** | **0.143 ± 1.776e-4** | 0.0477 ± 1.704e-3 | 2.42e-5 ± 9.24e-6 | **0.0332 ± 2.26e-3** | **0.143 ± 1.802e-4** | 0.0505 ± 1.572e-3 |
| | 30% | N | **0.159 ± 2.08e-3** | 0.0495 ± 1.532e-3 | 6.33e-3 ± 1.784e-3 | 2.23e-3 ± 3.3e-4 | **0.152 ± 3.12e-4** | 0.0473 ± 1.474e-3 | 0.0383 ± 0.01032 | 0.0442 ± 6.28e-3 | **0.165 ± 3.86e-3** | 0.0518 ± 1.654e-3 |
| | | B | **0.15 ± 1.58e-4** | 0.0448 ± 1.586e-3 | 2.28e-5 ± 7.84e-6 | **7.13e-8 ± 8.16e-9** | **0.15 ± 1.502e-4** | 0.0448 ± 1.586e-3 | **1.51e-4 ± 5.18e-5** | 1.86e-6 ± 2.5e-7 | **0.15 ± 1.658e-4** | 0.0448 ± 1.586e-3 |
| Non-Plethodontids | Fully-sampled (11% N, 13% B) | N | 0.0704 ± 3.22e-04 | 0.0641 ± 3.28e-03 | 9.92e-04 ± 1.004e-05 | **0.0642 ± 2.36e-03** | **0.0694 ± 3.32e-04** | -1.17e-04 ± 5.64e-03 | 0.0141 ± 2.06e-04 | **1.15 ± 0.0902** | 0.0714 ± 3.12e-04 | **0.128 ± 9.4e-04** |
| | | B | 0.0703 ± 4.58e-03 | 0.0721 ± 3.34e-03 | **6.34e-03 ± 4.54e-04** | 3.58e-03 ± 1.286e-04 | 0.064 ± 4.76e-03 | 0.0685 ± 3.46e-03 | 0.0997 ± 0.0162 | 0.0614 ± 9e-03 | 0.0767 ± 4.44e-03 | 0.0757 ± 3.24e-03 |
| | Emprical | N | 0.0815 ± 3.72e-3 | 0.0626 ± 3.42e-3 | 1.16e-3 ± 2.48e-5 | **7.88e-3 ± 1.07e-4** | 0.0803 ± 3.72e-3 | 0.0547 ± 3.34e-3 | 0.0144 ± 5.86e-4 | **0.145 ± 0.01188** | 0.0826 ± 3.74e-3 | 0.0705 ± 3.5e-3 |
| | | B | 0.0759 ± 2.12e-3 | 0.076 ± 1.174e-3 | 0.0115 ± 5.62e-3 | 9.68e-3 ± 3.04e-3 | 0.0644 ± 7.74e-3 | 0.0663 ± 4.2e-3 | 0.177 ± 0.0902 | 0.15 ± 0.0492 | 0.0874 ± 3.5e-3 | 0.0857 ± 1.878e-3 |
| | 20% | N | **0.0781 ± 3.38e-3** | 0.0412 ± 1.734e-3 | 1.89e-3 ± 4.92e-5 | **0.039 ± 7.88e-4** | **0.0762 ± 3.42e-3** | 2.2e-3 ± 2.5e-3 | 0.0245 ± 1.188e-3 | **1.04 ± 0.0662** | 8e-2 ± 3.32e-3 | 0.0803 ± 9.8e-4 |
| | | B | 0.0708 ± 5.42e-3 | 0.0719 ± 2.88e-3 | 1.44e-4 ± 1.038e-5 | 1.66e-4 ± 3.72e-7 | 0.0706 ± 5.42e-3 | 0.0717 ± 2.88e-3 | 2.38e-3 ± 4.8e-4 | 2.6e-3 ± 2.52e-4 | 0.0709 ± 5.42e-3 | 0.0721 ± 2.88e-3 |
| | 30% | N | **0.133 ± 2.46e-3** | 0.0458 ± 1.806e-3 | **0.0538 ± 1.12e-3** | **0.029 ± 6.06e-4** | **0.0789 ± 3.58e-3** | 0.0169 ± 1.418e-3 | 0.407 ± 0.01384 | **0.656 ± 0.0212** | **0.186 ± 1.35e-3** | 0.0748 ± 2.3e-3 |
| | | B | 0.0741 ± 3.52e-3 | 0.0735 ± 1.922e-3 | 8.98e-3 ± 4.44e-3 | 8.45e-3 ± 2.4e-3 | 0.0651 ± 7.98e-3 | 0.0651 ± 4.32e-3 | 0.163 ± 0.0864 | 0.153 ± 0.049 | 0.0831 ± 9.24e-4 | 0.082 ± 4.88e-4 |

**Table S13: All model averaged parameter estimates.** Results are reported for each diversification rate parameter quantified, for each tree, each dataset, and each assumed frequency of hybridization. Bold values indicate parameter estimates that differ significantly among character states, with the boldened values indicating the larger rate estimate. N and B correspond to the narrow and broad datasets respectively.

| Tree | Frequency of Hybridization | Dataset | Probability NH > H | | | | |
|---|---|---|---|---|---|---|---|
| | | | Speciation | Extinction | Net Diversification | Extinction Fraction | Turnover |
| All Species | Fully-sampled (11% N, 13% B) | N | 0.026 | 1.000 | 0.005 | 1.000 | 0.291 |
| | | B | 0.032 | 1.000 | 0.006 | 1.000 | 0.235 |
| | Emprical | N | 0.004 | 1.000 | 0.002 | 1.000 | 0.028 |
| | | B | 0.012 | 1.000 | 0.003 | 1.000 | 0.035 |
| | 20% | N | 0.000 | 1.000 | 0.000 | 1.000 | 0.000 |
| | | B | 0.004 | 1.000 | 0.000 | 1.000 | 0.022 |
| | 30% | N | 0.000 | 0.000 | 0.000 | 0.115 | 0.000 |
| | | B | 0.000 | 1.000 | 0.000 | 1.000 | 0.000 |
| Sympatric Species | Fully-sampled (11% N, 13% B) | N | 0.012 | 1.000 | 0.000 | 1.000 | 0.241 |
| | | B | 0.045 | 1.000 | 0.011 | 0.996 | 0.352 |
| | Emprical | N | 0.000 | 1.000 | 0.000 | 1.000 | 0.000 |
| | | B | 0.000 | 1.000 | 0.000 | 1.000 | 0.000 |
| | 20% | N | 0.000 | 0.999 | 0.000 | 1.000 | 0.000 |
| | | B | 0.000 | 1.000 | 0.000 | 1.000 | 0.003 |
| | 30% | N | 0.000 | 0.664 | 0.000 | 0.976 | 0.000 |
| | | B | 0.000 | 1.000 | 0.000 | 1.000 | 0.000 |
| Plethodontids | Fully-sampled (11% N, 13% B) | N | 0.165 | 0.279 | 0.166 | 0.296 | 0.164 |
| | | B | 0.000 | 0.949 | 0.000 | 0.975 | 0.000 |
| | Emprical | N | 0.000 | 1.000 | 0.000 | 1.000 | 0.000 |
| | | B | 0.000 | 0.981 | 0.000 | 0.996 | 0.000 |
| | 20% | N | 0.000 | 1.000 | 0.000 | 1.000 | 0.000 |
| | | B | 0.000 | 1.000 | 0.000 | 1.000 | 0.000 |
| | 30% | N | 0.000 | 0.181 | 0.000 | 0.604 | 0.000 |
| | | B | 0.000 | 0.000 | 0.000 | 0.000 | 0.000 |
| Non-Plethodontids | Fully-sampled (11% N, 13% B) | N | 0.329 | 1.000 | 0.037 | 1.000 | 1.000 |
| | | B | 0.583 | 0.038 | 0.652 | 0.150 | 0.508 |
| | Emprical | N | 0.211 | 1.000 | 0.100 | 1.000 | 0.324 |
| | | B | 0.423 | 0.455 | 0.505 | 0.457 | 0.309 |
| | 20% | N | 0.000 | 1.000 | 0.000 | 1.000 | 0.600 |
| | | B | 0.510 | 0.600 | 0.507 | 0.611 | 0.511 |
| | 30% | N | 0.000 | 0.000 | 0.000 | 1.000 | 0.000 |
| | | B | 0.251 | 0.725 | 0.258 | 0.728 | 0.168 |

**Table S14: Empirical P-values corresponding to Supplementary Table S2.** Results are reported for each diversification rate parameter quantified, for each tree, each dataset, and each assumed frequency of hybridization. Empirical P-values are obtained by calculating all possible ratios between non-hybridizing (NH) and hybridizing species' (H) model-averaged parameter estimates and calculating the proportion of comparisons in which the value for the non-hybridizing lineage is greater than that of hybridizing lineage. Thus, a value of 0 means in every comparison H > NH, and vice versa. N and B correspond to the narrow and broad datasets respectively.

| **Model** | **Trait-dependent Diversification** | *Best Model* | *ΔAICc < 4* | *Akaike Weights > 0.05* |
|---|---|---|---|---|
| BiSSE Null: One transition | No | 0 | 0 | 0 |
| BiSSE Null: Two transitions | No | 0 | 0 | 0 |
| CID-2: All transitions, all equal | No | 0 | 0 | 0 |
| CID-2: Three transitions, no double | No | 0 | 0 | 0 |
| CID-2: No double transitions, all equal | No | 0 | 0 | 0 |
| CID-4: Three transitions | No | 0 | 0 | 0 |
| CID-4: Equal transitions | No | 0 | 0 | 0 |
| BiSSE: All parameters free | Yes | 0 | 0 | 0 |
| HiSSE: No NH-B, all transitions | Yes | 0 | 0 | 0 |
| HiSSE: No NH-B, no double transitions | Yes | 0 | 0 | 0 |
| HiSSE: No H-B, all transitions | Yes | 6 | 12 | 16 |
| HiSSE: No H-B, no double transitions | Yes | 21 | 50 | 53 |
| HiSSE: All parameters free | Yes | **52** | **77** | **83** |
| HiSSE: No double transitions | Yes | 21 | 36 | 38 |

**Table S15: Model support for analysis of 100 phylogenies retrieved from the posterior distribution of Jetz & Pyron (2018).** Values represent the number of times each model was supported according to the given condition. This set of analyses was conducted using the tree containing all species, assuming 20% of species hybridize.

| Definition | Dataset | Number contrasts | P-value |
|---|---|---|---|
| Narrow | All Species | 15 | 0.0386 |
| | Sympatric | 15 | 0.0156 |
| | Plethodontids | 9 | 0.2513 |
| | Non-plethodontids | 6 | 0.0166 |
| Broad | All Species | 15 | 0.0411 |
| | Sympatric | 18 | 0.0032 |
| | Plethodontids | 9 | 0.0735 |
| | Non-plethodontids | 8 | 0.0047 |

**Table S16: Results of sister clade comparisons (Barraclough, Harvey and Nee, 1996) using each of the four datasets and two definitions of hybridization.** Only sister clade comparisons with three or more taxa were considered. P-values correspond to the alternative hypothesis that species richness of clades that hybridize is greater than those that do not.

| Model | Trait-dependent? | **Narrow - No Glutinosus Group** | | | **Broad - No Glutinosus Group** | | |
|---|---|---|---|---|---|---|---|
| | | AICc | Delta-AICc | Akaike Weights | AICc | Delta-AICc | Akaike Weights |
| BiSSE Null: One transition | No | 3970.47 | 144.44 | > 0.001 | 3970.55 | 140.81 | 0.000 |
| BiSSE Null: Two transitions | No | 3927.01 | 100.99 | > 0.001 | 3927.20 | 97.46 | 0.000 |
| CID-2: All transitions, all equal | No | 3913.05 | 87.02 | > 0.001 | 3914.49 | 84.75 | 0.000 |
| CID-2: Three transitions, no double | No | 3861.05 | 35.03 | > 0.001 | 3876.95 | 47.21 | 0.000 |
| CID-2: No double transitions, all equal | No | 3902.45 | 76.42 | > 0.001 | 3903.93 | 74.19 | 0.000 |
| CID-4: Three transitions | No | 3883.22 | 57.20 | > 0.001 | 3879.62 | 49.88 | 0.000 |
| CID-4: Equal transitions | No | 3901.09 | 75.07 | > 0.001 | 3928.09 | 98.34 | 0.000 |
| BiSSE: All parameters free | Yes | 3884.57 | 58.54 | > 0.001 | 3885.26 | 55.52 | 0.000 |
| HiSSE: No NH-B, no double transtions | Yes | 3862.00 | 35.97 | > 0.001 | 3863.32 | 33.58 | 0.000 |
| HiSSE: No NH-B, all transitions | Yes | 3913.07 | 87.05 | > 0.001 | 3885.62 | 55.88 | 0.000 |
| HiSSE: No H-B, no double transitions | Yes | 3890.56 | 64.54 | > 0.001 | 3844.78 | 15.03 | 0.001 |
| HiSSE: No H-B, all transitions | Yes | 3845.58 | 19.56 | > 0.001 | 3846.27 | 16.53 | 0.000 |
| HiSSE: All parameters free | Yes | 3834.01 | 7.99 | 0.018 | 3835.50 | 5.76 | 0.053 |
| **HiSSE: No double transitions** | **Yes** | **3826.02** | **0.00** | **0.982** | **3829.74** | **0.00** | **0.946** |

**Table S17. The 14 models fitted in our analysis excluding the *Glutinosus* group within genus *Plethodon* using both the narrow and broad definition of hybridization**. Models varied in number of hidden states included as well as in the number of unique transitions among states permitted. HiSSE models contain hidden states, whereas BiSSE models do not. Double transitions are those in which two-state jumps occur (i.e. NH-**A** to H-B or vice versa). CID stands for character independent, thus CID-2 and CID-4 represent the two novel null models of trait-independent diversification that may be implemented in the HiSSE framework. Rows in bold indicate best-fit models.
